# Chaperonin recognition of protein dynamics drives drug resistance

**DOI:** 10.64898/2026.06.03.729952

**Authors:** Junlang Liu, Zhihui Qi, João V. Rodrigues, Zhouyu Zhang, Qianxiang Xu, Abhinav Dubey, Harrison K. Wang, Dongtao Cui, Zeyang Li, Fatima Aly, Doeke R. Hekstra, Eugene I. Shakhnovich

## Abstract

The emergence of drug resistance is typically driven by mutations that alter drug-target affinity, yet the role of host cellular machinery regulating these processes remains unclear. Here, we reveal that the chaperonin GroEL/S promotes drug resistance through recognition of protein dynamics. Using directed evolution of *E. coli* DHFR under antibiotic stress and varying GroEL/S expression, we identify a well-folded resistance variant whose fitness, despite tight inhibitor binding, is critically potentiated by GroEL/S engagement. X-ray crystallography, NMR, molecular dynamics and kinetic modeling reveal that millisecond-timescale flipping of the M20 loop generates steric accessibility and a kinetic window for chaperonin interaction that forcibly displaces tightly bound inhibitors, thereby overriding thermodynamic equilibrium of inhibitor binding to restore the active enzyme pool and preserve metabolic flux. Our findings not only reveal a novel paradigm of “dynamic recognition” where both conformational kinetics and distribution govern chaperonin recognition but also establish chaperonins as “deligandases” that actively modulate *in vivo* drug binding, suggesting that chaperone surveillance of the cellular proteome extends beyond quality control to govern native protein function. This mechanism defines a previously unrecognized route for the rapid development of drug resistance, with implications for understanding therapeutics of microbial infections and human malignancies.

## Introduction

The rapid emergence of drug resistance poses a critical challenge to modern medicine^1,2^. While resistance is typically driven by mutations that directly alter drug-target affinity^3,4^, the expansion of these adaptive phenotypes depends critically on host cellular machineries, particularly molecular chaperones^5^. As cells explore the vast sequence space, chaperones sustain proteome homeostasis by ensuring proper protein folding, assisting nascent chain transport, and preventing lethal aggregation^6–10^. However, the precise mechanism by which chaperones shape evolution remains a subject of intense debate^11^. Classically, chaperones are viewed as evolutionary buffers that mask the deleterious effects of destabilizing mutations, allowing cryptic genetic variation to accumulate silently within a population as a genetic reservoir for future adaptation to severe environmental stress^12,13^. Beyond this passive buffering, chaperones can also act as active potentiators, facilitating specific genetic variants that unlock entirely novel phenotypes^14^. Resolving how chaperones dictate these evolutionary outcomes is critical for understanding how proteins rapidly adapt to evade targeted therapeutics.

A major bottleneck in uncovering how chaperones drive drug resistance is our incomplete understanding of the chaperone recognition mechanism. While chaperone such as GroEL/S is known to possess a well-characterized client library *in vivo*^15,16^, *in vitro* studies demonstrate its remarkable capacity to interact with and refold more than half of the denatured *E. coli* proteome^17,18^. This broad promiscuity indicates that the fundamental molecular prerequisites for chaperonin binding, such as hydrophobic motifs or amphiphilic peptides^19–21^, are not exclusive to a small subset of proteins, but are instead ubiquitous across the proteome^22^. Although this observation explains how chaperonins buffer severe folding defects, it fails to explain how they could distinguish their client proteins and act as active potentiators for certain mutations.

Consequently, true *in vivo* recognition cannot be dictated by static features, such as sequence and native fold, alone. Rather, productive recognition must be governed by the combination of this molecular basis and the temporal or kinetic feasibility of the interaction once chaperonin and client are in spatial proximity to chaperonins. Preliminary results on model proteins such as GFP suggest that chaperonin dependence is somewhat correlated with folding kinetics^23,24^. In this framework, GroEL/S maintains constant surveillance over the proteome as a dynamic sensor, engaging substrates based on the timescale of hydrophobic region exposure. Therefore, the *in vivo* client library is not a fixed repertoire, but a highly fluid demographic that shifts dynamically with the cellular state or mutations of a protein. This dynamic recognition hypothesis provides a compelling physical basis for evolutionary potentiation, where mutations that subtly alter the exposure timescale of recognition motifs, even in well-folded proteins, could seamlessly co-opt this chaperonin surveillance network, exploiting temporal windows to enable novel functional traits, such as drug resistance.

To experimentally validate this dynamic recognition hypothesis and resolve molecular-level mechanisms for how chaperonins drive drug resistance, we used GroEL/S and *E. coli* Dihydrofolate Reductase (DHFR) as our model system. Since DHFR is an essential enzyme targeted by the antibiotic trimethoprim (TMP), its evolutionary landscape can be defined by a strict biophysical trade-off between mutations that confer resistance and compromised conformational integrity^25–28^. By subjecting DHFR to continuous directed evolution under escalating TMP stress across varying GroEL/S expression levels, we observed that GroEL/S overexpression broadened the sequence space available to evolution by buffering DHFR overexpression and destabilizing mutations. Most crucially, we also identified a unique point mutation that acquired robust antibiotic resistance without compromising its global folding stability and binding affinity to TMP. By integrating genetic, biophysical, structural, *in vitro* biochemical, and simulation approaches, we demonstrate that GroEL/S recognizes this well-folded mutant by capturing its highly specific conformational dynamics. Upon recognition, GroEL/S physically displaces the tightly bound antibiotic through its unfolding-refolding process. This discovery explicitly establishes dynamics as a fundamental determinant of GroEL/S recognition, providing definitive molecular evidence for a new mechanism by which GroEL/S acts as a “deligandase” that regulates *in vivo* drug binding affinity and therefore serves as an active potentiator of evolutionary innovation.

### GroEL/S overexpression promotes the emergence of distinct DHFR mutations

We performed continuous directed evolution of DHFR by subjecting *E. coli* strains with varying GroEL/S expression levels to selection under the antibiotic TMP (Fig. 1a). Throughout the evolution, cell growth rates were monitored in real-time to dynamically adjust TMP selection pressure, ensuring that each evolutionary lineage was sustained under continuous selection. After 200–300 generations of evolution within 60-80 passages, terminal mutations at the *folA* locus were identified via Sanger sequencing (Supplementary Fig. 1-2, and Supplementary Table 1).

**Fig. 1.**
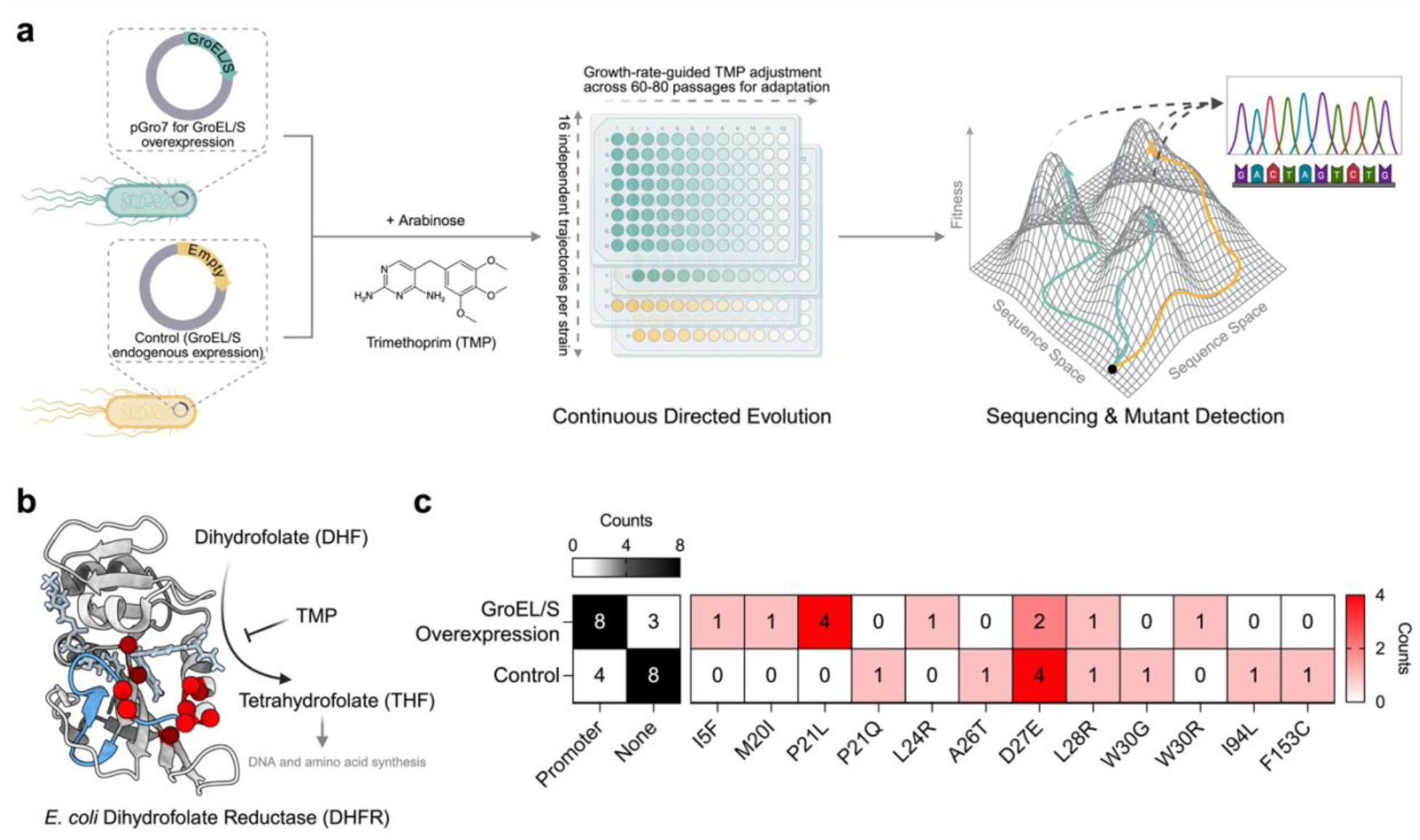
GroEL/S availability shapes the accessible fitness landscape of DHFR. **a**, Schematic of the continuous directed evolution platform. *E. coli* strains harboring either a GroEL/S overexpression plasmid (pGro7) or an empty control vector were subjected to TMP pressure over 45-60 days. Sixteen independent trajectories per strain were monitored in real-time, with TMP concentrations dynamically adjusted for adaptation and terminal mutations identified via deep sequencing. **b**, Structural representation of *E. coli* DHFR (PDB: 7FQ7) highlighting the active site and the M20 loop (blue). Red spheres indicate the distribution of mutations identified in this study. The chemical reaction catalyzed by DHFR (DHF to THF) and its inhibition by TMP are shown. **c**, Mutational landscape of evolved DHFR variants under different GroEL/S expression levels. The left heatmap displays the counts of trajectories with promoter mutations and with no substitutions in both promoter region and DHFR sequence (None). The right heatmap quantifies the occurrence of all identified amino acid substitutions in the DHFR coding region across 16 independent trajectories per strain.

While control lineages with endogenous GroEL/S levels exhibited constrained mutational exploration at the *folA* (DHFR) locus, GroEL/S overexpression promoted the accumulation of both regulatory and coding mutations (Fig. 1c, left). Specifically, strains with GroEL/S overexpression exhibited a significantly higher frequency of promoter mutations (Fig. 1c, left), which consistently drove DHFR overexpression (Supplementary Fig. 3). These results indicate that GroEL/S promotes evolvability by both enhancing proteostasis capacity, facilitating protein overexpression, and broadening the accessible sequence space, thereby enabling the exploration of mutational trajectories that would otherwise be evolutionarily inaccessible.

Analysis of the specific amino acid substitutions revealed a bifurcated evolutionary response (Fig. 1c, right). While all identified mutations across both conditions were localized to the TMP binding pocket (Fig. 1b), underscoring a common structural basis for antibiotic resistance, the specific mutational outcomes were highly dependent on GroEL/S expression levels. Specifically, beyond two shared mutational hotspots observed in both groups (i.e., D27 and L28), each condition yielded a different set of substitutions. To elucidate the molecular mechanisms underlying these divergent evolutionary outcomes, we characterized the biophysical properties of the emerging variants across two dimensions: (i) the extent of molten globule state, given GroEL/S is known to recognize molten globule proteins and buffer destabilizing yet adaptive mutations^12,13^ (Fig. 2a-b, and Supplementary Fig. 4); and (ii) TMP resistance quantified by inhibition constants (*K*_i_), since all mutations were evolved under sustained TMP pressure (Fig. 2b).

**Fig. 2.**
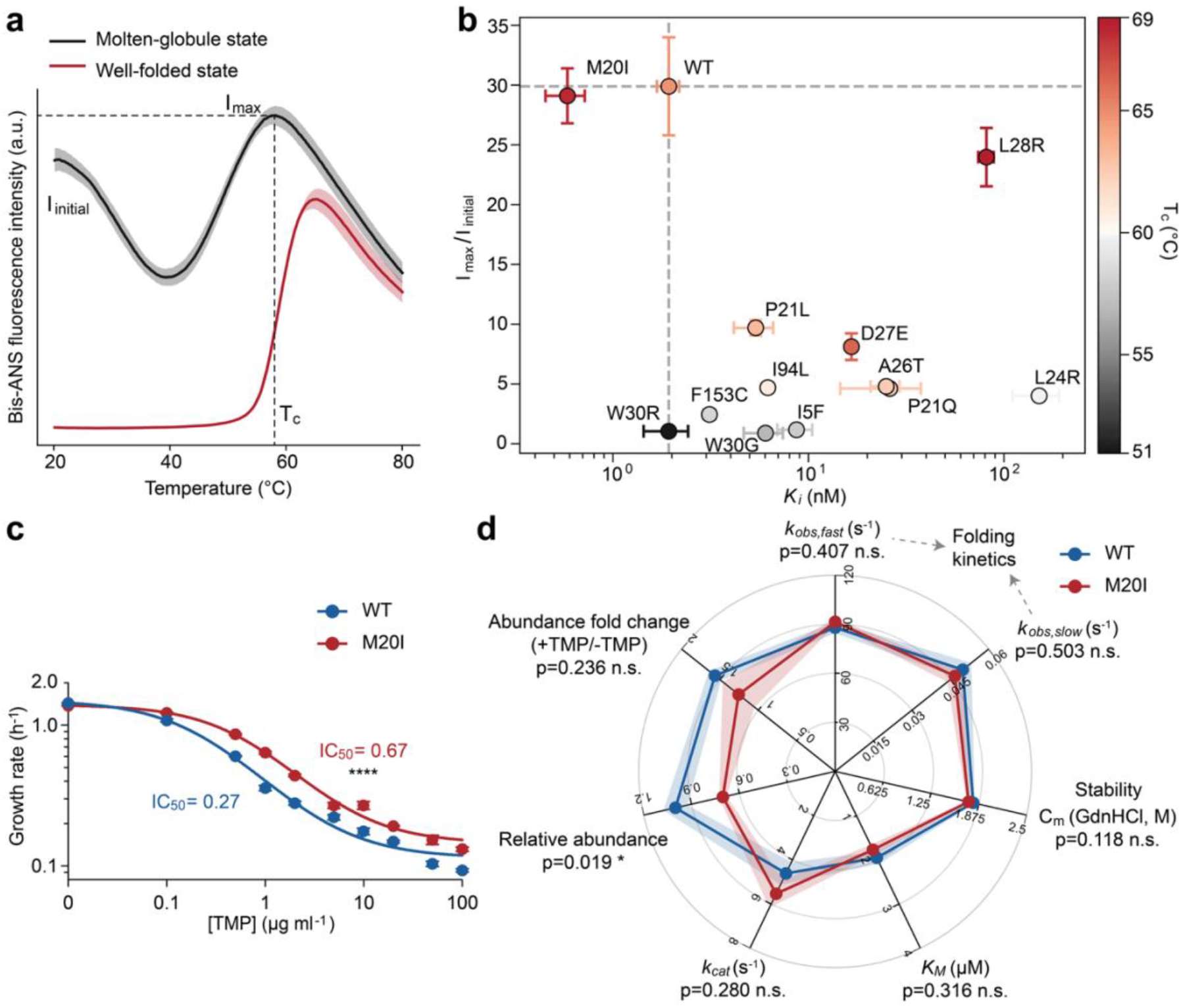
M20I emerges as a distinct mutation emerging under GroEL/S overexpression. **a**, Schematic of the bis-ANS fluorescence assay utilized to probe the extent of molten globule state and stability. Molten-globule-like proteins (black) show high fluorescence intensity both at room temperature (i.e., I_initial_) and at the critical temperature T_c_ (i.e., I_max_), while well-folded proteins show a sharp increase in fluorescence only upon reaching T_c_ (red). We used the intensity ratio I_max_/I_initial_ as a readout for molten globule states of DHFR variants. **b**, Two-dimensional biophysical properties (I_max_/I_initial_ and *K*_i_) screening of the resistance variants identified the M20I as a distinct outlier. Data points are color-coded by T_c_. **c**, Growth assays in M9 medium demonstrating that the M20I mutation (red), introduced into a clean *E. coli* BW25113 genetic background via CRISPR-Cas9, is sufficient to confer significant TMP resistance compared to the WT (blue). **d**, Systematic evaluation of biochemical and biophysical properties of M20I (red) and WT (blue). No single factor tested adequately accounts for the observed TMP-resistance of M20I, suggesting that M20I operates via a non-canonical antibiotic resistance mechanism.

The extent of molten globule state was quantified via bis-ANS fluorescence, which probes hydrophobic core accessibility and exhibits increased intensity upon binding^29^. While well-folded proteins exhibit low basal fluorescence signal at room temperature, molten-globule-like variants display high fluorescence due to flexible tertiary structures (Fig. 2a). The protein structure expands and destabilizes as temperature rises, eventually reaching a critical temperature (T_c_) where global unfolding occurs. Thus, T_c_ serves as a proxy for thermodynamic stability. Consequently, we defined the ratio of peak intensity (I_max_) at T_c_ to initial intensity (I_initial_) at room temperature as a sensitive metric of molten globule extent, where a lower ratio signifies more pronounced pre-existing hydrophobic exposure and less stability (Supplementary Fig. 5). While most evolved mutations clustered within an expected regime, where gains in resistance were coupled with decreased stability and tertiary structure integrity, M20I emerged as a significant outlier (Fig. 2b). Specifically, M20I exhibited a unique combination of a smaller TMP inhibition constant, *K*_i_, than WT (Supplementary Fig. 6)^27^, a markedly high fluorescence ratio similar to WT, and a higher T_c_ than WT when binding to NADPH. These results indicate it as a well-folded protein with strong TMP binding affinity.

We further confirmed the *in vivo* TMP resistance of the M20I variant by introducing this single point mutation on the WT genetic background (Fig. 2c, and Supplementary Fig. 7). Strikingly, the M20I strain exhibited no basal fitness cost, growing as robustly as WT in the absence of drug (Supplementary Fig. 8), yet achieved a 2.5-fold increase in *in vivo* IC_50_, enabling significantly enhanced growth under stringent TMP concentrations (Supplementary Fig. 9). To elucidate the underlying mechanism of resistance, we performed a comprehensive characterization of its biophysical and biochemical properties, including protein folding kinetics, thermodynamic stability, enzymatic activity, and *in vivo* protein abundance (Fig. 2d, and Supplementary Fig. 10-13). No significant differences were observed between M20I and WT across these parameters, with the notable counter-intuitive exception that M20I exhibited lower basal abundance *in vivo* than WT and showed a diminished increase in protein levels under equivalent TMP selection pressure (at their respective IC50 concentrations). This striking discrepancy between reduced cellular availability and robust resistance levels is difficult to reconcile through traditional antibiotic resistance models^25^, indicating that TMP resistance of M20I emerges via a non-canonical mechanism.

### GroEL/S is required for M20I antibiotic resistance

Given that the M20I mutation emerged from an evolutionary lineage under GroEL/S overexpression (Fig. 1c, right), we investigated whether its resistance phenotype is strictly dependent on GroEL/S. Indeed, overexpressing GroEL/S further enhanced the *in vivo* IC_50_ of M20I strain by 2-fold, whereas WT tolerance remained entirely unaffected (Supplementary Fig. 14). Conversely, GroEL/S knockdown completely abrogated M20I antibiotic resistance, reducing it to WT levels (Fig. 3a). This loss of resistance was further evidenced by the collapse of the M20I growth rate curve under GroEL/S knockdown, rendering its dose-response curve indistinguishable from that of the WT across a range of TMP concentrations (Fig. 3b, and Supplementary Fig. 15). As controls under the same knockdown conditions, the molten-globule-like W30C mutant completely fails to grow at a moderate concentration of TMP, whereas the GroEL/S-independent L28R strain remains much more resistant than the WT. This stark contrast confirms that the loss of M20I resistance upon GroEL/S knockdown is an intrinsic consequence of the mutation, rather than a pleiotropic artifact of global cellular stress induced by chaperonin depletion. This result also highlights a distinct mechanistic divergence between resistant strains. While W30C strictly depends on GroEL/S to maintain basic structural stability, M20I uniquely co-opts GroEL/S to restore its function *in vivo* in the presence of TMP. Consistent with this functional dependence, *in vivo* pull-down assays confirmed a robust physical interaction, revealing that M20I DHFR recruits significantly more GroEL under TMP stress compared to WT (Fig. 3c). As a control, W30C exhibited low abundance alongside high GroEL interaction, a hallmark of typical molten-globule-like behavior. The increased band intensity of GroEL after TMP exposure for the M20I strain may reflect the selective enrichment of populations with effective GroEL/S engagement under drug pressure. Whereas for the WT strain, a much slighter increase in band intensity may instead be mainly driven by a modest, TMP-induced upregulation of endogenous GroEL/S expression (Supplementary Fig. 16), although the underlying mechanism remains to be elucidated. To further verify that this enhanced recruitment is driven by direct physical affinity rather than indirect cellular factors, we performed complementary *in vitro* validation using size-exclusion chromatography (SEC), which similarly demonstrated a significantly stronger direct interaction between purified M20I and GroEL compared to WT (Supplementary Fig. 17). It is also worth noting that M20I confers a consistent 2.5-fold higher IC50 than WT in both M9 (Fig. 2c) and LB (Fig. 3a) media, further validating the robustness of this resistance phenotype across different metabolic conditions.

**Fig. 3.**
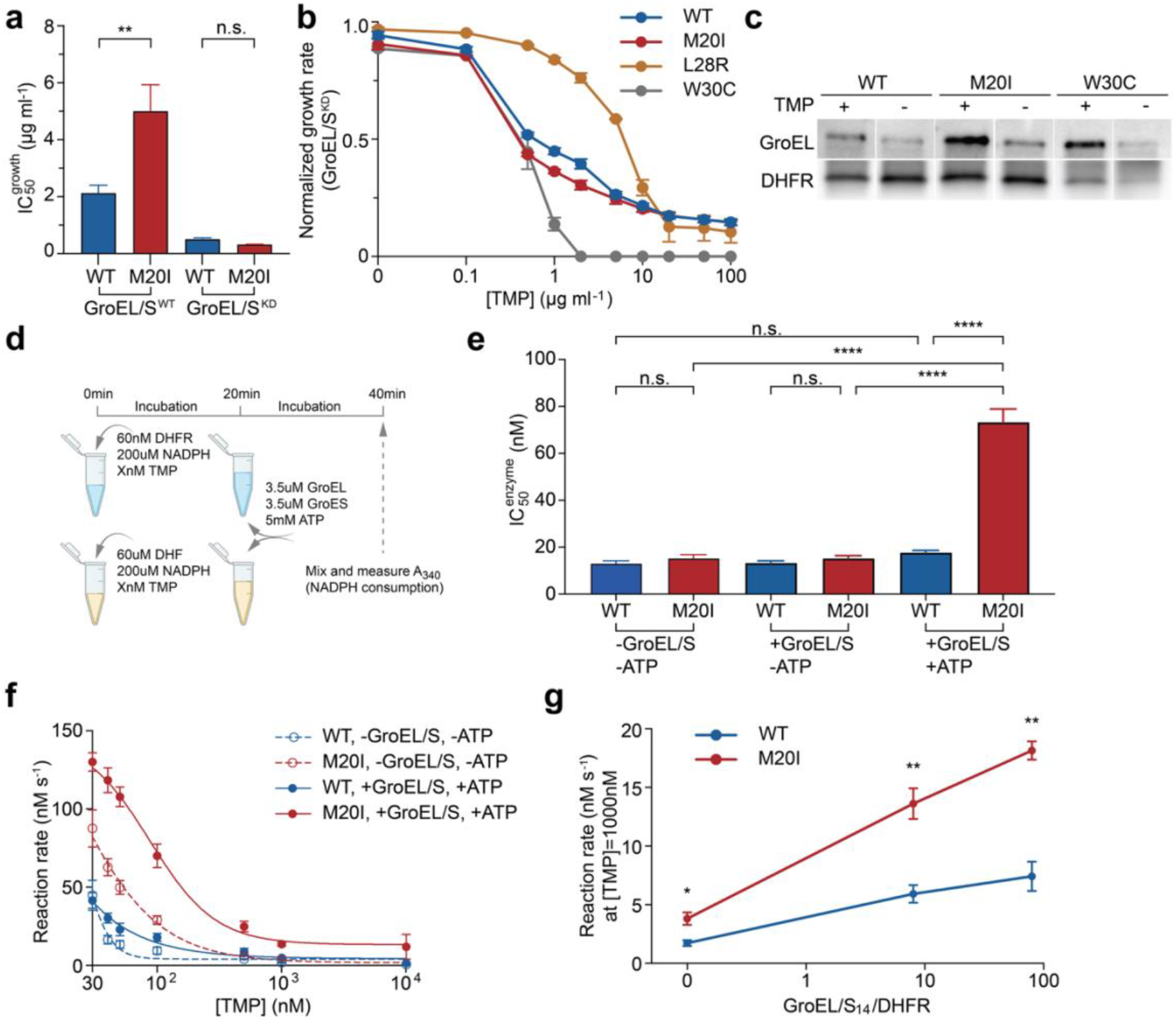
*In vivo* and *in vitro* validation of GroEL/S-dependent M20I antibiotic resistance. **a**, *In vivo* cell growth IC_50_ measurements show that M20I antibiotic resistance to TMP is reduced to WT level upon GroEL/S knockdown (KD). Luria-Bertani (LB) medium was utilized for all GroEL/S knockdown assays and their respective controls to ensure robust cell growth under GroEL/S knockdown conditions. **b**, Normalized growth rates for WT, M20I, L28R, and W30C strains across TMP concentrations after GroEL/S knockdown confirm that resistance loss is specific to M20I and not a result of general GroEL/S knockdown-induced toxicity. **c**, *In vivo* pull-down analysis of GroEL using DHFR as bait across various strains and TMP conditions. **d**, Schematic of the *in vitro* reconstructed system designed to evaluate the GroEL/S-dependent recovery of M20I enzymatic activity. **e**, *In vitro* recovery of enzymatic activity is specific to the M20I mutant and strictly requires both GroEL/S and ATP. **f**, M20I exhibited sustained enzymatic activity even at TMP concentrations three orders of magnitude above the baseline IC50 in the presence of GroEL/S and ATP. See Supplementary Fig. 18 for full concentration range. **g**, M20I enzymatic activity at high TMP concentrations (e.g., 1000nM) is titratable and strictly dependent on GroEL availability, establishing a causal link between chaperone levels and the resistance phenotype.

To further establish the mechanistic causality between GroEL/S and M20I antibiotic resistance, we reconstructed the enzymatic system *in vitro*. Deviating from traditional chaperonin assays that utilize denatured substrates, we employed folded DHFR proteins pre-incubated with NADPH and TMP (Fig. 3d). This design allowed us to test if GroEL/S could rescue a physically inhibited, rather than misfolded, enzyme population. The addition of GroEL/S and ATP led to dramatic in vitro rescue of inhibition of M20I by TMP, whereas interaction with GroEL/S alone, without ATP, had no recovery effect (Fig. 3e, and Supplementary Fig. 18). By contrast, the WT variant exhibited no significant rescue under identical GroEL/S and ATP conditions. Strikingly, M20I maintained a strictly ATP-dependent residual activity even at TMP concentrations that are three orders of magnitude above its basal IC_50_ (Fig. 3f). Titrating the GroEL/S_14_:DHFR ratio in the range 8:1 to 80:1 yielded a dose-dependent increase in M20I reaction activity, confirming that this rescue relies directly on GroEL/S function (Fig. 3g). Collectively, these *in vitro* data reveal a novel, unexplored “deligandase” function of GroEL/S, where the ATP-driven refolding reaction enables active clearance of a bound inhibitor with sub-nM-level binding affinity^30^.

### GroEL recognition of M20I is determined by protein conformational dynamics

To identify the mechanistic basis for GroEL/S-mediated antibiotic resistance in M20I, we first compared the crystal structures of WT and M20I in complex with NADP+ and TMP^31,32^. Despite their divergent cellular phenotypes, the two variants exhibit nearly identical ground state native conformations, with no significant deviations in the protein backbone or ligand-binding poses (Fig. 4a). This structural identity further underscored that M20I is well-folded and its recognition by GroEL/S is not triggered by misfolded states as traditionally thought. Instead, NMR chemical shift differences between WT and M20I revealed that the M20I mutation significantly alters the local chemical environments within the M20, F-G, and G-H loops in its ternary complex bound to NADPH and TMP (Fig. 4b, and Supplementary Fig. 19-20), pointing toward a dynamic rather than static distinction. We characterized these altered dynamics using CPMG relaxation dispersion experiments^33,34^, which quantified the population of the high-energy minor state unique to M20I and its exchange rate with ground state (Fig. 4c, and Supplementary Fig. 21). Notably, residues exhibiting millisecond-scale dynamics are all localized in these three loops, recapitulating the pattern of weighted chemical shift changes (Fig. 4b) and further confirming the dynamic nature of M20I-induced difference. Global fitting of the dispersion curves for these residues revealed that M20I undergoes conformational switching with an exchange rate of 316 s^−1^, populating a minor state at approximately 3.8%. Molecular dynamics (MD) simulations further characterized this minor state as an ‘open’ conformation, revealing that the M20 loop frequently flips out of the active site in the M20I variant (Fig. 4c). Specifically, the difference in Root Mean Square Fluctuation (ΔRMSF) analysis showed significantly increased dynamics in the M20 loop region for the M20I open state (Fig. 4d). This aberrant behavior is fundamentally different from the equilibrium dynamics of the WT, where the M20 loop remains constrained by its interactions with NADPH and TMP, despite the transition between the standard closed and occluded states (Fig. 4d)^31,33,34^. To understand why the M20I mutation triggers the flip-out of M20 loop, we performed a dihedral angle analysis of all available DHFR crystal structures in the PDB (Supplementary Fig. 22). This meta-analysis reveals that the native M20 residue possesses significant conformational plasticity, allowing it to adapt to varying ligands, whereas an isoleucine at position 20 is locally more rigid. We hypothesized that this inability of I20 to adapt to smaller inhibitors like TMP results in a weaker steric constraint, which facilitates the conformational flexibility observed in our simulations and experiments.

**Fig. 4.**
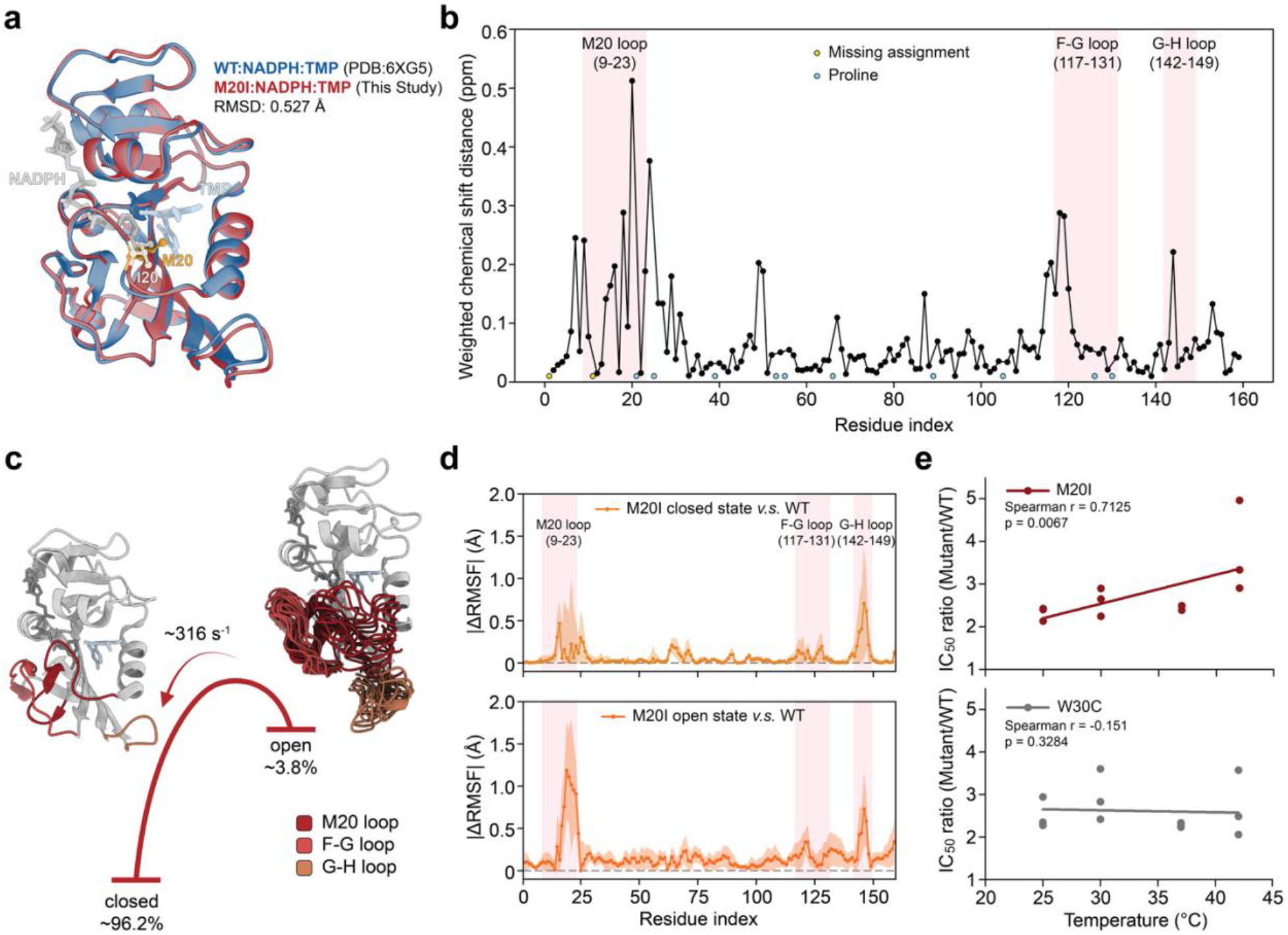
M20I alters DHFR conformational dynamics to enable GroEL/S recognition. **a**, Overlay of crystal structures of WT (PDB: 6XG5) and M20I (this study) in complex with NADPH and TMP reveals nearly identical ground-state conformations for both the protein backbone and ligand-binding poses. **b**, Weighted chemical shift distances between WT and M20I in complex with NADPH and TMP, highlighting significant changes in the chemical environments specifically within the M20 loop, F-G loop, and G-H loop regions. **c**, CPMG relaxation dispersion experiments identifying a two-state conformational exchange process within M20I, with the minor (open) state with M20 loop flipped out captured by molecular dynamics (MD) simulations. **d**, Difference in Root Mean Square Fluctuation (ΔRMSF) from MD simulations between the two M20I states and WT, demonstrating profoundly different equilibrium dynamics for M20I open state, particularly in the M20 loop region. **e**, *In vivo* growth assays show that elevated temperature specifically enhances the recognition of M20I by GroEL/S, unlike the molten-globule-like W30C, correlating increased protein dynamics with improved resistance phenotype.

Integrating X-ray crystallography, NMR, and MD simulations, we demonstrate that M20I exhibits significant structural plasticity even when fully inhibited by TMP, in contrast with the WT variant. The flip-out of the M20 loop populates a transient, high-energy ensemble that renders the loop sterically and temporally accessible for GroEL/S recognition. To demonstrate *in vivo* that conformational dynamics drive recognition, we used temperature as a biophysical probe of dynamic effects (Fig. 4e). Elevated temperatures specifically enhanced the GroEL/S-dependent resistance of M20I as compared to WT, whereas the molten-globule-like W30C control showed no such temperature dependence of enhancement.

### Kinetic model of GroEL-assisted rescue of DHFR inhibition

Together these experimental results point out to a novel mechanism by which GroEL recognizes native conformation ensembles featuring transiently open loops, engaging them in unfolding-refolding cycles to produce freshly folded active DHFR free of TMP inhibition. To quantitatively evaluate this active rescue mechanism, we developed an analytical kinetic model of the chaperonin-assisted pathway (Fig. 5a) based on five core premises: (i) M20I DHFR exists in distinct ‘open’ and ‘closed’ states defined by the M20 loop; (ii) these states interconvert dynamically, with exchange kinetics and equilibrium populations parameterized directly from our NMR CPMG data; (iii) TMP can bind both conformations; (iv) GroEL selectively recognizes the *open state* of M20I, driving it through an active unfolding-refolding cycle that removes the TMP inhibitor and releases fully active DHFR; and (v) WT DHFR remains entirely unrecognized by GroEL due to its lack of open state when bound to NADPH and TMP. Full mathematical derivations, simulation parameters, and their biophysical justifications for the numerical simulations are detailed in Methods and Supplementary Table 2.

**Fig. 5.**
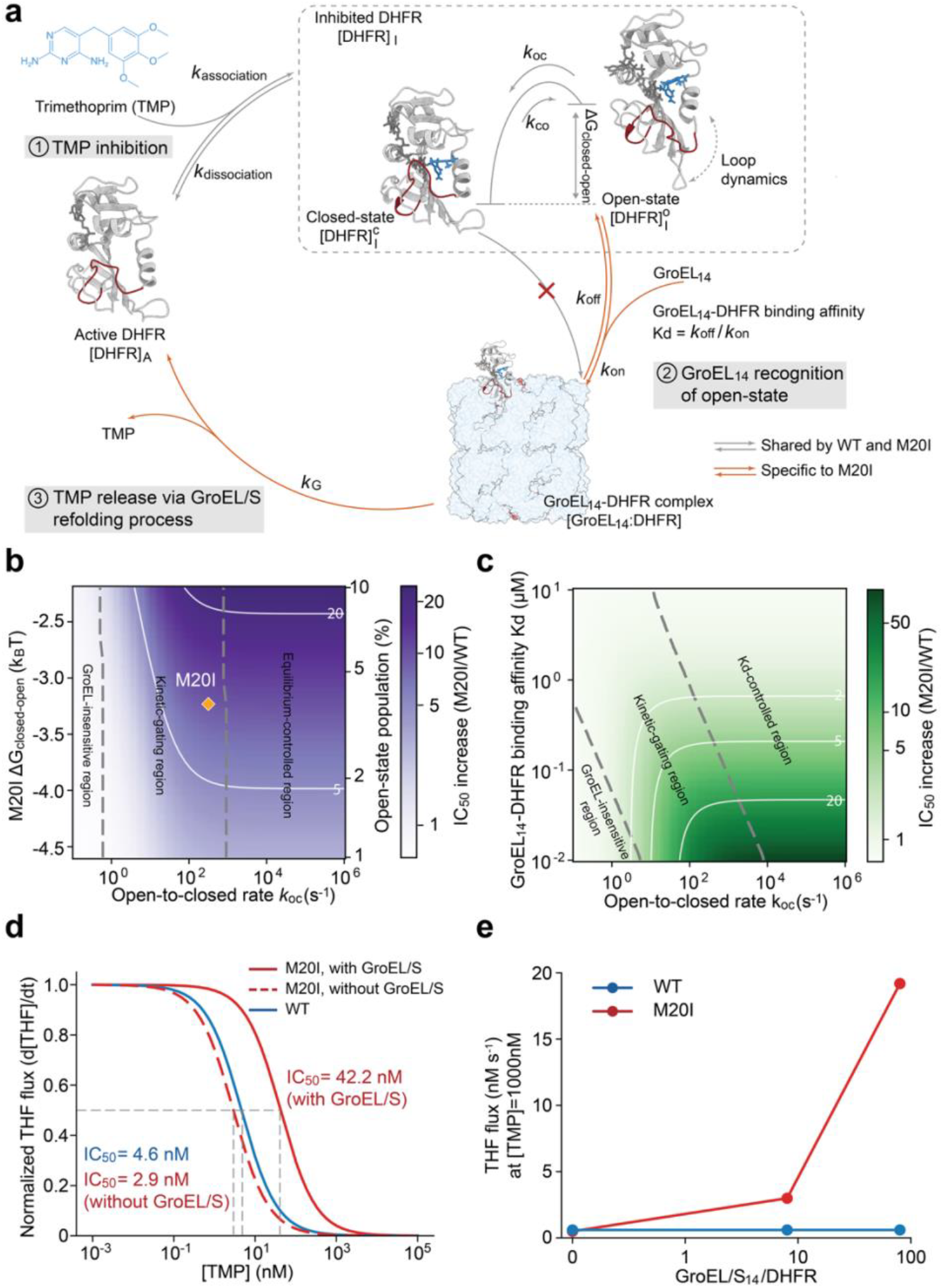
Kinetic model for GroEL/S-mediated M20I antibiotic resistance. **a**, Schematic of the kinetic model. TMP binding inhibits active DHFR. Within the inhibited ensemble, M20I exchanges between closed and open conformations with rate constants *k*_*co*_ and *k*_*co*_. GroEL_14_ selectively recognizes the open state and forms a GroEL_14_–DHFR complex. Productive refolding by GroEL/S regenerates active DHFR and promotes TMP release. Grey arrows indicate processes shared by WT and M20I, whereas orange arrows denote M20I-specific steps. **b**, Predicted IC_50_ increase of M20I relative to WT as a function of the open-to-closed rate constant *k*_*oc*_ and the free-energy difference Δ*G*_*closed*−*open*_. Three regimes emerge: a GroEL-insensitive region at low *k*_*oc*_, in which *k*_*oc*_ is sufficiently slow such that recognition of the open state does not substantially enhance flux; a kinetic-gating region, in which increasing *k*_*oc*_ markedly enhances the THF flux; and an equilibrium-controlled region, in which conformational exchange is fast enough that GroEL rescue is determined predominantly by equilibrium population of the open state rather than kinetics. The position of M20I is indicated, with a value determined from NMR CPMG results. **c**, Predicted IC_50_ increase of M20I relative to WT as a function of the GroEL_14_–DHFR binding affinity *K*_*d*_ and GroEL_14_ concentration. **d**, Predicted TMP dose-response curve of M20I and WT in presence or absence of GroEL/S rescue, recapitulating the experimental observation in Fig. 3e,f. **e**, Predicted catalytic flux at 1000 nM TMP, mirroring the GroEL/S availability-dependent recovery of M20I enzymatic activity shown in Fig. 3g.

To determine whether the chaperonin rescue effect is driven primarily by the conformational distribution or exchange kinetics, we first generated the phase diagrams mapping both THF metabolic flux, the primary determinant of cell growth under TMP stress, and the resulting IC_50_ increase, as a function of the thermodynamic equilibrium (Δ*G*_*closed*−*open*_) and transition rate constant (*k*_oc_) (Fig. 5b, and Supplementary Fig. 23). The landscape partitions into three distinct regimes based on *k*_oc_. At relatively low *k*_oc_ values, compared to the diffusion-limited GroEL encounter rate constant, infrequent exposure of the binding region reduces the probability of productive GroEL-DHFR interactions, resulting in a negligible IC_50_ or flux enhancement. In the millisecond-to-second regime, IC_50_ and the metabolic flux are kinetically controlled by the transition rate itself. Conversely, at rapid *k*_oc_ limits, GroEL encounters fast-exchanging conformational ensembles, where IC_50_ and the metabolic flux are determined by the equilibrium population of the recognizable open state rather than the kinetics of interconversion. Critically, the exchange parameters and substate populations experimentally derived from NMR CPMG measurements place M20I within the intermediate ‘kinetic-gating’ regime. This positioning suggests that for the M20I variant, GroEL recognition of the open state is predominantly governed by the kinetics of interconversion between open and closed states rather than the equilibrium population of the open state.

As GroEL_14_ complex-substrate binding affinity *K*_D_ values are technically challenging to measure and span orders of magnitude (Supplementary Table 3), we further explored this landscape by mapping the predicted IC_50_ increase, as well as THF flux, across a broad range of chaperonin-substrate binding affinities (*K*_D_) (Fig. 5c, and Supplementary Fig. 23), which allows us to test the boundaries of different mechanisms. We found that for the specific *k*_oc_ of M20I, GroEL recognition is strictly kinetically gated across a wide range of the *K*_D_ spectrum. It is only when binding affinity drops to the micromolar level that the mechanism crosses the phase boundary, transitioning into a regime governed primarily by *K*_D_ and the equilibrium distribution of the open state, where chaperonin-mediated rescue also becomes negligible.

It is noteworthy that both numerical simulations utilized a TMP concentration of 89 µM, reflecting intracellular TMP levels in *E. coli* MG1655 cells challenged with 5 µM external TMP^28^. Despite this *in silico* TMP concentration being 2-3 orders of magnitude higher than both *in vivo* and *in vitro* IC_50_, M20I maintains the IC_50_, as well as THF metabolic flux, a full order of magnitude greater than that of the WT, faithfully recapitulating both the sustained *in vivo* cell growth of the M20I variant at high concentration of TMP (Fig. 2c, and Supplementary Fig. 9) and substantial residual enzymatic activity in *in vitro* assays (Fig. 3f-g) of M20I under saturating TMP concentrations

Furthermore, our kinetic model quantitatively predicts IC_50_ of M20I under GroEL/S rescue (Fig. 5d), as well as the dependence of M20I enzymatic activity recovery on GroEL/S availability (Fig. 5e, and Supplementary Fig. 24), providing a robust quantitative framework bridging fundamental molecular dynamics and macroscopic cellular resistance.

### GroEL/S removes bound drugs as a conserved evolutionary strategy

To determine if the GroEL/S-mediated resistance observed in M20I reflects a broader evolutionary principle, we analyzed the natural distribution of amino acids aligned to *E. coli* reference position 20 across diverse DHFR sequences. Comparison of chromosomal *folA* sequences with clinically isolated, plasmid-encoded resistant *dfrA* variants revealed a striking enrichment of isoleucine in the resistant population (Fig. 6a). While I20 is present in only 9.9% of *folA* sequences, its prevalence increases nearly three-fold to 28.3% in *dfrA* isolates, significantly outcompeting the methionine (10.3%) in these high-TMP-stress clinical contexts. We validated this evolutionary preference by analyzing recently reported *in vivo* laboratory fitness of a library of DHFR homologs under varying TMP concentrations^35^. Across a diverse set of sequence backgrounds, homologs containing I20 exhibited consistently higher relative fitness at high TMP concentrations compared to those with M20 or L20 (Fig. 6b). Specifically, within this dataset, of the seven homologs evaluated with substitutions at position 20, the only two to show elevated fitness under TMP stress were those harboring the M-to-I mutation (Supplementary Fig. 25). This consistent performance across diverse homologs with low sequence identity suggests that the I20-driven resistance phenotype is not dependent on specific protein context but could be instead an intrinsic property of the residue’s effect on loop dynamics and chaperonin recognition.

**Fig. 6.**
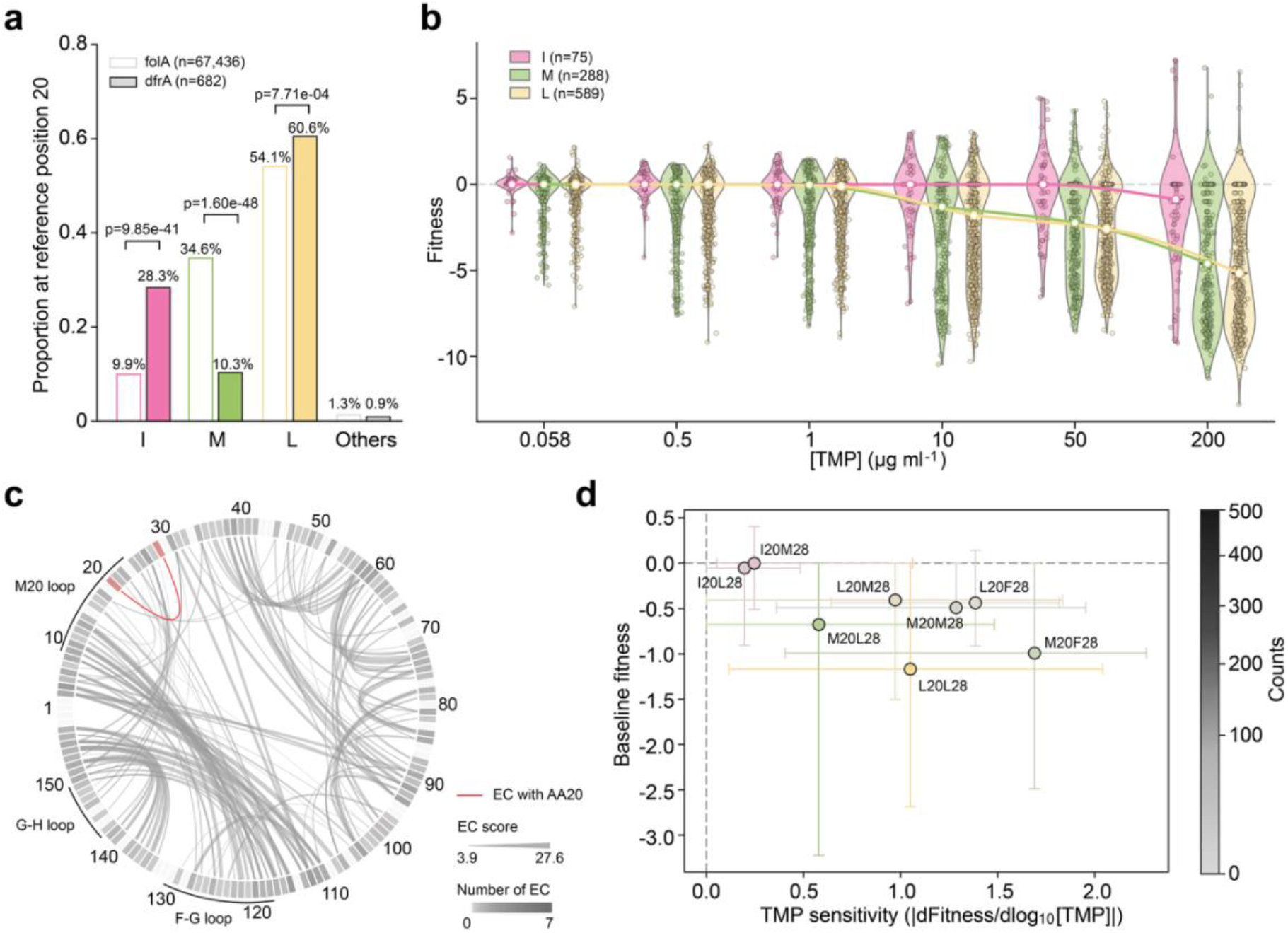
Evolutionary and epistasis analyses reveal the structural isolation of position 20 and the prevalence of I20 under TMP stress. **a**, Comparison of chromosomal (*folA*) and clinically isolated, plasmid-encoded resistant (*dfrA*) DHFR sequences reveals selection for isoleucine at position aligned to *E. coli* DHFR residue 20 (I20). **b**, Quantitative *in vivo* fitness measurements show that DHFR homologs with I20 exhibit consistently higher relative fitness than those with alternative amino acids at the 20th position under high TMP concentrations. **c**, Evolutionary coupling (EC) analysis identifies position 20 as an evolutionary and structurally isolated site, exhibiting weak coupling only with position 28. **d**, Epistasis analysis between positions 20 and 28 shows that the I20 resistance phenotype is independent of its co-evolved partner at position 28.

To further investigate the independence of this mechanism on the sequence background, we performed Evolutionary Coupling (EC) analysis^36,37^ to map the co-evolutionary constraints on position 20. We found that position 20 is remarkably isolated, exhibiting weak coupling only with position 28 (Fig. 6c). This lack of widespread epistatic constraints suggests that the M20 loop can evolve new dynamic behaviors, such as the “flip-out” open state, without disrupting the overall structural integrity or catalytic function of the DHFR scaffold. Utilizing this same evolutionary dataset, epistasis analysis between positions 20 and 28 confirmed that the I20 resistance phenotype is robust and independent of its primary co-evolutionary partner. Specifically, when mapping baseline fitness against TMP sensitivity, variants harboring I20 consistently cluster in a regime of high basal fitness and minimal drug sensitivity. In stark contrast, variants containing M20 or L20 exhibit profound TMP sensitivity and reduced baseline fitness, irrespective of the accompanying residue at position 28 (Fig. 6d and Supplementary Figs. 26–28).

## Discussion

One metaphor for the evolution of drug resistance is navigating a multidimensional strait between the Scylla and Charybdis of biophysical constraints on molecular properties of evolving proteins^38–40^. Most resistant mutations exert pleiotropic effects: while a substitution may successfully evade an antibiotic, it frequently compromises essential baseline traits like thermodynamic stability or aggregation propensity. The prevailing model posits that chaperones passively buffer these detrimental side effects, relaxing strict biophysical constraints and opening up sequence space for broader evolutionary exploration^12,13,41,42^. Nevertheless, this concept has rarely been validated in the context of real-life clinical challenges, and it remains entirely unclear how chaperone networks can transcend this buffering role to actively potentiate emerging functions, such as drug resistance, under acute therapeutic stress.

Here, we addressed these questions by investigating the role of the chaperonin GroEL/S in a well-established, clinically relevant model of drug resistance evolution: the essential *E. coli* enzyme DHFR under stress from the FDA-approved antibiotic TMP (Fig. 1, and Supplementary Fig. 1-3). Among all core chaperones, the chaperonin is uniquely ancient, existing since the last universal common ancestor (LUCA) dating back 3.6 billion years^5^. DHFR serves as a quintessential model for quantitatively linking protein biophysics to cellular fitness^25–27,43^. Together, they form an exceptionally representative and tractable system for interrogating chaperone-mediated evolution. Through parallel directed evolution experiments with varying GroEL/S expression level, we found that trajectories under GroEL/S overexpression were more likely to acquire upstream promoter mutations that increased DHFR abundance (Fig. 1c, and Supplementary Fig. 3). In these instances, GroEL/S functions potentially to mitigate the deleterious effects of DHFR overexpression^44^. Concurrently, most coding mutations isolated from these trajectories exhibited a pronounced propensity to adopt the partially unfolded molten-globule state, coupled with significantly reduced TMP binding affinity (Fig. 1c, and 2b). These observations strongly align with the established paradigm that GroEL/S relaxes stringent constraints on protein stability^13^, thereby promoting escape from TMP inhibition.

In stark contrast to these typical mutations, we observed a remarkable *E. coli* DHFR single mutation, M20I, which did not fit the prevailing paradigm. Paradoxically, M20I exhibits enhanced molecular affinity for the antibiotic TMP as well as robust structural integrity (Fig. 2a-b, 2d, and Supplementary Fig. 4-6, 10-13), yet it drives significant TMP resistance *in vivo* (Fig. 2c, and Supplementary Fig. 7-9). By integrating genetic knockdown, pull-down assays, and *in vitro* reconstitution, we established GroEL/S as the definitive causal agent of this anomalous resistance (Fig. 3, Supplementary Fig. 14-18). Crucially, our multi-modal structural analyses combining X-ray crystallography, NMR, and MD simulations reveal that chaperonin recognition is dictated not by global unfolding, but by the unique, localized conformational dynamics of the intact M20I enzyme (Fig. 4, Supplementary Fig. 19-22). Specifically, upon recognizing the transiently flipped-out M20 loop, GroEL/S initiates an active unfolding-refolding cycle that results in the release of refolded TMP-free DHFR into cytoplasm, i.e. physically dislodging the bound antibiotic TMP and effectively acting as a “deligandase”. This clearance process directly enhances the concentration of functional inhibitor-free enzyme, ultimately conferring antibiotic resistance. Accordingly, we developed a detailed kinetic model to dissect the relative contributions of conformational exchange rate, equilibrium distribution, and chaperonin-substrate binding affinity to the observed antibiotic resistance (Fig. 5, Supplementary Fig. 23-24). Our analysis suggests that the chaperonin-mediated rescue of M20I is primarily driven by the kinetics of interconversion between the closed and open states, rather than the populations of these conformations. Supported by bioinformatic evidence showing the prevalence of I20-mediated resistance across natural and laboratory landscapes, we propose that chaperone exploitation of local dynamics represents a broadly conserved evolutionary mechanism (Fig. 6, Supplementary Fig. 25-28).

The emergence of the M20I variant under GroEL/S overexpression showcases an unprecedented strategy for circumventing classic biophysical trade-off between protein stability and antibiotic binding affinity^2–4^. Rather than operating as an isolated target mutation that must balance this trade-off, M20I successfully harnesses host cellular machinery to safeguard its function under drug stress. By specifically tuning its local conformational landscape to populate a transient substate recognizable by GroEL, this variant recruits the chaperonin to actively drive drug dissociation. The clinical and evolutionary relevance of this novel antibiotic resistance mechanism is further underscored by our bioinformatic analysis. Rather than being a peculiarity from laboratory directed evolution, I20 mutation is a common resistant variant significantly enriched among clinically isolated, plasmid-encoded resistant *dfrA* variants. This extensive natural conservation, combined with the site’s epistatic independence, suggests that the exploitation of localized loop dynamics to harness endogenous proteostasis networks represents a fundamental strategy for achieving high-level resistance while preserving overall structural integrity.

Beyond its immediate implications for drug resistance evolution, the M20I-GroEL interaction redefines the foundational principles of chaperonin substrate recognition by shifting the focus from static structural states to a unified dynamic framework. Traditionally, GroEL is thought to target misfolded intermediates or molten globules that expose hydrophobic surfaces. We propose that the mechanisms underlying traditional substrates and the dynamic variants identified here are not mutually exclusive but rather represent different positions along a continuous conformational dynamics spectrum. From this perspective, chaperonin engagement is governed by two decisive factors: steric accessibility (the exposure of the recognition motif) and temporal persistence (the duration and frequency of that exposure). Under this unified model, the constant surveillance by chaperone networks extends beyond misfolded substrates to encompass the dynamic fluctuations of the folded proteome, meaning both states are continually monitored but handled differently based on their kinetic properties. Traditional misfolded or molten-globule states represent proteins that remain in a recognizable state for an effectively infinite duration. In contrast, well-folded enzymes like M20I only become sterically accessible for GroEL interaction during transient, millisecond-timescale flipping of recognizable motifs.

The combination of steric and temporal accessibility is the primary filter for chaperonin selection *in vivo*. This framework also helps explain the discrepancy between *in vitro* refolding assays, in which ~850 proteins out of ~1000 detectable proteins are refolded by GroEL/S^18^, and *in vivo* proteomic studies, which identify only 85 proteins as core clients^15^. *In vitro* refolding assays may present an altered kinetic landscape, enabling GroEL/S to engage transient states that would otherwise be sterically or temporally inaccessible within the crowded cellular milieu. In contrast, traditional *in vivo* interactome profiling is generally unable to capture transient yet potentially functional interactions with GroEL and thus biased toward highly expressed proteins that exhibit pronounced intrinsic folding defects and remain in misfolded states long enough to be detected.

Taken together, these findings reveal the new “deligandase” function of GroEL/S beyond protein refolding: through an active unfolding–refolding cycle, it can reconfigure ligand-bound protein states and thereby regulate enzyme availability. Owing to its distinctive barrel-shaped architecture and promiscuous client interactions, GroEL/S may be uniquely positioned within the cell to physically dissociate high-affinity small molecules from their protein targets, thus promoting drug resistance. Mirroring our observations here that GroEL/S overrides the thermodynamic equilibrium of the drug–protein complex by forcibly removing bound TMP, such ATP-driven deliganding mechanism may universally enable cells to preserve essential metabolic flux under chemical stress, irrespective of intrinsic drug-binding affinities. These findings position GroEL/S not merely as a folding machine, but as a specialized proteostasis factor capable of actively remodeling the protein–ligand interactions to promote survival in the presence of potent inhibitors. Identifying additional examples of such regulation will be a critical next step, particularly for understanding chaperone-enabled drug resistance in more complex systems. This question is especially pertinent in cancer, where human chaperonin homologues, such as TRiC/CCT and Hsp60, are frequently overexpressed and may similarly contribute to therapeutic resistance by actively sequestering or clearing drug-bound states.

## Methods

All the reagent, resources, plasmids, primers, and cell strains used in this study are listed in Supplementary Table 5-8.

### Laboratory directed evolution

Continuous laboratory directed evolution was conducted using a Tecan Freedom EVO liquid handling platform, with cells cultured at 37 °C in a LiCONiC StoreX automated incubator and cell density continuously monitored via a Tecan Infinite M200 PRO plate reader. Two *E. coli* strains were constructed in the BW25113 background: an experimental strain harboring the pGro7 plasmid for arabinose-inducible GroEL/S overexpression, and a control strain transformed with an empty vector (pEmpty) generated by removing the GroEL/S genes from pGro7. Cells were cultured in standard M9 minimal medium (1× M9 salts, 0.2% glucose, 2 mM MgSO_4_, 0.1 mM CaCl_2_) supplemented with 0.01% arabinose, starting at an initial optical density (OD_600_) of 0.01. The average growth rate of uninhibited cultures was defined as the reference growth rate. Trimethoprim (TMP) selection pressure was introduced on Day 2 at an initial concentration of 0.16 µg ml^−1^. Cell density was recorded at 15-minute intervals throughout the experiment, and growth rates were calculated as described previously^45^. Once the plate-averaged OD_600_ reached 0.35, the cultures were automatically diluted to an OD_600_ of 0.025. To sustain continuous growth while maintaining stringent selective pressure, the TMP concentration was dynamically adjusted: it was increased twofold if the culture growth rate recovered to >75% of the reference rate, and reduced twofold if the growth rate dropped to <5% of the reference rate. After 60-80 passages of evolution, the final adapted strains were harvested and submitted to Genewiz (Azenta Life Sciences) for Sanger sequencing of the chromosomal *folA* locus. A total of sixteen independent evolutionary trajectories were maintained for each strain.

### Protein production

#### Non-isotope-labeled His-tagged *E. coli* dihydrofolate reductase (DHFR)

Plasmids encoding wild-type (WT) *E. coli* DHFR (UniProt: P0ABQ4) and 14 single-point mutants (I5F, M20I, P21L, P21Q, L24R, A26T, D27E, L28R, W30G, W30R, I94L, F153C, M20L, and W30C) on the pET-28a(+) vector with a C-terminal 6×His-tag were synthesized by GenScript Biotech (Nanjing, China) and subsequently transformed into *E. coli* BL21(DE3) cells. Cells harboring these DHFR plasmids were inoculated from frozen cell stocks into Terrific Broth (TB) medium and grown overnight (12-16 h) at 37 °C with shaking at 250 rpm. The overnight cultures were then used to inoculate fresh production medium at a 1% (v/v) ratio. This production medium consisted of TB supplemented with final concentrations of 0.4% (v/v) glycerol, 4 g L^−1^ glucose, 25 mM MOPS (pH 7.2), 0.4 mM citric acid, 40 µl L^−1^ of 1% (w/v) ferric ammonium citrate, and 50 µg ml^−1^ kanamycin. The production cultures were grown at 37 °C and 250 rpm until starting the mid-logarithmic phase (OD600 ≈ 0.6, typically ~3.5 h), at which point protein overexpression was induced via adding a final concentration of 100 µM isopropyl β-D-1-thiogalactopyranoside (IPTG). Following induction, the incubation temperature was reduced to 18 °C for 16 hours, and the cells were subsequently harvested by centrifugation (6,000 × *g*, 4 °C).

Harvested cell pellets were resuspended in Lysis Buffer A (100 mM bicine, 250 mM NaCl, 5 mM imidazole, pH 9.0) at a ratio of 1.5 ml g^−1^, supplemented with 1 mg ml^−1^ lysozyme, Complete Inhibitor Cocktail, and 0.01% (v/v) nuclease. The cell suspension was sonicated on wet ice for 1-2 minute per gram of pellet (1 s on, 2 s off, amplitude 20). The resulting lysate was centrifuged at 30,000 × *g* for 25 minutes to pellet the insoluble material, and the supernatant was extracted. The clarified supernatant was applied to a gravity-flow column packed with cOmplete™ His-Tag Purification Resin to isolate the DHFR protein. The resin was washed with Buffer A (20 mM Tris, 250 mM NaCl, 5 mM imidazole, pH 8.5) to remove non-specifically bound host proteins, and the recombinant DHFR was subsequently eluted with Buffer B (20 mM Tris, 250 mM NaCl, 50 mM imidazole, pH 8.0). Eluate fractions were evaluated using a NanoDrop spectrophotometer (Thermo Fisher Scientific), and those exhibiting an A_280_/A_260_ ratio > 1.7 were pooled. The pooled protein was loaded onto a HiLoad 16/600 Superdex 75 pg column for final purification via size exclusion chromatography (SEC) and buffer-exchange into Tris-buffered saline (20 mM Tris, 250 mM NaCl, pH 7.0). Corresponding elution fractions were concentrated to a final concentration of 3-30 mg ml^−1^ using a Vivaspin^®^ ultrafiltration unit (Sartorius). Protein purity and identity were verified by SDS-PAGE (Bio-Rad) and mass spectrometry (Bruker Impact II q-TOF), respectively, before the samples were aliquoted and stored at −70 °C.

The M20I mutant often co-purifies with endogenous NADPH, resulting in a reduced A_280_/A_260_ absorbance ratio. Due to energy transfer between W22 and the bound NADPH, the holo-complex exhibits decreased fluorescence emission at 340 nm and an emergent peak at 450 nm upon excitation at 295 nm. To isolate the apo-M20I form, anion exchange chromatography (IEX) was employed. Briefly, a HiPrep™ Q Fast Flow 16/10 column (20 ml column volume) was equilibrated with 3 column volumes (CVs) of IEX Buffer A (100 mM NaCl, 50 mM Tris, 1 mM TCEP, pH 7.5). Following sample loading, the column was washed with 20 CVs of IEX Buffer A at a flow rate of 1-2 ml min^−1^. Elution was performed using a 20-CV linear gradient from 0% to 100% IEX Buffer B (400 mM NaCl, 50 mM Tris, 1 mM TCEP, pH 7.5), which resolved the sample into two distinct peaks. Fractions were evaluated via fluorescence spectroscopy, and the apo-protein was pooled based on the Em_450_/Em_340_ emission ratio.

#### Non-isotope-labeled untagged *E. coli* DHFR

The cell culture protocol for the non-isotope-labeled, untagged *E. coli* DHFR (WT and M20I) was identical to that of the His-tagged variants, utilizing the pET-28a(+) vector with the C-terminal 6×His-tag removed (GenScript Biotech, Nanjing, China).

The purification protocol was adapted from previously established methods outlined in Greisman *et al*. (2022)^32^. Cell pellets were resuspended in Lysis Buffer B (50 mM Tris, 1 mM EDTA, 1 mM DTT, pH 8.0) supplemented with 1 mg ml^−1^ lysozyme, Complete Inhibitor Cocktail, and 0.01% (v/v) nuclease, utilizing a ratio of 1.5 mL of buffer per 1 g of pellet. The cell suspension was sonicated on wet ice for 1-2 minute per gram of pellet (1 second on, 2 seconds off, amplitude 20). The resulting lysate was centrifuged at 30,000 × *g* for 25 minutes to pellet the insoluble material, and the supernatant was extracted. To precipitate nucleic acids, a 10% (w/v) streptomycin solution was slowly added to the clarified supernatant while stirring over 10 minutes to reach a final concentration of 1% (w/v). The mixture was stirred for an additional 10 minutes and centrifuged at 30,000 × *g* for 25 minutes to recover the supernatant. Next, solid ammonium sulfate was gradually added to the supernatant at 4°C with gentle stirring to reach 40% saturation. After an additional 10 minutes of stirring, the mixture was centrifuged (30,000 × *g*, 25 minutes) to remove the precipitated proteins, and the supernatant was collected. The ammonium sulfate concentration in this supernatant was subsequently increased to 90% saturation while stirring to precipitate the target DHFR protein. The solution was centrifuged once more, and the resulting protein pellet was collected.

The 90% ammonium sulfate pellet was resuspended in MTX Wash Buffer (0.2 M KH_2_PO4, 1 M KCl, 1 mM EDTA, 1 mM DTT, pH 6.0) until the solution achieved clarity. Affinity purification was performed using a self-packed Methotrexate (MTX)-Agarose column (30 ml column volume). The column was equilibrated with 3 CVs of MTX Wash Buffer prior to sample loading, maintaining a flow rate of 1-2 ml min^−1^. Following loading, the column was washed with 10-20 CVs of MTX Wash Buffer until the UV absorbance fell below 20 mAU. The DHFR protein was then eluted using 45 CVs of MTX Elution Buffer (50 mM K_2_B4O_7_·4H_2_O, 2 M KCl, pH 10.15). Eluate fractions were collected and evaluated via SDS-PAGE, and those containing the target protein were pooled. To eliminate the high salt concentration present from the elution step, the pooled fractions were substantially diluted and buffer-exchanged into IEX Exchange Buffer (50 mM Tris, pH 7.5) using an Amicon Stirred Cell. The buffer-exchanged sample was loaded onto a HiPrep™ Q Fast Flow 16/10 anion exchange column (20 ml column volume) that had been equilibrated with 3 CVs of IEX Buffer A (100 mM NaCl, 50 mM Tris, 1 mM TCEP, pH 7.5). Operating at a flow rate of 1-2 ml min^−1^, the column was washed with 20 CVs of IEX Buffer A. The protein was subsequently eluted utilizing a 20-CV linear gradient from 0% to 100% IEX Buffer B (400 mM NaCl, 50 mM Tris, 1 mM TCEP, pH 7.5). Elution fractions were collected and analyzed via SDS-PAGE. Finally, the pure fractions corresponding to the target DHFR protein were pooled, buffer-exchanged, and concentrated into ~100 mg ml^−1^ in the final working buffer (10 mM HEPES, 1 mM DTT, pH 7.0). The purified DHFR solution was divided into 5 µL aliquots, flash-frozen, and stored at −70 °C.

#### Isotopically labeled untagged *E. coli* DHFR

For NMR experiments, ^15^N-labeled and ^15^N,^13^C-labeled *E. coli* DHFR (WT and M20I) were prepared utilizing the same expression plasmids described above for the untagged variants. Briefly, *E. coli* BL21(DE3) cells harboring these plasmids were inoculated from frozen stocks into 10 ml TB medium and cultured overnight (12-16 h) at 37 °C and 250 rpm. The overnight cultures were harvested by centrifugation and washed three times with 0 °C 1× phosphate-buffered saline (PBS). The cells were then resuspended in fresh isotope-labeling production medium containing 1× M9 minimal salts, 4 g L^−1^ glucose (or ^13^C-glucose), 0.5 g L^−1 15^NH4Cl, 0.5 g L^−1^ (^15^NH4)_2_SO4, 2 mM MgSO4, 0.1 mM CaCl_2_, 1× MEM vitamin solution, and 50 µg ml^−1^ kanamycin. These production cultures were incubated at 37 °C and 250 rpm until reaching the mid-logarithmic phase (OD_600_ ≈ 0.6). Protein expression was then induced with 1 mM IPTG. Following induction, the cultures were shifted to 18°C for 16 hours, after which the cells were harvested by centrifugation (6,000 × *g*, 4 °C).

All downstream protein purification steps were identical to the protocol described for the non-isotope-labeled, untagged *E. coli* DHFR, except that all the samples were buffer-exchanged and concentrated into NMR buffer (75 mM KPi, 25 mM KCl, 1 mM EDTA, 1 mM DTT, pH 7.6).

#### Untagged *E. coli* GroEL and His-tagged *E. coli* GroES

The *E. coli* TG1 strain harboring the pOA plasmid was utilized to overexpress *E. coli* GroEL and 6xHis-tag GroES. Cells were inoculated from frozen stocks into TB medium and first cultured overnight (12-16 h) at 37°C and 250 rpm. This overnight starter culture was then used to inoculate fresh TB medium at a 1% (v/v) ratio. These production cultures were incubated at 37°C and 250 rpm until reaching the mid-logarithmic phase (OD_600_ ≈ 0.6). Protein expression was then induced with 0.4 mM IPTG. Following induction, the cultures were shifted to 30°C for 16 hours, after which the cells were harvested by centrifugation (6,000 × *g*, 4°C). The cell lysis was performed exactly as described above for non-isotope-labeled untagged *E. coli* DHFR. The resulting clarified supernatant was applied to a gravity-flow column packed with cOmplete™ His-Tag Purification Resin. The column flow-through containing the untagged GroEL was collected for separate downstream purification, while the Ni-NTA resin was retained for the isolation of His-tagged GroES.

For the purification of His-tagged GroES, the retained Ni-NTA resin was washed with Buffer A to eliminate non-specifically bound host proteins. The target GroES was then eluted using Buffer B. This eluate was concentrated and buffer-exchanged into a storage buffer (25 mM Tris-HCl, 150 mM KCl, 1 mM DTT, pH 8.0). Final purification was achieved via SEC using a HiLoad 16/600 Superdex 75 pg column. Elution fractions were evaluated by SDS-PAGE, and the pure GroES fractions were pooled and concentrated using centrifugal ultrafilters.

For the purification of untagged GroEL, the column flow-through was incubated in a 60°C water bath for 7 minutes to precipitate non-target proteins, yielding a milky white suspension. The clarified supernatant was recovered following centrifugation at 4,800 × *g* for 20 minutes. Solid ammonium sulfate was gradually added at 4°C with gentle stirring to achieve 67% saturation. The resulting precipitated proteins were pelleted at 4,800 × *g* for 20 minutes and resuspended in 20 mL of storage buffer (25 mM Tris-HCl, 1 mM DTT, pH 8.0). Acetone was subsequently added dropwise under rapid stirring to a final concentration of 45%. Following another centrifugation step (4,800 × *g*, 20 minutes), the sample resolved into three distinct layers (a low-density top layer, a floppy precipitate middle layer, and a high-density, dark yellow bottom layer). The middle-layer protein pellet was collected, concentrated, and buffer-exchanged into a storage buffer (25 mM Tris-HCl, 150 mM KCl, 1 mM DTT, pH 8.0). Final purification was performed by SEC using a HiLoad 16/600 Superose 6 pg column and the storage buffer, and the elution fractions were concentrated via centrifugal ultrafiltration after SDS-PAGE.

Both the purified GroEL and GroES were subjected to a rigorous quality control pipeline: (1) SDS-PAGE to visually verify purity; (2) fluorescence spectroscopy to confirm the absence of tryptophan (Trp) residues in GroEL, which exhibited the expected flat emission profile upon excitation at 295 nm; and (3) a malate dehydrogenase (MDH) refolding assay to confirm retained chaperone activity, as described previously^46^. Once identity, purity, and functional activity were confirmed, the proteins were aliquoted, flash-frozen, and stored at −70 °C.

### *In vitro* biophysical and biochemical characterization

#### Bis-ANS thermal unfolding assays

To evaluate protein thermal stability and assess the degree of molten globule formation, a bis-ANS fluorescence-based thermal unfolding assay was employed. The reaction mixtures consisted of 5 µM purified protein and 50 µM of the fluorescent probe bis-ANS, prepared in a buffer containing 20 mM Tris, 250 mM NaCl, and 750 µM NADPH. To monitor thermal denaturation, the samples were subjected to a continuous temperature ramp of 2 °C min^−1^ from 20 to 80 °C. The unfolding transition was quantified using a Cary Eclipse fluorescence spectrometer (Varian) by recording the increase in bis-ANS fluorescence with excitation and emission wavelengths set to 395 nm and 490 nm, respectively. Finally, all fluorescence measurements were corrected by subtracting the background signal obtained from a blank control sample, with only buffer and 50 µM bis-ANS. All unfolding experiments were performed in triplicate for each sample.

#### DHFR enzymatic assays and kinetic analysis

DHFR enzymatic activity was measured at room temperature (25 °C) by continuously monitoring the oxidation of NADPH, which corresponds to a decrease in absorbance at 340 nm, using a Cary 60 UV-Vis spectrometer (Agilent). All reactions were carried out in a standard assay buffer consisting of 20 mM Tris, 136 mM NaCl, and 2 mM DTT at pH 7.4. The assay was executed by preparing two 2X working solutions to ensure proper enzyme equilibration. Solution A contained 60 nM purified DHFR enzyme and 200 µM NADPH in assay buffer and was pre-incubated at room temperature for at least 10 minutes. Solution B contained 60 µM dihydrofolate (DHF) and 200 µM NADPH in assay buffer. The reaction was initiated by mixing equal volumes of Solution A and Solution B, yielding final assay concentrations of 30 nM DHFR, 200 µM NADPH, and 30 µM DHF. Upon mixing, the sample was immediately vortexed, transferred to a cuvette, and the absorbance at 340 nm was recorded. For inhibition studies, different concentrations of TMP were included in Solution B, yielding final concentrations of 0.1, 0.5, 1, 2, 5, 10, 20, 30, 40, 50, 100, 500, and 1,000 nM. For L28R, final TMP concentrations included 0.1, 10, 100, 200, 450, 600, 700, 850, 1,000, 2,000, 5,000, and 10,000 nM. All enzymatic activity measurements, across all protein variants and TMP concentrations, were performed in independent triplicates.

Stock solutions of NADPH and dihydrofolate (DHF) were prepared and calibrated spectrophotometrically prior to the assays. NADPH was dissolved in Milli-Q water, and its concentration was determined by measuring absorbance at 340 nm utilizing an extinction coefficient of 6,200 M^−1^ cm^−1^. DHF was initially dissolved in 0.1 M NaOH, and its concentration was calibrated in assay buffer at 282 nm using an extinction coefficient of 28,000 M^−1^ cm^−1^. To prevent degradation, DHF and NADPH stocks were aliquoted in light-protected vials and stored at −70 °C until use. DHF aliquots were used strictly within two weeks of preparation. For inhibition measurements, TMP stock solutions were prepared freshly by dissolving 50-200 mg in 1 ml dimethyl sulfoxide (DMSO). Due to its limited aqueous solubility, this concentrated stock was subsequently diluted at least 100-fold into the final assay buffer to prevent TMP precipitation.

Kinetic parameters *k*_cat_ and *K*_M_ were determined using a progress curve analysis approach adapted from previously reported methods^27^. Rather than measuring initial velocities across a series of varied DHF concentrations, *k*_cat_ and *K*_M_ were extracted directly from individual, continuous enzymatic progress curves. Raw absorbance data were first subjected to baseline correction by fitting a linear regression to the final 30 seconds of each progress curve that represents the depleted substrate plateau and subtracting this baseline from each progress curve respectively. The baseline-corrected absorbance values were subsequently converted to DHF concentrations utilizing an effective reaction extinction coefficient of 12,300 M^−1^ cm^−1^. This continuous DHF concentration curve was divided into uniform 12-second intervals. Within each discrete window, a linear regression was performed to determine the local slope, the negative of which represents the instantaneous reaction velocity. Concurrently, the mean DHF substrate concentration was calculated for that corresponding interval. Finally, the array of instantaneous reaction velocities and their corresponding substrate concentrations were fitted to the Michaelis-Menten equation to resolve *k*_cat_ and *K*_M_.

The half-maximal inhibitory concentration IC_50_ was determined by fitting the initial velocities across varied TMP concentrations to a standard three-parameter logistic dose-response model assuming a fixed Hill coefficient of −1:

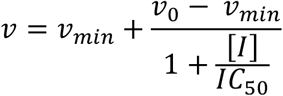

where *v* is reaction velocity at a given inhibitor concentration [*I*], *v*_0_ is the uninhibited reaction velocity, *v*_*min*_ is the baseline reaction velocity when the enzyme is fully inhibited. The inhibition constant *K*_i_ was then calculated from the derived IC_50_ values utilizing the Cheng-Prusoff equation:

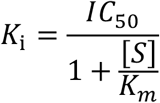

where [*S*] is DHF concentration.

#### GroEL/S rescue assays

To evaluate the GroEL/S-dependent recovery of M20I enzymatic activity *in vitro*, GroEL/S rescue assays were performed on WT and M20I with specific modifications to the standard enzymatic protocol described above. The assays utilized a two-step pre-incubation strategy to ensure stable DHFR-ligands binding and GroEL/S-mediated refolding reaction prior to initiating catalysis. DHFR, NADPH, and the respective TMP concentrations were first combined and pre-incubated at room temperature for 20 minutes. Subsequently, the pre-mixed GroEL and GroES solution was added to the mixture. For active rescue conditions, ATP was concurrently added at this stage. This combined mixture was incubated for an additional 20 minutes. Following equilibration, the enzymatic reaction was initiated by the addition of DHF alongside matching GroEL, GroES, NADPH, and/or ATP components to maintain constant concentrations. Upon mixing and brief vortexing, the decrease in absorbance at 340 nm was continuously monitored for 2 minutes. The final reaction mix contained 30 nM DHFR, 200 µM NADPH, 30 µM DHF, 3.5 µM GroEL (protomer), 3.5 µM GroES (protomer), 5 mM ATP and varying concentrations of TMP. All enzymatic activity measurements, across protein variants, GroEL/S and ATP conditions, and TMP concentrations, were performed in independent triplicates.

Since both DHFR enzymatic activity and GroEL/S refolding activity are highly sensitive to ionic strength, the standard assay buffer was replaced with a specialized working buffer consisting of 20 mM Tris, 10 mM MgCl_2_, 20 mM KCl, and 1.5 mM DTT at pH 7.4. Furthermore, to prevent non-specific protein adsorption at low working concentrations, all reaction mixtures were prepared in low-binding microcentrifuge tubes pre-treated with a Western blot blocking buffer. To mimic physiological conditions in *E. coli*, the standard GroEL/S to DHFR molar ratio was set to 8:1. Additionally, to evaluate how the recovery effect scales with GroEL/S availability, reactions utilizing an 80:1 ratio was also measured under 1,000 nM TMP.

Because the addition of the GroEL/S and ATP introduces baseline absorbance shifts, a dedicated background control containing a fully saturating concentration of TMP (10 mM) was measured for the active rescue group. The baseline rate obtained from this fully inhibited control was systematically subtracted from all corresponding experimental progress curves during data processing to yield the true chaperone-rescued DHFR kinetic rates. The determination of IC_50_ were conducted as described above.

#### Protein folding kinetics measurements

The spontaneous refolding kinetics of the WT, M20I, and W30C DHFR variants were measured at 25 °C by monitoring changes in intrinsic tryptophan fluorescence using a Jasco SFS-602T stopped-flow system equipped with a Jasco FDT-538 fluorescence detector (excitation at 282 nm, emission at 342 nm). All samples and denaturant solutions were prepared in a standard assay buffer (20 mM Tris, 136 mM NaCl, 2 mM DTT, pH 7.4). Briefly, DHFR variants were fully denatured in 6 M urea at a concentration of 10 µM. Refolding was initiated via rapid 1:10 dilution, achieved by co-injecting 1 volume of denatured protein with 9 volumes of standard buffer, yielding a final refolding condition of 1 µM DHFR in 0.6 M urea. Fluorescence signals were continuously recorded every 5 ms for 60 s, following an instrumental flow dead-time of 180 ms.

To ensure system equilibration, 10 consecutive injections were performed for each condition; the first three measurements served to flush the stopped-flow pipeline and were discarded, while the remaining seven replicates were retained for data processing. Raw fluorescence progress curves were background-corrected, and each of the seven independent replicates was individually fitted to a double-exponential decay function. This allowed for the extraction of the apparent rate constants for the fast and slow folding phases without assuming a specific mechanistic folding pathway.

#### Thermodynamic stability assays

Equilibrium unfolding and refolding experiments were performed to determine the thermodynamic stabilities of the apo WT and apo M20I DHFR variants. All protein samples and denaturant solutions were prepared in a standard assay buffer (20 mM Tris, 136 mM NaCl, 2 mM DTT, pH 7.4). For the unfolding pathway, 200 µM of native DHFR was diluted into 25 distinct concentrations of guanidinium chloride (GdnHCl) to a final protein concentration of 4 µM and incubated for at least 30 minutes. For the refolding pathway, 200 µM DHFR was first fully denatured in 4 M GdnHCl for 30 minutes, then diluted into 25 GdnHCl concentrations with final protein concentration of 4 µM and incubated overnight at 4 °C to ensure complete equilibration.

Measurements were performed at 25 °C using a Cary Eclipse fluorescence spectrophotometer (Varian). Samples were excited at 282 nm, and emission spectra were recorded from 300 nm to 800 nm. Upon unfolding, the intrinsic tryptophan emission peak of DHFR red-shifts from 342 nm (native) to 357 nm (fully unfolded). The fluorescence intensity ratio at these two wavelengths (I_342_/I_357_) was extracted to monitor the folded state across the denaturant gradient. These ratios were plotted against GdnHCl concentrations and fitted to a two-state thermodynamic model to determine the transition midpoint (C_m_), representing the denaturant concentration at which the protein population is 50% unfolded.

### GroEL-DHFR interaction analyses

#### *In vitro* binding and SEC co-elution

To assess the physical interaction between GroEL and the fully liganded DHFR ternary complex (DHFR:NADPH:TMP), size-exclusion chromatography (SEC) coupled with Western blotting was performed on both the WT and M20I variants. All protein samples and the SEC mobile phase were prepared using a standard GroEL storage buffer (25 mM Tris, 150 mM KCl, 1 mM DTT, pH 8.0). DHFR was first pre-incubated with NADPH and TMP for 30 minutes to ensure ternary complex formation, yielding pre-mix concentrations of 90 µM DHFR, 200 µM NADPH, and 200 µM TMP. An equal volume of 90 µM GroEL (protomer) solution was subsequently added to the DHFR mixture to initiate binding. Following a 30-minute incubation at 22 °C, the mixed samples were resolved on a Superdex 200 Increase 10/300 GL analytical column. To accurately assign the elution peaks, individual components (GroEL alone, DHFR alone, and the NADPH+TMP buffer) were run independently as reference standards. Equal volumes of the fractions corresponding to the high-molecular-weight GroEL elution peak were collected, resolved by SDS-PAGE, and subjected to Western blotting to confirm the presence of GroEL and co-eluting DHFR. The binding and SEC co-elution experiments were performed in independent duplicates for each DHFR variant.

#### *In vivo* crosslinking and pull-down assays

To assess the strength of endogenous GroEL-DHFR interactions *in vivo*, chemical crosslinking coupled with co-immunoprecipitation and Western blotting was performed for the WT, M20I, and W30C variants. Isogenic *E. coli* strains harboring the M20I and W30C single mutations within the chromosomal *folA* gene were engineered from the BW25113 background utilizing CRISPR-Cas9; the unmodified BW25113 strain served as the WT DHFR control. Overnight recovery cultures were inoculated from frozen cell stocks into Luria-Bertani (LB) medium and incubated for 12–16 hours at 37 °C and 250 rpm. These starter cultures were then diluted 1:100 (1% v/v) into fresh LB medium either in the presence or absence of 50 µg ml^−1^ TMP. Following a 12-hour incubation at 37 °C and 250 rpm, the cells were harvested, washed three times with ice-cold 1× PBS, and resuspended in ice-cold 1× PBS to a final OD_600_ of 5.0. All harvesting and washing steps were strictly performed on ice or at 4 °C to halt cellular processes. For *in vivo* crosslinking, 75 µl aliquots of the cell suspension were treated with DSP to a final concentration of 0.25 mM. The crosslinking reaction proceeded for 30 minutes at 25 °C with 250 rpm agitation. To terminate the reaction, an equal volume of quench buffer (100 mM Tris, pH 8.0) was added, and the samples were incubated for an additional 30 minutes under identical conditions before harvest.

Following crosslinking and quenching, the cell samples were lysed in 400 µL of Lysis Buffer C (1× B-PER™ reagent, 20 mM Tris, 150 mM NaCl, 0.05% Tween-20, 25 U ml^−1^ nuclease, cOmplete™ Protease Inhibitor Cocktail, pH 7.4) for 40 minutes at 25 °C with 250 rpm agitation. DHFR was then immunoprecipitated using rabbit anti-DHFR antibody-conjugated Pierce™ Protein A/G Magnetic Beads. The captured complexes were subsequently analyzed via Western blotting to detect and quantify DHFR alongside co-immunoprecipitated GroEL. To prevent signal overlap between the eluting rabbit IgG heavy chain (~50 kDa) and the GroEL protomer (~57 kDa), a mouse anti-GroEL primary antibody was specifically utilized for immunodetection. All crosslinking and immunoprecipitation experiments across all variants and conditions were performed in independent biological triplicates.

#### Endogenous DHFR abundance quantification

To quantify the endogenous steady-state expression levels of the WT and M20I DHFR variants, the unmodified BW25113 *E. coli* strain and the isogenic M20I mutant strain were utilized. Following overnight recovery in LB medium, starter cultures were inoculated into fresh LB medium either in the presence or absence of TMP at their respective 1× IC_50_ concentrations (3 µg m^−1^ for WT and 6 µg ml^−1^ for M20I). The cultures were incubated at 37 °C with 250 rpm agitation and harvested precisely at the mid-logarithmic growth phase, corresponding to OD_600_ of ~0.6. Following harvest, cell lysis and Western blotting were performed exactly as described above to detect and quantify the cellular abundance of the DHFR proteins under both unstressed and TMP-inhibited conditions.

### Cell growth assays

#### Cell growth IC_50_ determination with endogenous GroEL/S expression

Cell growth curves for the unmodified BW25113 *E. coli* strain and isogenic strains harboring single mutations (M20I, L28R, W30C) within the chromosomal *folA* gene were measured across varying TMP concentrations and temperatures to determine cell growth IC_50_. All cultures were grown in supplemented M9 minimal medium (1× M9 minimal salts, 0.4% glucose, 2 mM MgSO4, 0.1 mM CaCl_2_, 0.5 µg ml^−1^ thiamine, and 0.1% (w/v) casamino acids). Overnight recovery cultures were inoculated from frozen stocks into supplemented M9 minimal medium and incubated for 12– 16 hours at 37 °C and 250 rpm. These starter cultures were then diluted 1:100 (v/v) into fresh supplemented M9 minimal medium containing the respective TMP concentrations and grown for 6 hours at the target experimental temperatures to allow for cellular adaptation. For continuous growth curve data collection, 96-well plates were prepared with the outermost perimeter of wells filled with 200 µL of Milli-Q water to maintain humidity and minimize evaporation. The inner experimental wells were loaded with 100 µL of the adapted cell suspensions, normalized to an initial OD_600_ of 0.01. Cell density (OD_600_) was continuously monitored every 10 minutes using a BioTek Epoch microplate reader over a period of 10 to 24 hours. All cell growth measurements were performed in six independent biological replicates for each DHFR variant across all tested TMP concentrations and temperatures.

To accurately determine the cell growth IC_50_ for each DHFR variant under different temperatures, the resulting growth curve data were analyzed using a two-step mathematical fitting process. First, the background-corrected OD_600_ values from the initial 12 hours of growth were fitted to a four-parameter Gompertz growth model to extract the maximum specific growth rate *μ* for each condition:

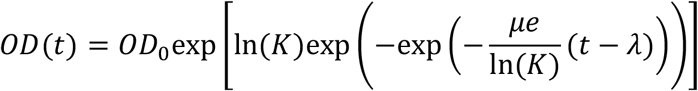

where *OD*(*t*) is the cell density at time *t, OD*_0_ is the initial baseline cell density, *K* is the carrying capacity (i.e., the maximum fold increase over the initial population at saturation), *λ* is the lag time, and *e* is Euler’s number. Subsequently, these extracted maximum specific growth rates *μ* were plotted against their respective TMP concentrations as dose-response curves and then fitted to a three-parameter logistic equation to calculate the final IC_50_ values:

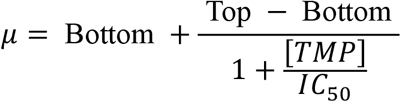

where Top and are Bottom the maximum and minimum growth rates. All curve fitting and parameter extraction were performed using custom Python scripts utilizing the SciPy optimization library.

#### Cell growth IC_50_ determination with GroEL/S knockdown

To assess how GroEL/S abundance affects TMP resistance, a two-plasmid CRISPR-dCas9 system was utilized to selectively knock down GroEL/S expression. The unmodified *E. coli* BW25113 strain and the isogenic mutant strains (M20I, L28R, W30C) harboring single mutations within the chromosomal *folA* gene were co-transformed with pdCas9-bacteria and either pgRNA-bacteria-groSP (for GroEL/S knockdown) or pgRNA-bacteria-empty (as control). Expression of the dCas9 protein was induced by anhydrotetracycline (aTc), while the guide RNA (gRNA) was constitutively expressed. Overnight recovery cultures were inoculated from frozen stocks into LB medium containing 100 µg ml^−1^ ampicillin and 34 µg ml^−1^ chloramphenicol, and incubated for 12– 16 hours at 37 °C and 250 rpm. These starter cultures were subsequently diluted 1:100 (v/v) into fresh LB medium supplemented with 100 µg ml^−1^ ampicillin, 34 µg ml^−1^ chloramphenicol, and 10 ng ml^−1^ aTc. The diluted cultures were grown for an additional 6 hours at 37 °C and 250 rpm to ensure robust GroEL/S knockdown and cellular adaptation prior to TMP exposure. For continuous growth curve data collection in 96-well plates, this same supplemented LB medium was used. The plate setup and cell density monitor proceeded exactly as previously described, with the exception that the initial cell density was normalized to an OD_600_ of 0.05. Rich LB medium was specifically chosen for these assays to alleviate the combined cellular stress induced by GroEL/S depletion and the presence of multiple antibiotics, thereby facilitating measurable cell growth within a practical timeframe. All subsequent data processing and IC_50_ analyses were conducted exactly as described above.

#### Cell growth IC_50_ determination with GroEL/S overexpression

To achieve GroEL/S overexpression, the pGro7 plasmid was transformed into the unmodified *E. coli* BW25113 strain (WT) and the isogenic mutant strain (M20I), utilizing a pEmpty plasmid as a negative control. Overnight recovery cultures were inoculated from frozen stocks into a modified supplemented M9 minimal medium (containing 0.2% glucose instead of the standard 0.4%) containing 50 µg ml^−1^ kanamycin. These cultures were grown for 12–16 hours at 37 °C with 250 rpm agitation. The starter cultures were subsequently diluted 1:100 (v/v) into fresh supplemented M9 minimal medium containing 50 µg ml^−1^ kanamycin and 0.05% arabinose to induce expression. The diluted cultures were incubated for an additional 6 hours at 37 °C and 250 rpm to ensure robust GroEL/S overexpression and cellular adaptation prior to TMP exposure. The 96-well microplate setup, continuous growth curve data collection, and downstream analyses proceeded exactly as described above.

### NMR spectroscopy

#### Sample preparation

All the NMR samples consisted of 1 mM ^13^C/^15^N isotopically labeled untagged DHFR, 15 mM NADPH, 2 mM TMP, 10 mM glucose-6-phosphate, 10 U glucose-6-phosphate dehydrogenase, 1 mM DTT, 10% D_2_O, and 1 µg ml^−1^ DSS. Low-pressure/vacuum NMR tubes were utilized for all measurements, allowing for the complete removal of air and gas-tight sealing under a nitrogen atmosphere using a Schlenk line. All samples were freshly prepared and protected from light using aluminum foil prior to loading into the NMR spectrometer.

#### Data collection and analysis

3D NMR spectra (HNCA, HNCACB, and HNCOCACB) were acquired at 298 K on a 600 MHz Bruker AVANCE NEO NMR spectrometer equipped with QCI-F Cryoprobe, utilizing a uniformly ^15^N,^13^C-labeled M20I sample. Raw NMR data was processed using NMRPipe^47^ and visualized with NMRDraw and NMR View^48^. Backbone resonances were assigned using standard triple-resonance methods within NMRFAM-Sparky^49^. All computational processing and assignment analyses were executed on a virtual machine hosted by the NMRbox platform^50^. Backbone resonances for the WT:NADPH:TMP ternary complex were previously reported^34^ and were confirmed to be consistent with the 2D [^1^H, ^15^N]-HSQC spectra collected in this study. The *nmrglue* package was used for 2D HSQC spectra plotting^51^. Chemical shift differences between M20I and WT were quantified by calculating the weighted Euclidean distance^52^ of the amide chemical shifts using the following equation.

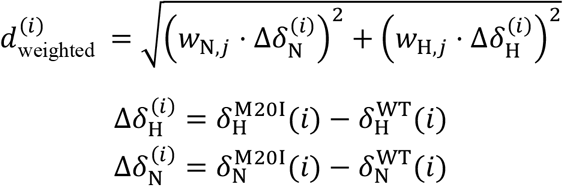

where 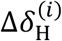 and 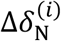 are measured chemical shift differences between M20I and WT for residue *i* along the H and N dimensions, respectively; *w*_N,*j*_ and *w*_H,*j*_ are amino-acid-specific weights obtained from Ref. 52, with *j* denoting the amino acid type corresponding to residue *i*.

To quantify the millisecond-timescale conformational dynamics of the DHFR variants, ^15^N Carr-Purcell-Meiboom-Gill (CPMG) relaxation dispersion experiments were performed on uniformly ^15^N labeled DHFR M20I. Data were collected at 298 K using a 700 MHz Bruker AVANCE NEO spectrometer equipped with an ultra-high sensitivity helium-cooled TCI CryoProbe. Measurements utilized a temperature-compensated, constant-time CPMG pulse sequence^53^ to ensure uniform sample heating across all relaxation delays. The total constant-time relaxation period was set to 40 ms. A series of 28 interleaved 2D spectra were acquired, corresponding to CPMG refocusing frequencies (ν_CPMG_) of 25, 50, 75, 100, 150, 175, 200, 225, 250, 275, 300, 350, 400, 450, 500, 550, 600, 650, 700, 750, 800, and 875 Hz. Duplicate measurements were collected at ν_CPMG_ of 100, 650, and 875 Hz, alongside a reference spectrum. Reproducibility across duplicate measurements was evaluated as part of data quality control but was not used for error estimation.

The raw NMR data were processed using NMRPipe^47^, and peak intensities were extracted using PeakFit^54^. The extracted intensity data were subsequently analyzed to extract exchange parameters using the ChemEx software package^55^. Errors were estimated based on peak integration by PeakFit.

Given the susceptibility of the NADPH to degradation via air and light exposure, strict quality control measures were employed for all multidimensional NMR acquisitions. A reference 2D [^1^H, ^15^N]-HSQC spectrum was acquired immediately before and after every extended NMR measurement, including all 3D assignment experiments and ^15^N CPMG relaxation dispersion runs. Only datasets exhibiting superimposable pre- and post-acquisition spectra, verifying the complete structural integrity of the DHFR ternary complex throughout the duration of the experiment, were retained for downstream analysis.

### X-ray crystallography

#### Sample preparation

To form the M20I:NADPH:TMP ternary complex, M20I was mixed with a threefold molar excess of NADPH and TMP, yielding the protein concentration of 22.5 mg ml^−1^, in a buffer containing 20 mM imidazole (pH 7.0) and 2 mM DTT, followed by a 30-minute incubation on ice. Crystallization was performed using the sitting-drop vapor-diffusion method, adapted from the conditions described by Greisman *et al*. (2022)^32^. The well solution consisted of 0.2 M imidazole, 0.25 M MnCl_2_, and 15–30% (v/v) PEG 400. Crystallization drops were set up by mixing 0.2 µL of the protein complex (adjusted to 12–22.5 mg ml^−1^) with 0.2 µL of the well solution. Rod-shaped crystals (approximately 0.1 × 0.3–0.5 mm) appeared after 2–4 weeks of incubation at 4 °C. The specific crystal selected for final data collection measured approximately 0.1 × 0.1 mm along the short axes and 0.5 mm along the long axis.

#### Data collection and analyses

A crystal of M20I:NADPH:TMP in a nylon loop and a B3S (ALS-style) goniometer base was covered with a MicroRT™ capillary (MiTeGen). The base of the sleeve was sealed with vacuum grease. The goniometer base was mounted on a Bruker D8 VENTURE with a PHOTON III detector operating in sensitivity mode and Cu IμS DIAMOND 3.0 rotating anode at 1.2 mA operating current. 180° of data were collected at 277 K and 1° φ oscillations with a 1 second exposure time. The beam was not attenuated and a 0.8 mm collimator was used. Data were reduced in DIALS v3.21.1 and cut to 1.9 Å resolution. The experimental geometry was provided to DIALS using a custom python script provided in the Supplementary Information. Isomorphous replacement and coordinate, B-factor, and occupancy refinement were performed in PHENIX v1.21.2 starting from PDB ID 6XG5. Model fit was manually inspected in COOT v0.9.8.96. The final structural model was deposited into the Protein Data Bank as PDB ID XXXX.

#### Statistics

All statistical analyses were performed using customized Python scripts mainly utilizing the NumPy, pandas, and SciPy libraries. Quantitative data are presented as the mean ± standard error of the mean (SEM), and error bars in all figures represent the SEM unless otherwise noted. Statistical comparisons between two independent experimental groups were conducted using an unpaired, two-tailed Student’s t-test. The threshold for statistical significance was set at an alpha level of 0.05. In all graphical representations, the exact levels of statistical significance are denoted by asterisks as follows: n.s. (not significant) indicates *P* > 0.05, * indicates *P* ≤ 0.05, indicates *P* ≤ 0.01, *** indicates *P* ≤ 0.001, and **** indicates *P* ≤ 0.0001.

#### Kinetic modeling

All the notations and values (if applicable) used in this kinetic model are listed in Supplementary Table 2.

Based on the kinetic model in fig. 5a, we have the following kinetic equations for M20I DHFR:

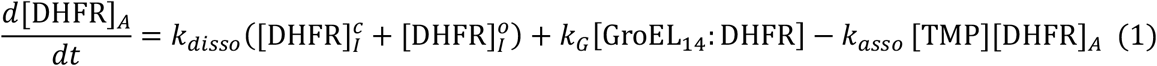

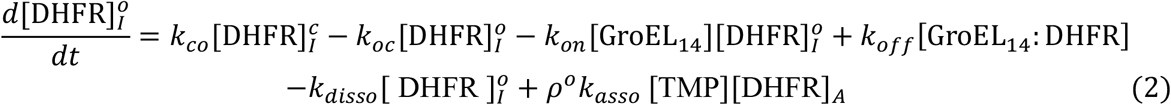

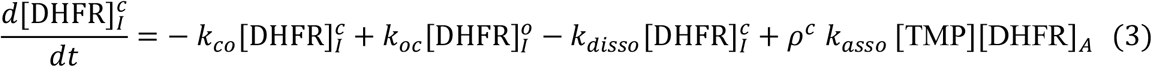

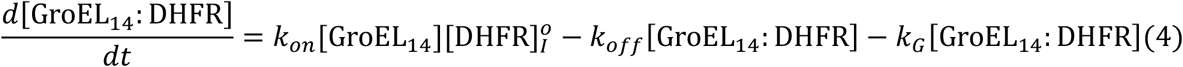

Where

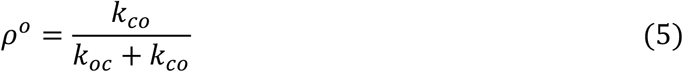

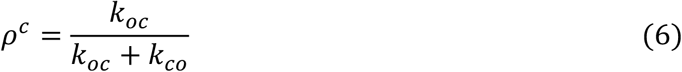

This model assumes total DHFR concentration is in its steady state. This is to say, DHFR production and degradation is not considered in this model. Therefore,

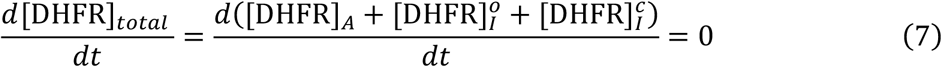

Because cell growth is directly coupled to the metabolic flux of THF, THF flux was used as the model output. Specifically, THF flux was formulated as a function of three classes of parameters: (i) the rate constants governing interconversion between the two conformational states; (ii) the relative population of the two states, represented by the free-energy difference between the open and closed conformations; and (iii) the binding affinity of GroEL for the open state. Therefore, we have the following equation:

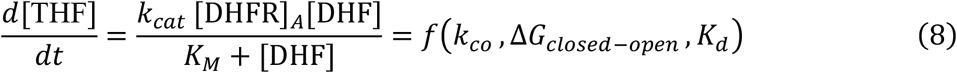

where

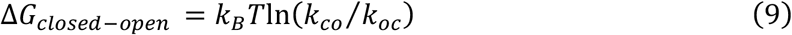

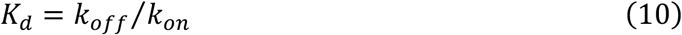

At steady state, setting Eq. 1–4 to zero gives the concentrations of the different protein conformations. We thus obtained:

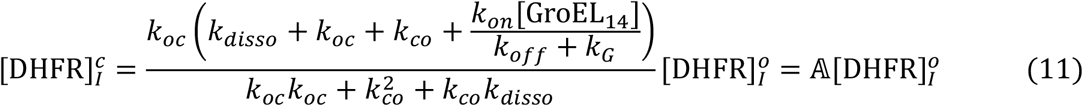

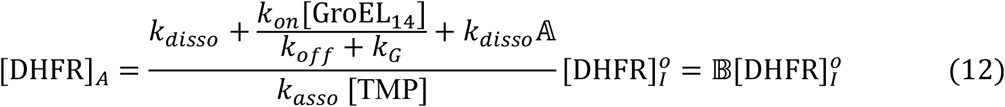

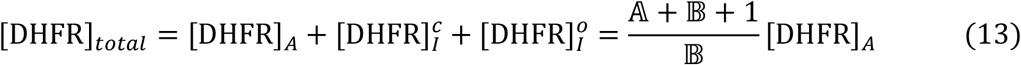

where equations 11 and 12 define 𝔸 and 𝔹. Therefore,

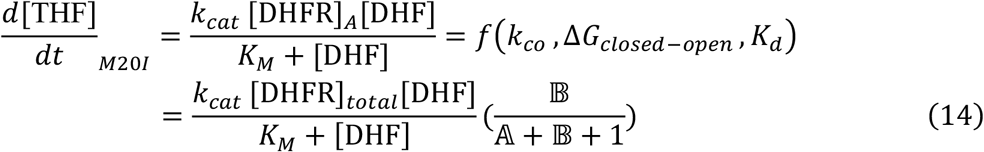

Although we experimentally observed weak GroEL_14_ recognition of WT DHFR (Fig. 3c, 3e-g, and Supplementary Fig. 17), we used a simplified model that does not include WT–GroEL_14_ engagement.

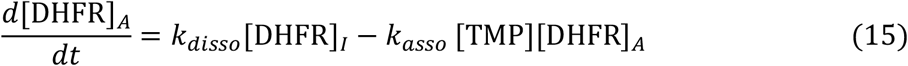

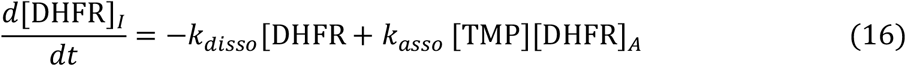

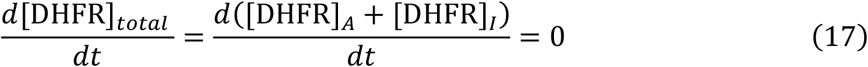

At steady state, we have

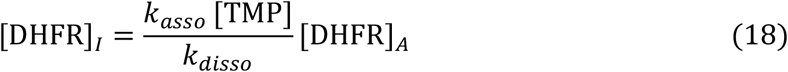

Therefore,

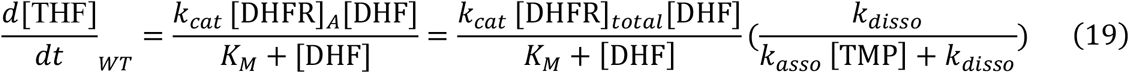

By varying *k*_*co*_, *k*_*oc*_, and *K*_*d*_, we assessed the relative contributions of conformational kinetics, conformational equilibrium, and GroEL_14_ binding affinity to the rescue of M20I growth, in comparison to WT.

Intracellular DHF/TMP competition after DHFR release is not explicitly included in this kinetic model. Previous measurements showed that intracellular DHF in *E. coli* rapidly accumulates upon treatment with 1–2 μM TMP, rising from undetectable baseline levels to the 10–100 μM range^56^. This range is comparable to the reported intracellular TMP concentration of ~89 μM^28^. Because association rate *k*_*asso*_ between DHFR and TMP or DHF are both on the order of 10^7^ M^−1^ s^−1 57,58^, the relative association flux 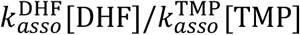 is expected to be on the order of 0.1–1. Thus, DHF and TMP are expected to compete on comparable kinetic scales for refolded active DHFR *in vivo*, providing a plausible kinetic window in which GroEL-mediated TMP removal can promote DHF rebinding and functional rescue.

### Molecular dynamics simulations

#### General simulation setup

Molecular dynamics (MD) simulations of the WT:NADPH:TMP and M20I:NADPH:TMP ternary complexes were performed using GROMACS 2023^59^. The systems were parameterized using the CHARMM36m force field^60^ for proteins, CGenFF v4.6^61,62^ for small-molecule ligands, the TIP3P water model for explicit solvation, and Beglov/Roux ion parameters^63^. Initial simulation boxes were constructed using the CHARMM-GUI Solution Builder v3.7^64,65^. System preparation proceeded through energy minimization followed by restrained constant volume (NVT) equilibration. All production simulations were conducted under constant pressure (NPT) with periodic boundary conditions. Temperature was maintained at 303.15 K using a velocity-rescaling thermostat (coupling constant of 1.0 ps) applied separately to the solute and solvent. Isotropic pressure was maintained at 1.0 bar using the C-rescale barostat with a 5.0 ps coupling constant and a compressibility of 4.5 × 10^−5^ bar^−1^. Nonbonded electrostatic and van der Waals interactions were truncated at 12 Å, with a force-switch modifier smoothly tapering van der Waals forces between 10 and 12 Å. Long-range electrostatics were computed using the Particle Mesh Ewald (PME) method. All bonds involving hydrogen atoms were constrained using the LINCS algorithm. An integration time step of 2 fs was employed, with the neighbor list updated every 20 steps (corresponding to a 40 fs interval). Hydrogen mass repartitioning, WYF cation–π parameters, and the multi-site Ca^2+^ model were not enabled. Trajectories were analyzed and visualized using VMD v1.9.3^66^, unless otherwise noted.

#### Initial configuration preparation

The crystal structure of *E. coli* DHFR in complex with NADPH and TMP (PDB ID: 6XG5) served as the structural template for both simulated complexes. For the mutant, the M20I substitution was introduced computationally using CHARMM-GUI. The protein chain and bound ligands were retained, and solvated in a rectangular water box containing TIP3P water molecules, maintaining a minimum solute-box edge distance of 10 Å. To neutralize the system net charge and mimic physiological conditions (pH 7.4), 0.15 M NaCl and appropriate counterions were introduced using a Monte Carlo ion placement method. Protein parameters were assigned using the CHARMM36m force field. The ligands (PDB hetero-IDs: NDP and TOP) were parameterized using the CHARMM General Force Field (CGenFF) via the CHARMM-GUI/ParamChem workflow, utilizing SDF files from the RCSB Protein Data Bank as initial inputs. Finally, GROMACS-formatted topology and parameter files were exported from CHARMM-GUI for all subsequent simulations.

#### Trajectory processing and analysis

Aggregate simulation times of 5 µs and 6 µs were collected for the WT:NADPH:TMP and M20I:NADPH:TMP ternary complexes, respectively. Initial visual inspection identified two major structural events: TMP dissociation (defined by a ligand–protein center-of-mass distance > 1 nm) in both systems and a pronounced outward displacement of the M20 loop specifically in the M20I system. Trajectories were segmented into 100-ns intervals using the GROMACS trjconv utility. Intervals capturing TMP dissociation or M20 loop opening were designated as transition intervals. To ensure robust analysis, all transition intervals and any trajectory segments following TMP dissociation were excluded. For the M20I complex, segments preceding the M20 loop transition were assigned to State 1 (closed conformation), while segments following the event were assigned to State 2 (open conformation).

Backbone root-mean-square fluctuation (RMSF) was calculated for each 100-ns segment using MDAnalysis^67,68^. Trajectory frames were first aligned to a reference structure using backbone atoms to eliminate global translational and rotational motions. RMSF was calculated for individual backbone atoms, and residue-level RMSF was derived by averaging the atomic values within each residue to facilitate comparative analysis between the WT and the two M20I conformational states.

### Bioinformatics

#### Comparative sequence analysis between *folA* and *dfrA* families

To evaluate the evolutionary divergence at the position corresponding to residue 20 in *E. coli* DHFR, sequence datasets for the canonical (*folA*) and TMP-resistant (*dfrA*) DHFR families were bioinformatically assembled. For the *folA* family, sequences were retrieved from both the RefSeq and UniProtKB databases using queries designed to deliberately exclude DHFR-like paralogs and known resistant variants. These datasets were merged, deduplicated based on exact amino acid identity, and queried against the DHFR Pfam profile (PF00186) using HMMER3 `hmmsearch`. After filtering out sequences containing gaps or non-standard amino acids at the position aligning with *E. coli* DHFR residue 20, a final dataset of 67,436 *folA* sequences was obtained. This dataset was subsequently aligned using MAFFT (`--auto` mode). An identical extraction, filtering, and alignment procedure was applied to the *dfrA* family, yielding a final set of 682 sequences sourced exclusively from the UniProt database. Using the resulting multiple sequence alignments, the compositional differences at the 20^th^ position between both protein families were statistically compared using a two-tailed Fisher’s exact test.

#### Evolutionary coupling analysis

Evolutionary coupling analysis to identify co-evolving residue pairs within DHFR was performed using the EVcouplings web server^69^. The *E. coli* DHFR sequence (UniProt ID: P0ABQ4) was submitted as the input query. The analysis was executed using the server’s default parameters, and the evolutionary couplings derived from the `target_b0.5` sequence alignment were utilized in this study.

### DHFR homolog fitness analyses

#### Source data

We re-analyzed a previously published mutational scanning dataset encompassing 1,536 designed DHFR homologs^35^. In the original study, these homologs and their mutants were synthesized via DropSynth into two distinct codon-version libraries (designated lib15/Codon 1 and lib16/Codon 2). The pooled libraries were assayed using an *E. coli* ER2566 Δ*folA* Δ*thyA* complementation system under both untreated and TMP-selection conditions, with barcode abundances quantified via sequencing (raw data available under NCBI BioProject PRJNA1189478).

For the present study, we directly utilized the processed barcode counts, barcode-to-variant assignments, and variant annotation tables provided by the source study rather than reprocessing the raw FASTQ files. To eliminate potential confounding effects from codon-specific biases in barcode representation or normalization, we restricted our primary fitness analysis to a single library (lib15/Codon 1), which was originally designed using weighted random assignment based on *E. coli* codon usage to optimize expression.

#### Barcode count normalization and fitness calculation

The fitness of DHFR homologs was recalculated from the lib15/Codon 1 processed count tables. Sample D05, corresponding to M9 medium without supplementation and lacking TMP, was used as the normalization baseline. Samples D06–D11 were assayed under varying TMP concentrations (0.058, 0.5, 1, 10, 50, and 200 μg ml^−1^, respectively). The barcode count tables across all TMP concentrations were merged by barcode identity after strict quality control. Following the protocols of the original study, a rigorous two-tier filtering strategy was applied: first, individual barcode sequences were retained only if they possessed ≥10 reads in at least one experimental condition; second, homologs were included only if they were represented by ≥5 of these high-confidence barcodes.

To quantify variant fitness, barcode read counts were processed as outlined in the original study. To account for variations in sequencing depth between different datasets, the raw count of each barcode *b* in condition *j* (*c*_*bj*_) was first normalized to the total sequencing depth of the baseline D05 condition. The normalized count (*r*_*bj*_) was calculated as:

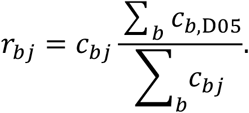

Following depth normalization, barcode fitness (*F*_*bj*_) was defined as the Log2 fold change in the normalized barcode recovery relative to the D05 baseline, incorporating a pseudocount of 1 to handle potential zero-count dropouts:

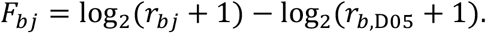

By definition, the fitness value for the baseline condition (D05) was fixed at 0. Finally, individual barcode-level fitness values were mapped to their corresponding homologs, and the overall fitness for each homolog or specific mutant was defined as the median fitness of its constituent barcodes at each TMP concentration.

#### Wild-type homologs fitness analyses

Following barcode filtering and variant-level fitness calculations, 961 unmutated DHFR homologs were retained for comparison. These sequences were aligned to the *E. coli* DHFR reference sequence (UniProt ID: P0ABQ4) using the procedure described above. Reference positions were defined strictly relative to the *E. coli* DHFR sequence. Homologs containing gaps or non-standard amino acids at either position 20 or 28 were excluded from the corresponding residue-state analyses.

Homologs were first grouped based on their residue identity at position 20, and the fitness trajectories of each group were plotted as a function of TMP concentration. Subsequently, homologs were grouped based on their specific residue pairs at positions 20 and 28. For each homolog, fitness values were fitted using the linear model:

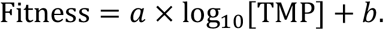

The intercept *b* was used as the fitted baseline fitness, corresponding to the estimated fitness at 1 μg ml^−1^ TMP. The absolute value of slope, |*a*|, was used as the TMP sensitivity, corresponding to the magnitude of fitness change per unit increase in log_10_[TMP]. The fitted baseline fitness and TMP sensitivity were evaluated for each residue-pair group by calculating median values and interquartile ranges. To ensure statistical robustness, only groups containing a minimum of 10 homologs were included in the analysis.

#### Fitness analysis of position 20 single mutants

For the 961 wild-type homologs analyzed above, corresponding variants containing a single amino acid substitution at reference position 20 were extracted from the lib15/Codon 1 library. The fitness of these variants was calculated using the identical pipeline described previously. A total of seven such single mutants were identified, and their corresponding TMP-response curves were plotted.

## Data availability

Structural data have been deposited in the PDB under accession code XXXX. Source data are provided with this paper.

## Code availability

Custom Python scripts used in this study are available in Github (to be added).

## Acknowledgements

We thank Dr. Shimon Bershtein (Ben-Gurion University of the Negev) for generously sharing the TG1/pOA bacterial strain and the GroEL/GroES purification protocol; Dr. Andrew L. Lee (University of North Carolina at Chapel Hill) for sharing NMR peak assignments; and Dr. Danny Fung (University of Wisconsin–Madison) for helpful discussions and suggestions regarding bacterial genetics. We thank Dr. Shao-Liang Zheng for assistance with X-ray data collection; Yifan Zhao for assistance with NMR tube sealing; Dr. Dennis E. Brookner for guidance on protein crystallization; and Tianyu Dong, Oliver Cheng, and Lilian Smith for experimental assistance during the early stages of this project. We also thank all current members of the Shakhnovich Biophysics Lab and the Hekstra Lab, especially Kibum Park and Dr. Dennis E. Brookner, for helpful discussions. This work was supported by the National Science Foundation (DGE2140743 to H.K.W.) and the National Institutes of Health (DP2-GM141000 to D.R.H. and R35GM139571 to E.I.S.). This work was performed in part at the Harvard University Laukien-Purcell Instrumentation Center (LPIC) and the CCB Center for Crystallographic Studies. We thank the support to the X-ray core facility from the Major Research Instrumentation (MRI) Program of the National Science Foundation (NSF) under Award Number 2216066. Computations for this project were performed on the FASRC Cannon cluster supported by the FAS Division of Science Research Computing at Harvard University.

## Author contributions

J.L. and E.I.S. conceived the study. J.V.R. performed the directed evolution experiments. J.L., Z.Z. and F.A. carried out molecular cloning. J.L. and Z.Z. established the CRISPR–Cas9 protocol and constructed the cell strains. J.L., Q.X. and Z.L. purified His-tagged DHFR proteins and performed enzymatic activity measurements. J.L. performed bis-ANS assays, endogenous DHFR abundance measurements, and GroEL–DHFR interaction assays. J.L. performed cell-growth assays, with assistance from F.A. Q.X. performed protein-folding kinetics measurements. J.L., Z.Z. and Q.X. performed thermodynamic stability assays. J.L. purified GroEL, GroES and all untagged DHFR proteins, including isotopically labelled samples. J.L. and Z.L. designed and performed the GroEL/S rescue assays. J.L. and F.A. quantified DHFR expression in promoter mutants. J.L. prepared NMR samples, acquired 3D NMR spectra with D.C., and performed HSQC peak assignments. A.D. acquired CPMG relaxation-dispersion data. J.L. analyzed CPMG data with assistance from A.D. J.L. performed protein crystallization. H.K.W. collected X-ray crystallography data and performed analysis. J.L. performed all other data analyses. J.L. and E.I.S. developed the kinetic model, and J.L. performed the numerical simulations. Z.Z. performed molecular dynamics simulations under the supervision of J.L. Z.Q. performed the bioinformatics analyses under the supervision of J.L. J.L. prepared and assembled the figures with input from Z.Q. J.L. and E.I.S. wrote the manuscript with input from all authors. D.R.H. acquired funding and supervised the X-ray crystallography component. E.I.S. acquired funding and supervised the project.

## Reference

1. Murray, C. J. L. et al. Global burden of bacterial antimicrobial resistance in 2019: a systematic analysis. The Lancet 399, 629–655 (2022).

2. Holohan, C., Van Schaeybroeck, S., Longley, D. B. & Johnston, P. G. Cancer drug resistance: an evolving paradigm. Nat. Rev. Cancer 13, 714–726 (2013).

3. Blair, J. M. A., Webber, M. A., Baylay, A. J., Ogbolu, D. O. & Piddock, L. J. V. Molecular mechanisms of antibiotic resistance. Nat. Rev. Microbiol. 13, 42–51 (2015).

4. Darby, E. M. et al. Molecular mechanisms of antibiotic resistance revisited. Nat. Rev. Microbiol. 21, 280–295 (2023).

5. Rebeaud, M. E., Mallik, S., Goloubinoff, P. & Tawfik, D. S. On the evolution of chaperones and cochaperones and the expansion of proteomes across the Tree of Life. Proc. Natl. Acad. Sci. 118, e2020885118 (2021).

6. Hartl, F. U., Bracher, A. & Hayer-Hartl, M. Molecular chaperones in protein folding and proteostasis. Nature 475, 324–332 (2011).

7. Hipp, M. S., Kasturi, P. & Hartl, F. U. The proteostasis network and its decline in ageing. Nat. Rev. Mol. Cell Biol. 20, 421–435 (2019).

8. Morales-Polanco, F., Lee, J. H., Barbosa, N. M. & Frydman, J. Cotranslational Mechanisms of Protein Biogenesis and Complex Assembly in Eukaryotes. Annu. Rev. Biomed. Data Sci. 5, 67–94 (2022).

9. Wentink, A., Rosenzweig, R., Kampinga, H. & Bukau, B. Mechanisms and regulation of the Hsp70 chaperone network. Nat. Rev. Mol. Cell Biol. 27, 110–128 (2026).

10. Diaz Arenas, C., Alvarez, M., Wilson, R. H., Shakhnovich, E. I. & Ogbunugafor, C. B. Protein Quality Control is a Master Modulator of Molecular Evolution in Bacteria. Genome Biol. Evol. 17, evaf010 (2025).

11. Iyengar, B. R. & Wagner, A. GroEL/S Overexpression Helps to Purge Deleterious Mutations and Reduce Genetic Diversity during Adaptive Protein Evolution. Mol. Biol. Evol. 39, msac047 (2022).

12. Bershtein, S., Mu, W., Serohijos, A. W. R., Zhou, J. & Shakhnovich, E. I. Protein quality control acts on folding intermediates to shape the effects of mutations on organismal fitness. Mol. Cell 49, 133–144 (2013).

13. Tokuriki, N. & Tawfik, D. S. Chaperonin overexpression promotes genetic variation and enzyme evolution. Nature 459, 668–673 (2009).

14. Cowen, L. E. & Lindquist, S. Hsp90 Potentiates the Rapid Evolution of New Traits: Drug Resistance in Diverse Fungi. Science 309, 2185–2189 (2005).

15. Kerner, M. J. et al. Proteome-wide Analysis of Chaperonin-Dependent Protein Folding in Escherichia coli. Cell 122, 209–220 (2005).

16. Chapman, E. et al. Global aggregation of newly translated proteins in an Escherichia coli strain deficient of the chaperonin GroEL. Proc. Natl. Acad. Sci. 103, 15800–15805 (2006).

17. Viitanen, P. V., Gatenby, A. A. & Lorimer, G. H. Purified chaperonin 60 (groEL) interacts with the nonnative states of a multitude of Escherichia coli proteins. Protein Sci. 1, 363–369 (1992).

18. To, P. et al. A proteome-wide map of chaperone-assisted protein refolding in a cytosol-like milieu. Proc. Natl. Acad. Sci. 119, e2210536119 (2022).

19. Wang, Z., Feng, H., Landry, S. J., Maxwell, J. & Gierasch, L. M. Basis of Substrate Binding by the Chaperonin GroEL. Biochemistry 38, 12537–12546 (1999).

20. Chen, L. & Sigler, P. B. The Crystal Structure of a GroEL/Peptide Complex: Plasticity as a Basis for Substrate Diversity. Cell 99, 757–768 (1999).

21. Stan, G., Brooks, B. R., Lorimer, G. H. & Thirumalai, D. Residues in substrate proteins that interact with GroEL in the capture process are buried in the native state. Proc. Natl. Acad. Sci. 103, 4433–4438 (2006).

22. Azia, A., Unger, R. & Horovitz, A. What distinguishes GroEL substrates from other Escherichia coli proteins? FEBS J. 279, 543–550 (2012).

23. Bandyopadhyay, B. et al. Local energetic frustration affects the dependence of green fluorescent protein folding on the chaperonin GroEL. J. Biol. Chem. 292, 20583–20591 (2017).

24. Bandyopadhyay, B., Mondal, T., Unger, R. & Horovitz, A. Contact Order Is a Determinant for the Dependence of GFP Folding on the Chaperonin GroEL. Biophys. J. 116, 42–48 (2019).

25. Rodrigues, J. V. et al. Biophysical principles predict fitness landscapes of drug resistance. Proc. Natl. Acad. Sci. 113, E1470–E1478 (2016).

26. Toprak, E. et al. Evolutionary paths to antibiotic resistance under dynamically sustained drug selection. Nat. Genet. 44, 101–105 (2012).

27. Tamer, Y. T. et al. High-Order Epistasis in Catalytic Power of Dihydrofolate Reductase Gives Rise to a Rugged Fitness Landscape in the Presence of Trimethoprim Selection. Mol. Biol. Evol. 36, 1533–1550 (2019).

28. Manna, M. S. et al. A trimethoprim derivative impedes antibiotic resistance evolution. Nat. Commun. 12, 2949 (2021).

29. Ptitsyn, O. b., Pain, R. h., Semisotnov, G. v., Zerovnik, E. & Razgulyaev, O. i. Evidence for a molten globule state as a general intermediate in protein folding. FEBS Lett. 262, 20–24 (1990).

30. Krucinska, J. et al. Structure-guided functional studies of plasmid-encoded dihydrofolate reductases reveal a common mechanism of trimethoprim resistance in Gram-negative pathogens. Commun. Biol. 5, 459 (2022).

31. Greisman, J. B. et al. Perturbative diffraction methods resolve a conformational switch that facilitates a two-step enzymatic mechanism. Proc. Natl. Acad. Sci. 121, e2313192121 (2024).

32. Greisman, J. B. et al. Native SAD phasing at room temperature. Acta Crystallogr. Sect. Struct. Biol. 78, 986–996 (2022).

33. Boehr, D. D., McElheny, D., Dyson, H. J. & Wright, P. E. The Dynamic Energy Landscape of Dihydrofolate Reductase Catalysis. Science 313, 1638–1642 (2006).

34. Mauldin, R. V., Carroll, M. J. & Lee, A. L. Dynamic Dysfunction in Dihydrofolate Reductase Results from Antifolate Drug Binding: Modulation of Dynamics within a Structural State. Structure 17, 386–394 (2009).

35. Romanowicz, K. J., Resnick, C., Hinton, S. R. & Plesa, C. Exploring antibiotic resistance in diverse homologs of the dihydrofolate reductase protein family through broad mutational scanning. Sci. Adv. 11, eadw9178 (2025).

36. Morcos, F. et al. Direct-coupling analysis of residue coevolution captures native contacts across many protein families. Proc. Natl. Acad. Sci. 108, E1293–E1301 (2011).

37. Marks, D. S. et al. Protein 3D Structure Computed from Evolutionary Sequence Variation. PLOS ONE 6, e28766 (2011).

38. Bloom, J. D., Labthavikul, S. T., Otey, C. R. & Arnold, F. H. Protein stability promotes evolvability. Proc. Natl. Acad. Sci. 103, 5869–5874 (2006).

39. Zeldovich, K. B., Chen, P. & Shakhnovich, E. I. Protein stability imposes limits on organism complexity and speed of molecular evolution. Proc. Natl. Acad. Sci. 104, 16152–16157 (2007).

40. Huot, M., Wang, D., Shakhnovich, E., Monasson, R. & Cocco, S. Constrained evolutionary funnels shape viral immune escape. Proc. Natl. Acad. Sci. 123, e2536956123 (2026).

41. Rutherford, S. L. & Lindquist, S. Hsp90 as a capacitor for morphological evolution. Nature 396, 336–342 (1998).

42. Çetinbaş, M. & Shakhnovich, E. I. Catalysis of Protein Folding by Chaperones Accelerates Evolutionary Dynamics in Adapting Cell Populations. PLOS Comput. Biol. 9, e1003269 (2013).

43. Papkou, A., Garcia-Pastor, L., Escudero, J. A. & Wagner, A. A rugged yet easily navigable fitness landscape. Science 382, eadh3860 (2023).

44. Bhattacharyya, S. et al. Transient protein-protein interactions perturb E. coli metabolome and cause gene dosage toxicity. eLife 5, e20309 (2016).

45. Zhang, Y., Chowdhury, S., Rodrigues, J. V. & Shakhnovich, E. Development of antibacterial compounds that constrain evolutionary pathways to resistance. eLife 10, e64518 (2021).

46. Ben-Zvi, A. P., Chatellier, J., Fersht, A. R. & Goloubinoff, P. Minimal and optimal mechanisms for GroE-mediated protein folding. Proc. Natl. Acad. Sci. 95, 15275–15280 (1998).

47. Delaglio, F. et al. NMRPipe: A multidimensional spectral processing system based on UNIX pipes. J. Biomol. NMR 6, 277–293 (1995).

48. Johnson, B. A. & Blevins, R. A. NMR View: A computer program for the visualization and analysis of NMR data. J. Biomol. NMR 4, 603–614 (1994).

49. Lee, W., Tonelli, M. & Markley, J. L. NMRFAM-SPARKY: enhanced software for biomolecular NMR spectroscopy. Bioinformatics 31, 1325–1327 (2015).

50. Maciejewski, M. W. et al. NMRbox: A Resource for Biomolecular NMR Computation. Biophys. J. 112, 1529–1534 (2017).

51. Helmus, J. J. & Jaroniec, C. P. Nmrglue: an open source Python package for the analysis of multidimensional NMR data. J. Biomol. NMR 55, 355–367 (2013).

52. Schumann, F. H. et al. Combined chemical shift changes and amino acid specific chemical shift mapping of protein–protein interactions. J. Biomol. NMR 39, 275–289 (2007).

53. Hansen, D. F., Vallurupalli, P. & Kay, L. E. An Improved 15N Relaxation Dispersion Experiment for the Measurement of Millisecond Time-Scale Dynamics in Proteins. J. Phys. Chem. B 112, 5898–5904 (2008).

54. Bouvignies, G. gbouvignies/PeakFit. (2026).

55. Bouvignies, G. gbouvignies/ChemEx. (2026).

56. Kwon, Y. K. et al. A domino effect in antifolate drug action in Escherichia coli. Nat. Chem. Biol. 4, 602–608 (2008).

57. Penner, M. H. & Frieden, C. Kinetic analysis of the mechanism of Escherichia coli dihydrofolate reductase. J. Biol. Chem. 262, 15908–15914 (1987).

58. Cayley, P. J., Dunn, S. M. J. & King, R. W. Kinetics of substrate, coenzyme, and inhibitor binding to Escherichia coli dihydrofolate reductase. Biochemistry 20, 874–879 (1981).

59. Abraham, M. J. et al. GROMACS: High performance molecular simulations through multi-level parallelism from laptops to supercomputers. SoftwareX 1–2, 19–25 (2015).

60. Huang, J. et al. CHARMM36m: an improved force field for folded and intrinsically disordered proteins. Nat. Methods 14, 71–73 (2017).

61. Huang, J. & MacKerell Jr, A. D. CHARMM36 all-atom additive protein force field: Validation based on comparison to NMR data. J. Comput. Chem. 34, 2135–2145 (2013).

62. Vanommeslaeghe, K. et al. CHARMM general force field: A force field for drug-like molecules compatible with the CHARMM all-atom additive biological force fields. J. Comput. Chem. 31, 671–690 (2010).

63. Beglov, D. & Roux, B. Finite representation of an infinite bulk system: Solvent boundary potential for computer simulations. J. Chem. Phys. 100, 9050–9063 (1994).

64. Jo, S., Kim, T., Iyer, V. G. & Im, W. CHARMM-GUI: A web-based graphical user interface for CHARMM. J. Comput. Chem. 29, 1859–1865 (2008).

65. Lee, J. et al. CHARMM-GUI Input Generator for NAMD, GROMACS, AMBER, OpenMM, and CHARMM/OpenMM Simulations Using the CHARMM36 Additive Force Field. J. Chem. Theory Comput. 12, 405–413 (2016).

66. Humphrey, W., Dalke, A. & Schulten, K. VMD: Visual molecular dynamics. J. Mol. Graph. 14, 33–38 (1996).

67. Gowers, R. J. et al. MDAnalysis: A Python Package for the Rapid Analysis of Molecular Dynamics Simulations. SciPy 2016 https://doi.org/10.25080/Majora-629e541a-00e (2016) doi:10.25080/Majora-629e541a-00e.

68. Michaud-Agrawal, N., Denning, E. J., Woolf, T. B. & Beckstein, O. MDAnalysis: A toolkit for the analysis of molecular dynamics simulations. J. Comput. Chem. 32, 2319–2327 (2011).

69. Hopf, T. A. et al. The EVcouplings Python framework for coevolutionary sequence analysis. Bioinformatics 35, 1582–1584 (2019).

