## Supplementary Materials for "Chaperonin recognition of protein dynamics drives drug resistance"

[1]Department of Chemistry and Chemical Biology, Harvard University, Cambridge, MA, USA. [2]Department of Cancer Biology, Dana-Farber Cancer Institute, Boston, MA, USA. [3]Department of Biological Chemistry and Molecular Pharmacology, Harvard Medical School, Boston, MA, USA. [4]Department of Molecular and Cellular Biology, Harvard University, Cambridge, MA, USA. [5]Graduate Program in Biophysics, Harvard University, Boston, MA, USA. [6]School of Engineering and Applied Sciences, Harvard University, Boston, MA, USA.

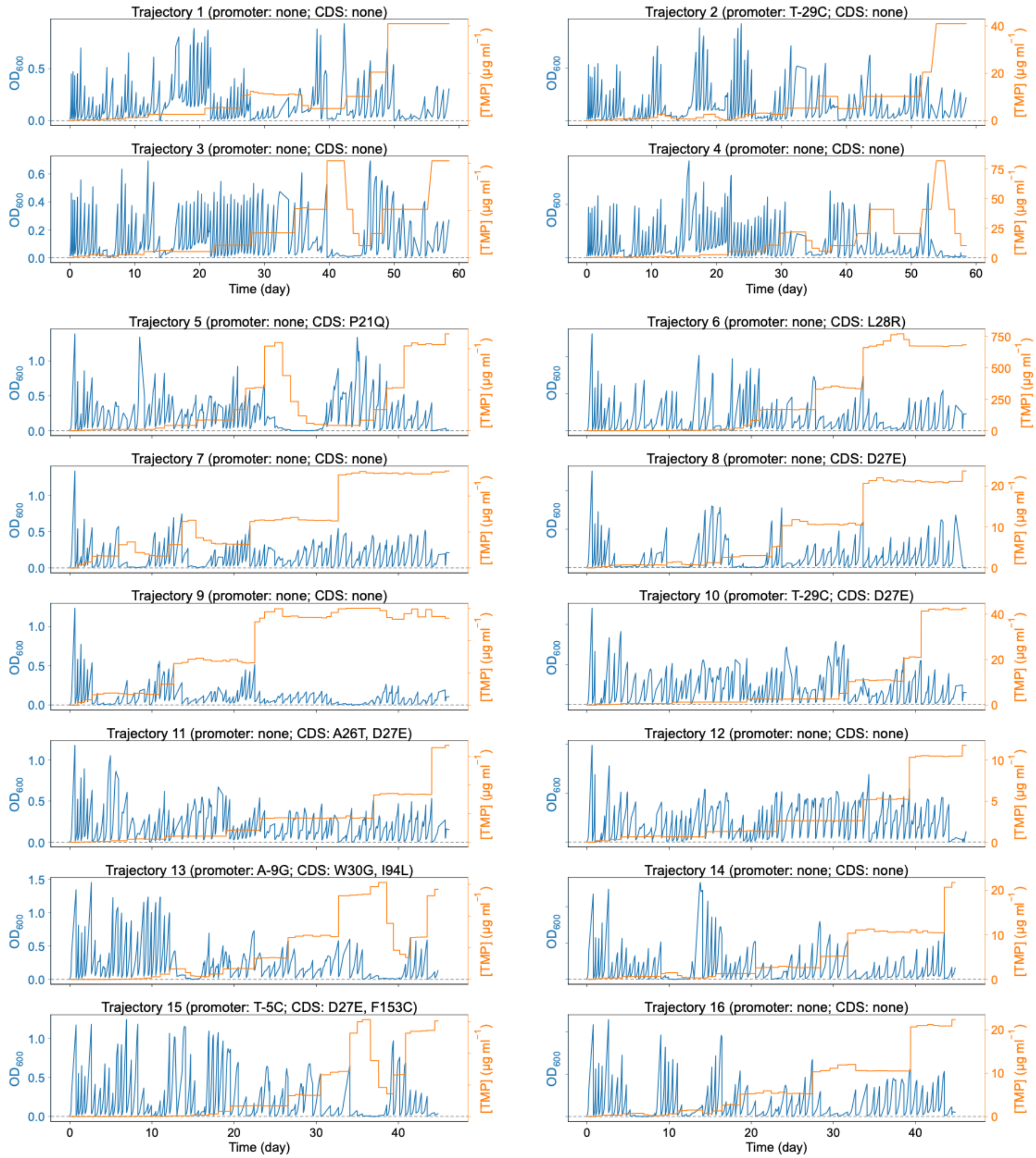


**Supplementary Fig. 1. Trajectories of TMP concentration and cell density for control group during evolution.** Sixteen parallel evolutionary trajectories were maintained for 45 days (~60 passages), with trajectories 1–4 extended to 60 days (~80 passages). Genomic mutations were identified from strains isolated at passage 60 for all trajectories. Trajectories 1–4 underwent additional sequencing at passage 80; any novel mutations emerging at this extended time point are explicitly annotated, whereas the absence of such labels indicates no further genetic changes were detected.

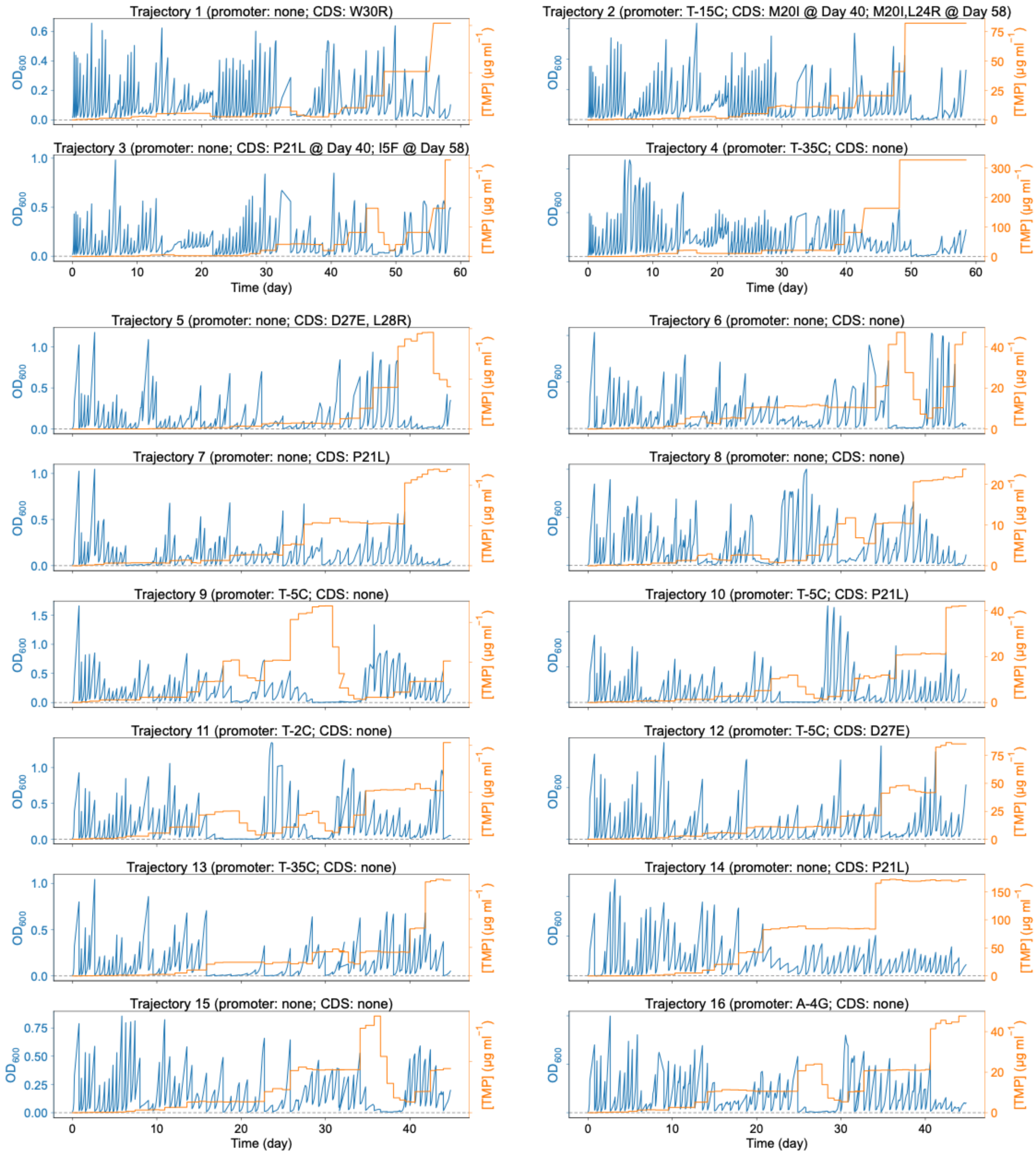


**Supplementary Fig. 2. Trajectories of TMP concentration and cell density for GroEL/S overexpression group during evolution.** Sixteen parallel evolutionary trajectories were maintained for 45 days (~60 passages), with trajectories 1–4 extended to 60 days (~80 passages). Genomic mutations were identified from strains isolated at passage 60 for all trajectories. Trajectories 1–4 underwent additional sequencing at passage 80; any novel mutations emerging at this extended time point are explicitly annotated, whereas the absence of such labels indicates no further genetic changes were detected.

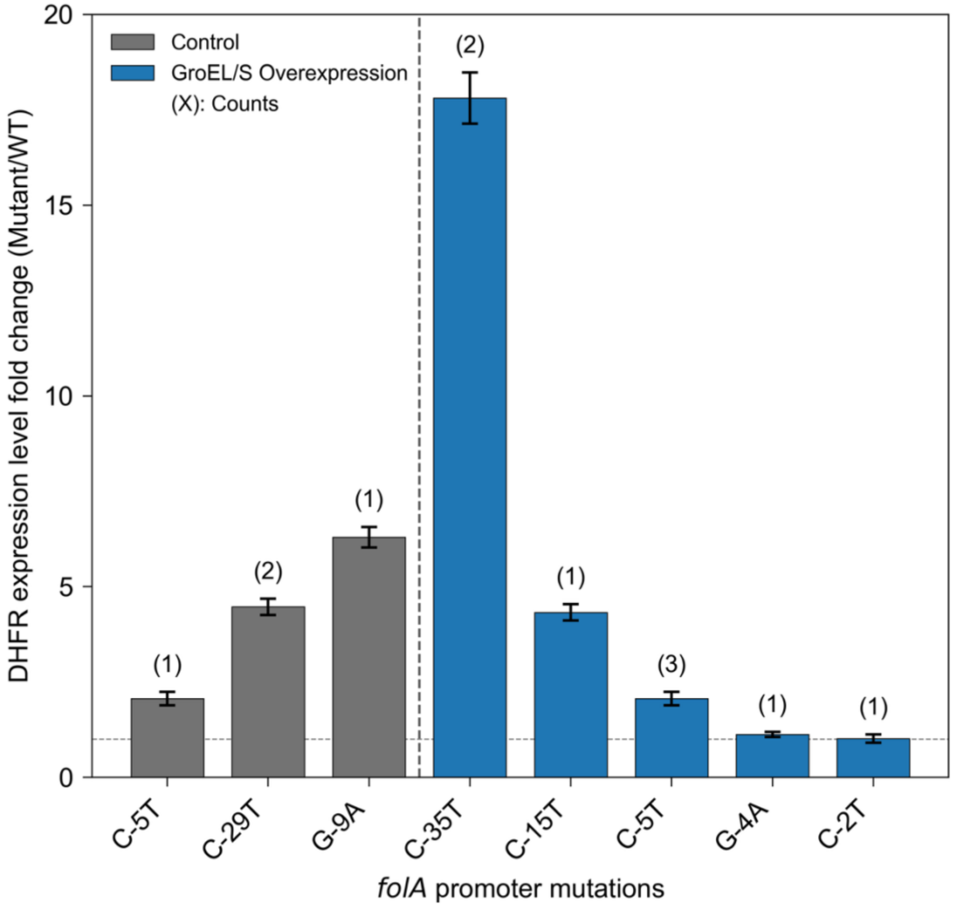


**Supplementary Fig. 3. Fold change in DHFR expression driven by *folA* promoter mutations identified through laboratory directed evolution.** GFP fluorescence was measured as a proxy for DHFR expression in *E. coli* DH5α strains harboring either the wild-type pfolA-GFP plasmid or its mutant variants. Briefly, overnight cultures were inoculated from frozen stocks into LB medium supplemented with 50 μg $ml^{-1}$ kanamycin and incubated for 12–16 h at 37 °C and 250 rpm. Starter cultures were subsequently diluted 1:100 (v/v) into fresh LB medium with with 50 μg $ml^{-1}$ kanamycin and grown for 6 h under identical conditions. Cell suspensions were normalized to an $OD_{600}$ of 0.1 prior to fluorescence measurements. Background-subtracted fluorescence intensities for each mutant were normalized to the wild-type promoter to calculate the fold change in expression. Data represent the mean ± SEM (standard error of the mean) of triplicate. Counts indicate the number of independent trajectories out of 16 total per group that acquired the specified mutations.

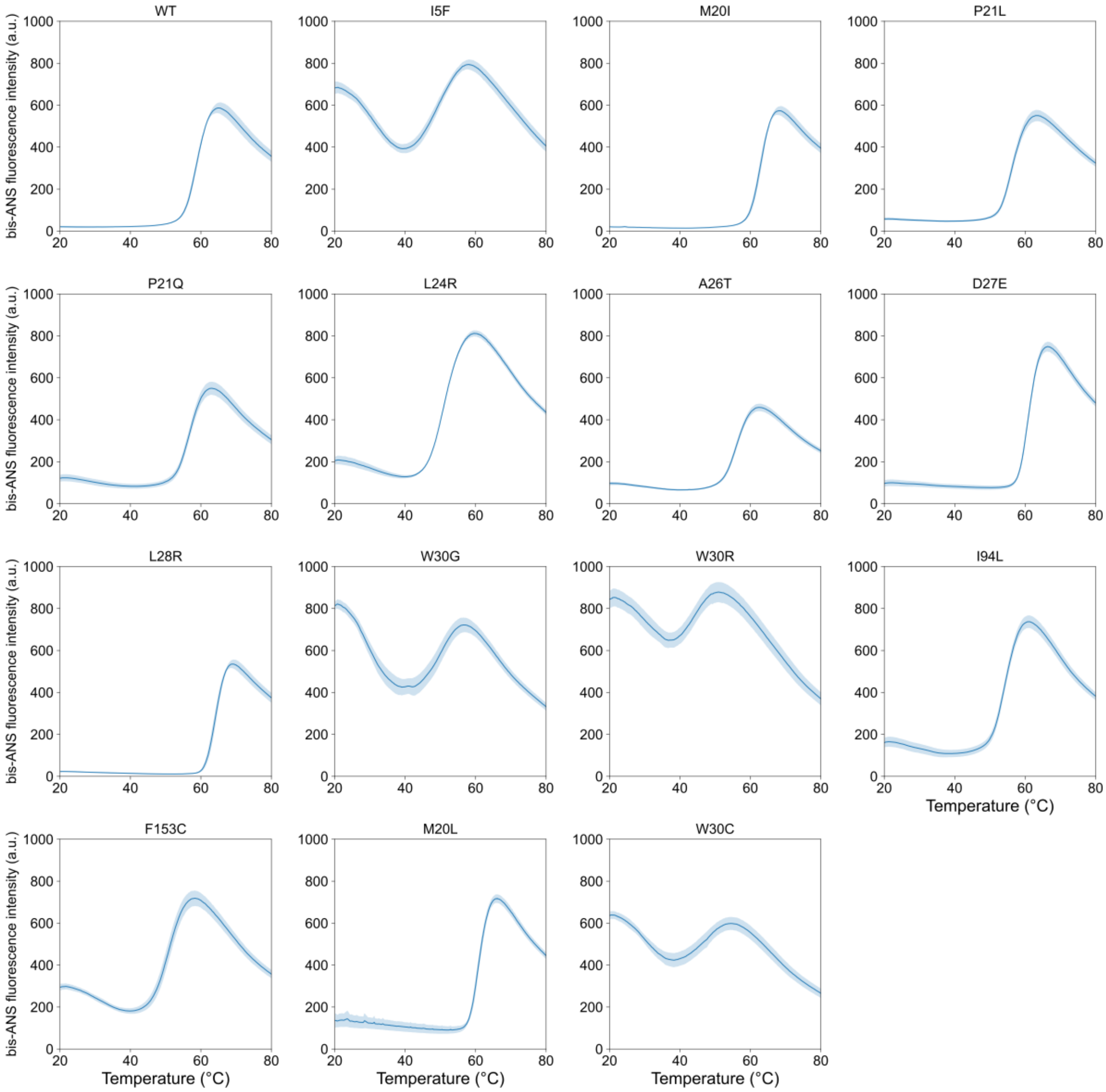


**Supplementary Fig. 4. Thermal unfolding profiles of WT DHFR and mutants monitored by bis-ANS fluorescence.** Solid blue lines represent the mean fluorescence intensity (arbitrary units, a.u.) across 3 independent replicates for each DHFR. The light blue shaded regions indicate the ± SEM.

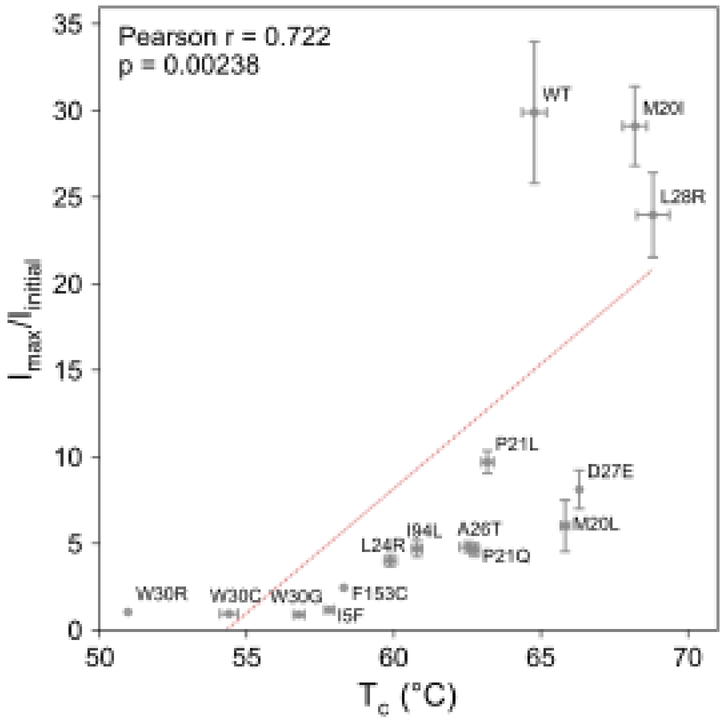


**Supplementary Fig. 5. Correlation between critical temperature ($T_c$) and molten globule degree ($I_{max}/I_{initial}$) across DHFR variants.** $I_{max}/I_{initial}$ and $T_c$ values were taken from Fig. 2b, revealing a statistically significant positive correlation (Pearson $r = 0.722$, $p = 0.00238$). Mutants with smaller $I_{max}/I_{initial}$ values exhibit more pronounced molten globule state, accompanied by a correspondingly lower $T_c$. The dashed line is shown as a guide to the eye.

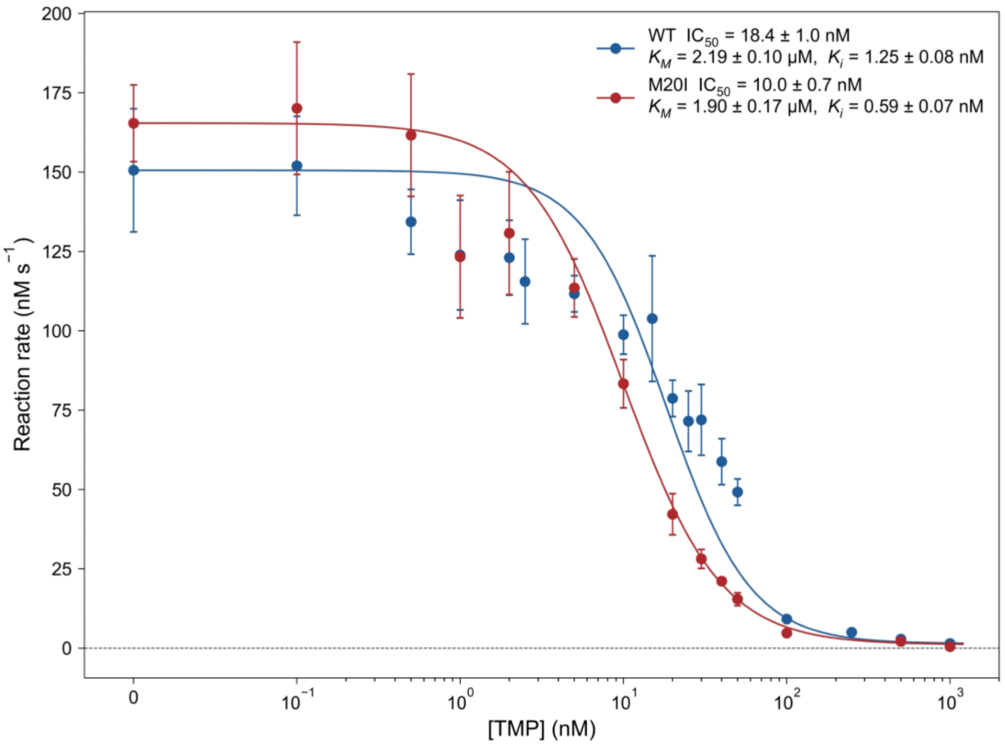


**Supplementary Fig. 6. TMP dose-response curves for WT and M20I DHFR.** *In vitro* reaction rates for WT and M20I DHFR were measured as a function of TMP concentration. Data represent the mean ± SEM of four independent replicates. Solid lines represent fits used to determine $IC_{50}$ and $K_i$ values.

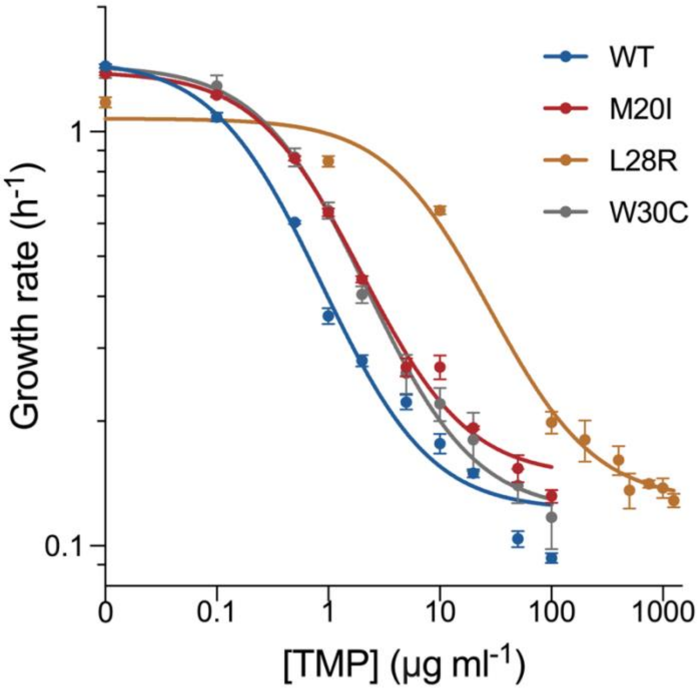


**Supplementary Fig. 7. TMP-response curves of WT and mutant DHFR strains.** Exponential growth rates (h$^{-1}$) measured at 37 °C are plotted as a function of TMP concentration (μg ml$^{-1}$) for WT, M20I, L28R, and W30C variants. Data points and error bars represent the mean ± SEM of 6 replicates. Solid lines indicate three-parameter logistic dose-response fits as described in Methods section.

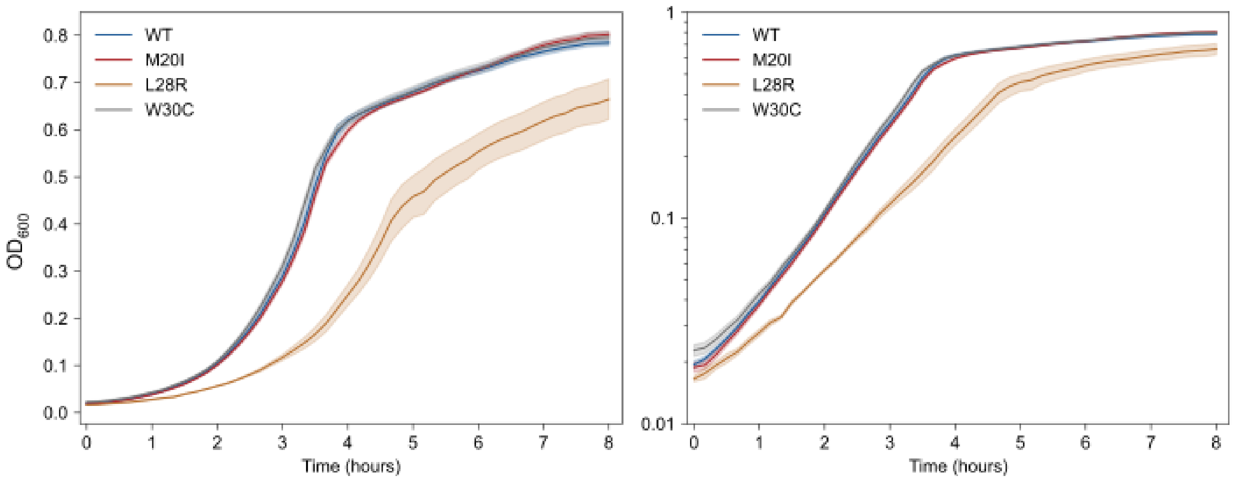


**Supplementary Fig. 8. Growth curves of WT and mutant DHFR strains.** Time-course $OD_{600}$ measurements of *E. coli* BW25113 strains expressing WT, M20I, L28R or W30C DHFR chromosomally in the absence of antibiotic stress at 37 °C. The left panel shows $OD_{600}$ on a linear scale, and the right panel shows the same data on a semi-log scale to highlight the exponential growth phase. Solid lines and shaded regions represent the mean ± SEM of 6 replicates.

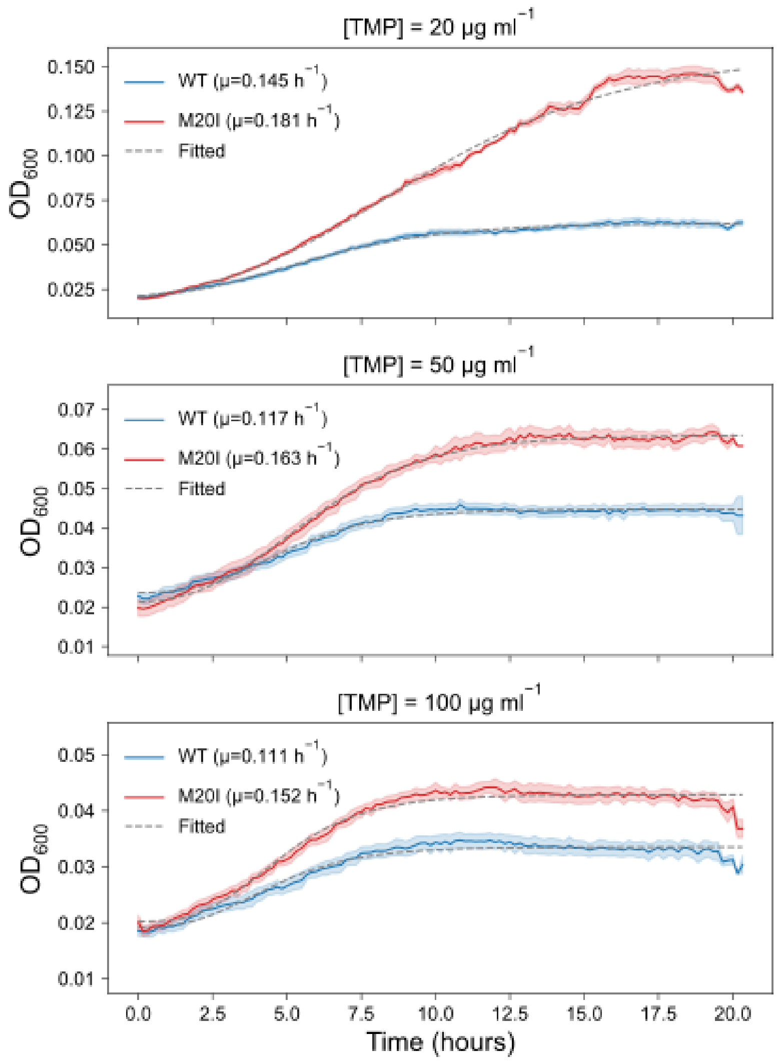


**Supplementary Fig. 9. TMP concentration-dependent growth curves for WT and M20I DHFR.** Cell density ($OD_{600}$) of WT and M20I strains was measured over 20 h in the presence of 20, 50, and 100 μg $ml^{-1}$ TMP at 37 °C. Solid lines and shaded regions represent the mean ± SEM of 6 replicates. Dashed lines represent the four-parameter Gompertz growth model fit to the experimental data, as detailed in Methods section. μ is maximum specific growth rate obtained by fitting the mean growth curves.

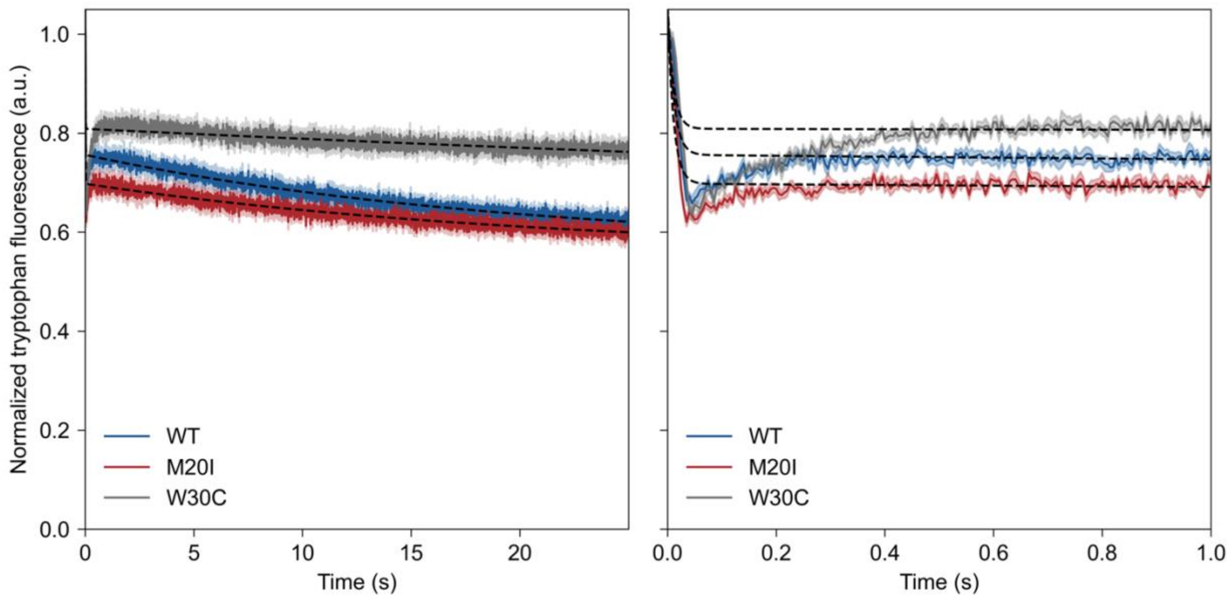


**Supplementary Fig. 10. Folding kinetics of DHFR variants monitored by stopped-flow tryptophan fluorescence.** The right panel displays a magnified view of the initial one-second folding phase from the full trace shown on the left. Solid lines and shaded regions represent the mean ± SEM of 7 independent replicates. Dashed black lines indicate double-exponential kinetic fits, as detailed in the Methods section.

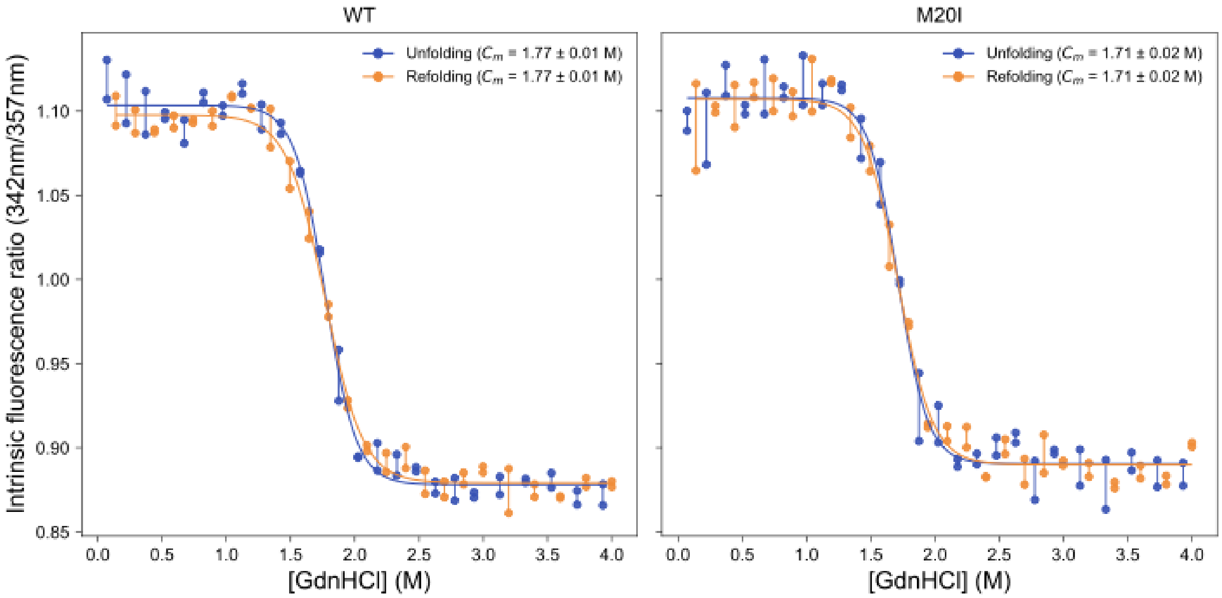


**Supplementary Fig. 11. Equilibrium unfolding and refolding of WT and M20I DHFR.** GdnHCl-induced denaturation of WT and M20I DHFR monitored by intrinsic fluorescence. Overlapping forward and reverse curves demonstrate thermodynamic reversibility. Calculated transition midpoints ($C_m$) are inset. Data points represent individual measurements from two replicates at each GdnHCl concentration and are connected by lines to guide the eye. Solid lines indicate the two-state model fit as described in Methods section.

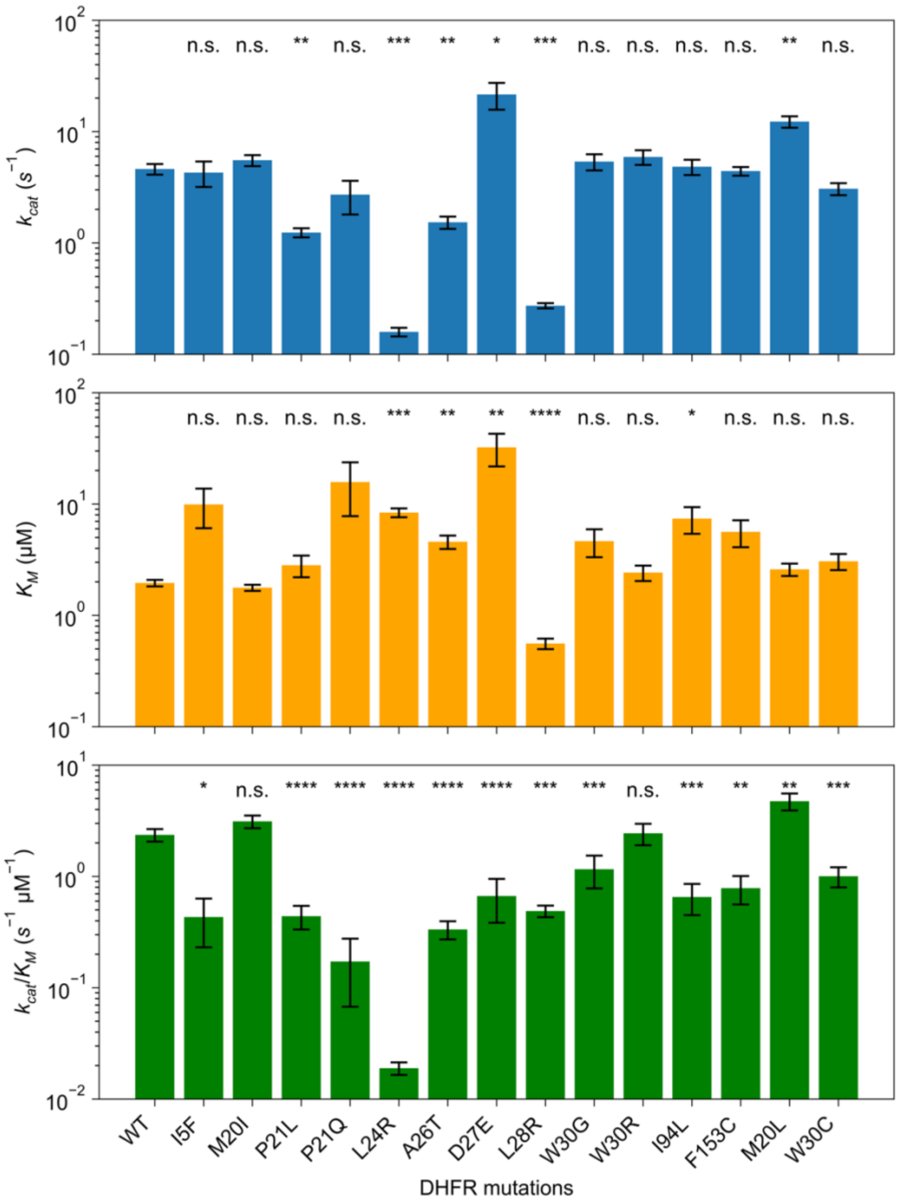


**Supplementary Fig. 12. Kinetic parameters ($k_{cat}$, $K_M$, and $k_{cat}/K_M$) for all DHFR variants.** Statistical significance was determined by comparing each individual mutant to WT.

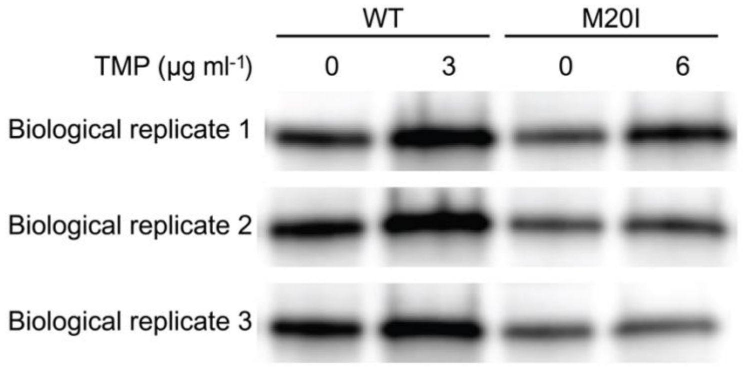


**Supplementary Fig. 13. Expression levels of chromosomally encoded WT and M20I DHFR.** Western blot analysis of DHFR from *E. coli* BW25113 expressing either WT or M20I DHFR chromosomally. Cultures were grown in the absence of TMP (0 µg $ml^{-1}$) or at a TMP concentration corresponding to the respective $IC_{50}$ for each strain (3 µg $ml^{-1}$ for WT and 6 µg $ml^{-1}$ for M20I). Data represent three independent biological replicates.

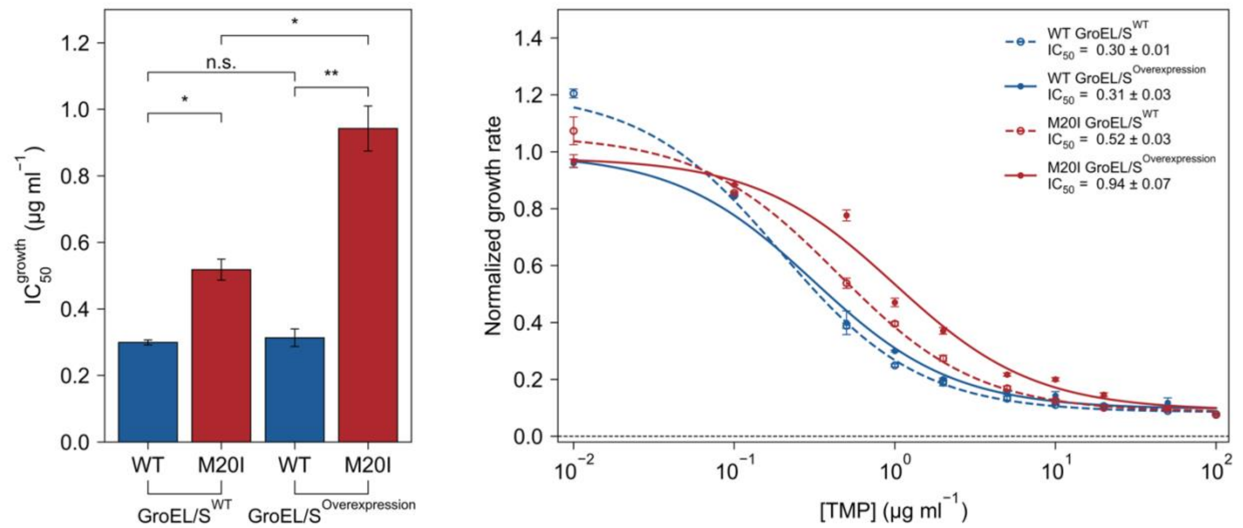


**Supplementary Fig. 14. GroEL/S overexpression selectively enhances TMP resistance of M20I. (Left)** $IC_{50}$ values derived from cell growth assays for WT and M20I DHFR strains in endogenous (GroEL/$S^{WT}$) versus GroEL/S overexpression backgrounds measured at 37 °C. Bars denote the mean ± SEM of 3 biological replicates. Statistical analysis highlights the specific enhancement of M20I resistance upon GroEL/S overexpression. **(Right)** Corresponding TMP dose-response curves at 37 °C. Normalized growth rates of WT and M20I strains are plotted as a function of TMP concentration (μg $ml^{-1}$). Dashed and solid lines represent three-parameter logistic fits for strains in the endogenous (GroEL/$S^{WT}$) and GroEL/S overexpression backgrounds, respectively. These fits were used to derive the inset $IC_{50}$ values, as detailed in the Methods section. Data points and error bars represent the mean ± SEM of 3 biological replicates.

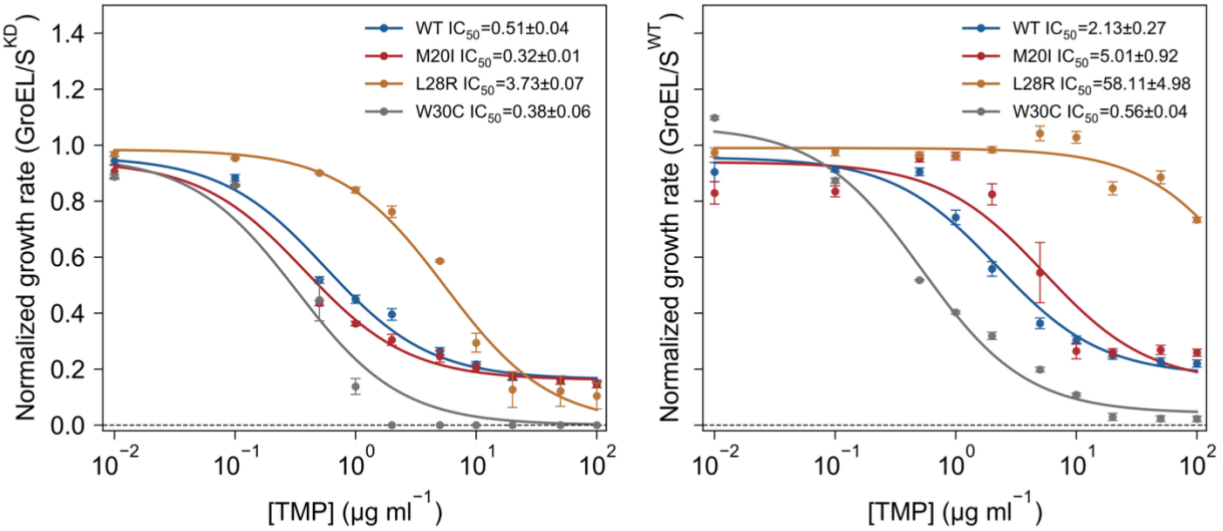


**Supplementary Fig. 15. TMP-response curves of DHFR variants under GroEL/S knockdown and endogenous conditions.** Normalized growth rates of WT, M20I, L28R, and W30C strains measured at 37 °C are plotted as a function of TMP concentration (μg $ml^{-1}$) in GroEL/S knockdown (left, GroEL/$S^{KD}$) and endogenous (right, GroEL/$S^{WT}$) backgrounds. Solid lines indicate three-parameter logistic dose-response fits used to derive the inset $IC_{50}$ values, as detailed in Methods section. Data points and error bars represent the mean ± SEM of 6 replicates.

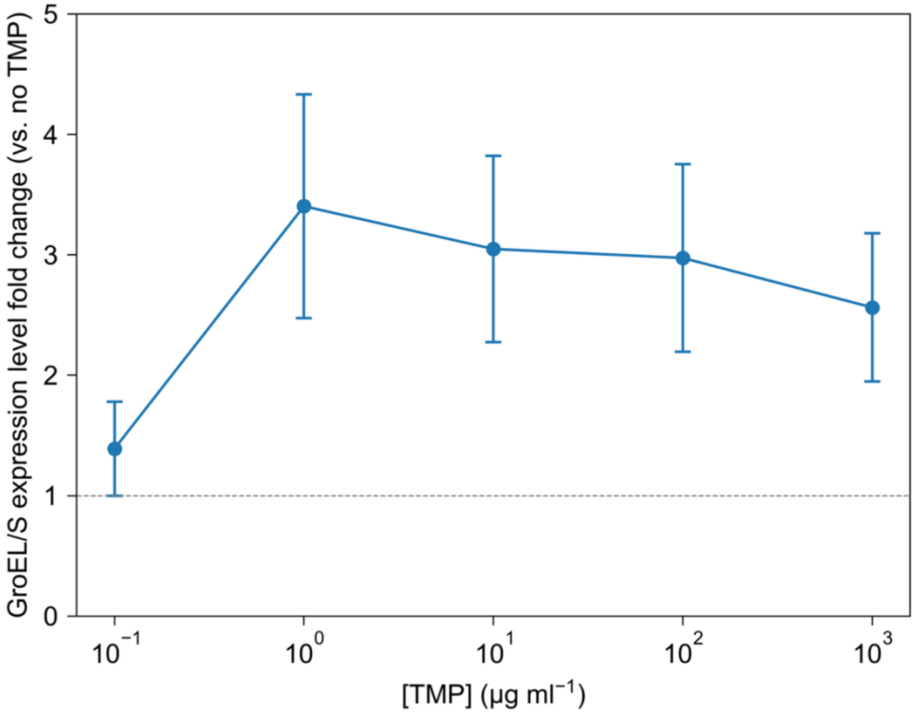


**Supplementary Fig. 16. Fold change in GroEL/S expression under varying concentrations of TMP.** GFP fluorescence was measured as a proxy for GroEL/S expression in *E. coli* DH5α strains harboring the pgroS-GFP plasmid. Briefly, overnight cultures were inoculated from frozen stocks into LB medium supplemented with 50 μg ml$^{-1}$ kanamycin and incubated for 12–16 h at 37 °C and 250 rpm. Starter cultures were subsequently diluted 1:100 (v/v) into fresh LB medium containing 50 μg ml$^{-1}$ kanamycin and the respective TMP concentrations, followed by growth for 6 h at 37 °C and 250 rpm to allow for cellular adaptation. Cell suspensions were normalized to an $OD_{600}$ of 0.1 prior to fluorescence measurements. To calculate the fold change in expression, background-subtracted fluorescence intensities for each TMP concentration were normalized to those measured in the untreated control (LB medium without TMP). Data represent the mean ± SEM of triplicate.

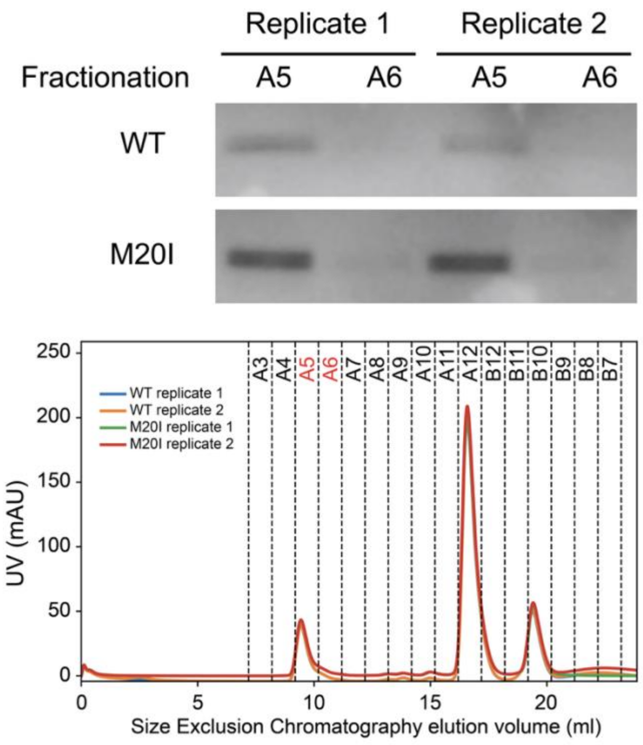


**Supplementary Fig. 17. SEC co-elution of GroEL and DHFR ternary complexes.** SEC chromatograms (bottom) of GroEL–DHFR mixtures capture the high-molecular-weight GroEL peak (red fractions A5, A6), unbound DHFR (second peak), and small-molecule ligands (third peak). Western blotting of the A5 and A6 fractions (top) reveals that while both WT and M20I DHFR co-elute with GroEL, the M20I variant exhibits a darker band intensity, indicating an enhanced physical interaction. All assays were performed in independent duplicates.

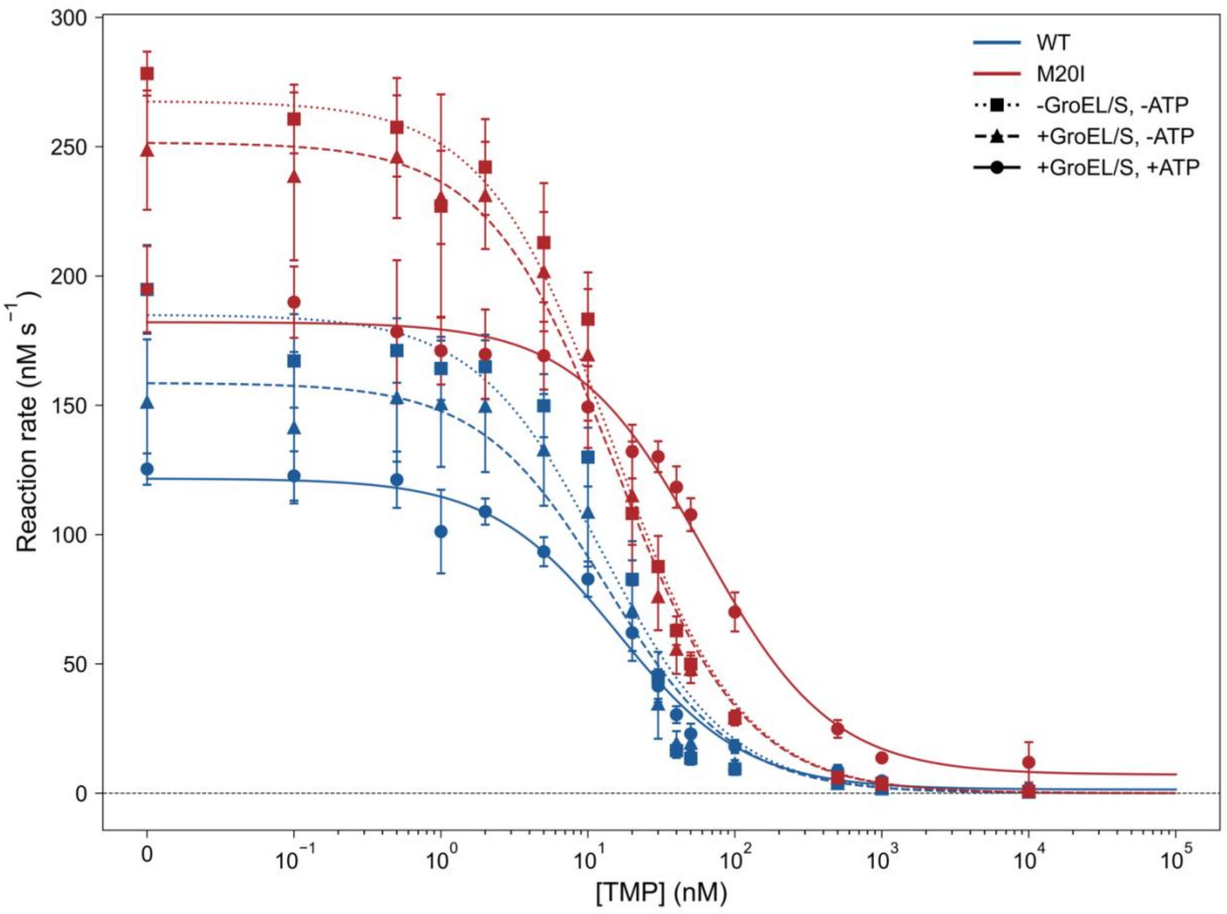


**Supplementary Fig. 18. GroEL/S-mediated rescue of DHFR activity.** *In vitro* reaction rates of WT and M20I DHFR plotted as a function of TMP concentration. Reactions were evaluated without chaperone (-GroEL/S, -ATP), with inactive chaperone (+GroEL/S, -ATP), and under active refolding conditions (+GroEL/S, +ATP). The M20I variant exhibits substantial functional recovery exclusively under active rescue conditions, even under extremely high TMP concentrations. Data represent the mean ± SEM of three independent replicates.

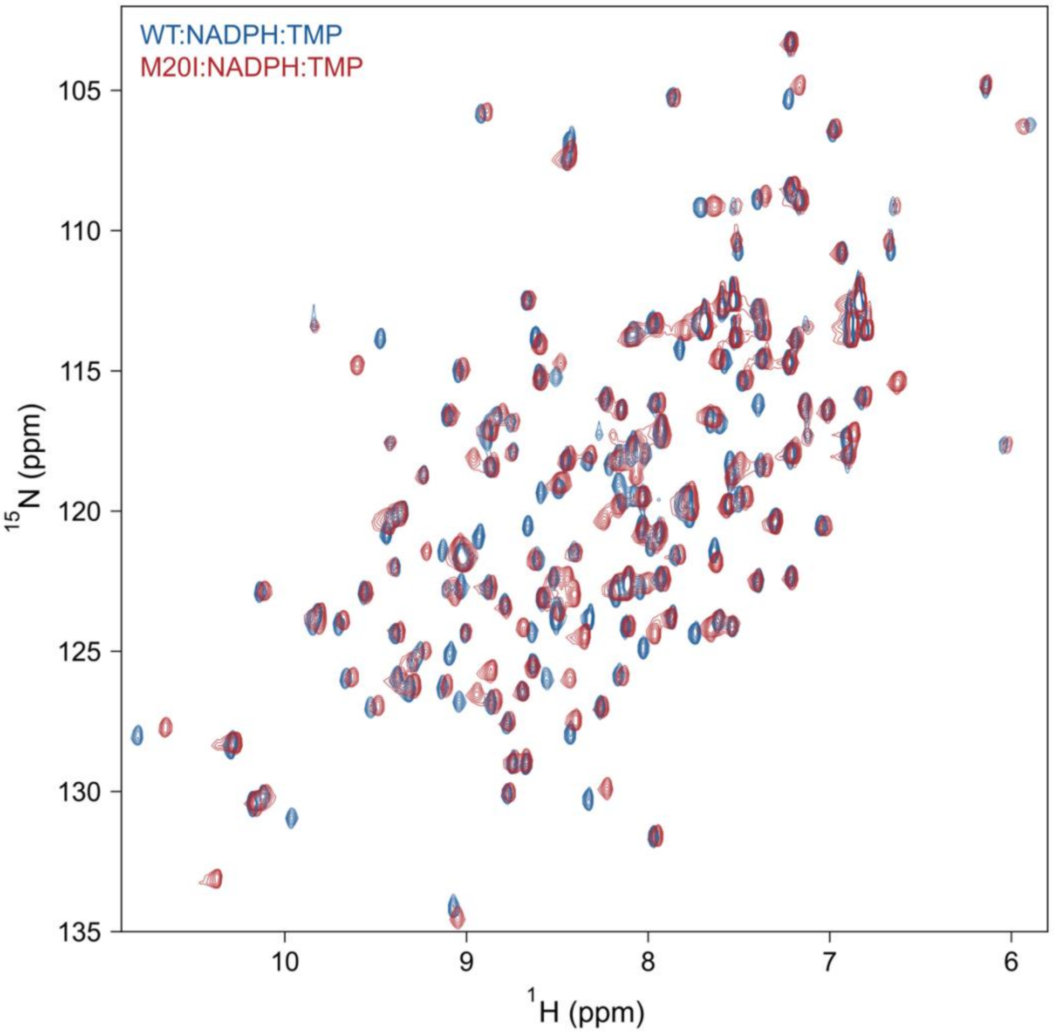


**Supplementary Fig. 19. 2D $^{1}$H-$^{15}$N HSQC spectra of WT and M20I DHFR.** Spectral overlay of the WT (blue) and M20I (red) DHFR ternary complexes fully liganded with NADPH and TMP. Data were acquired at 298 K at 600 MHz. The dispersion and alignment of the cross-peaks confirm that the mutant retains the WT global fold while exhibiting specific chemical shift perturbations.

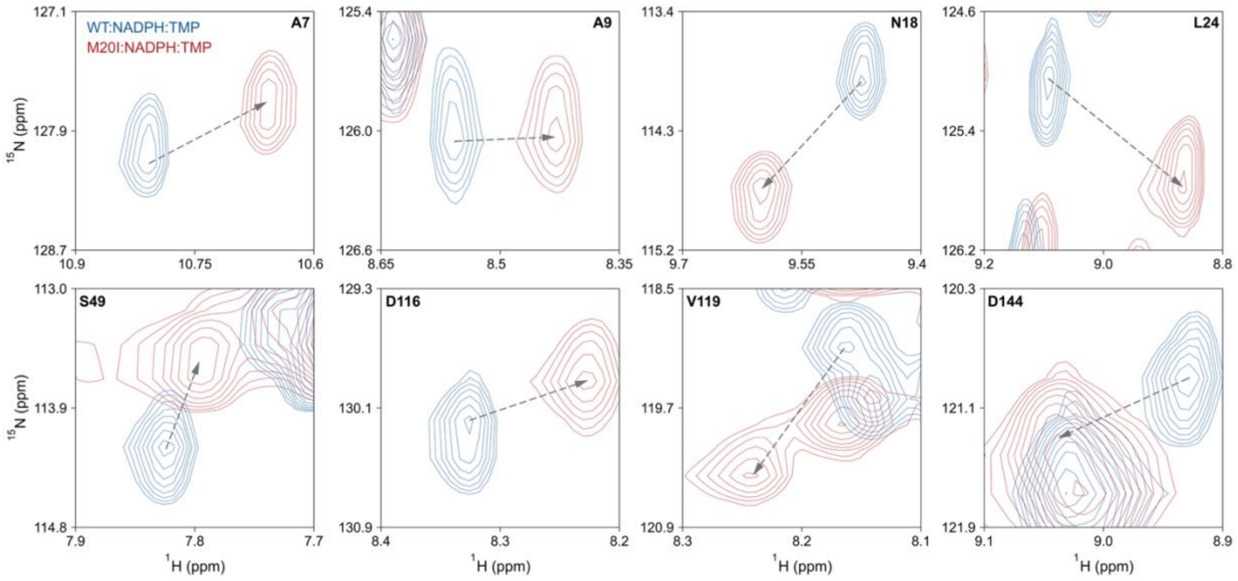


**Supplementary Fig. 20. Localized chemical shift perturbations induced by the M20I mutation in DHFR.** Expanded regions of the 2D $^{1}$H-$^{15}$N HSQC spectra comparing the WT (blue) and M20I (red) DHFR ternary complexes. Dashed arrows track the movement of selected cross-peaks that exhibit significant weighted chemical shift distances (> 0.2 ppm). These highly perturbed resonances map to critical functional domains, including the M20 loop (A7, A9, N18, L24), the substrate-binding pocket (S49), the F-G loop (D116, V119), and the G-H loop (D144).

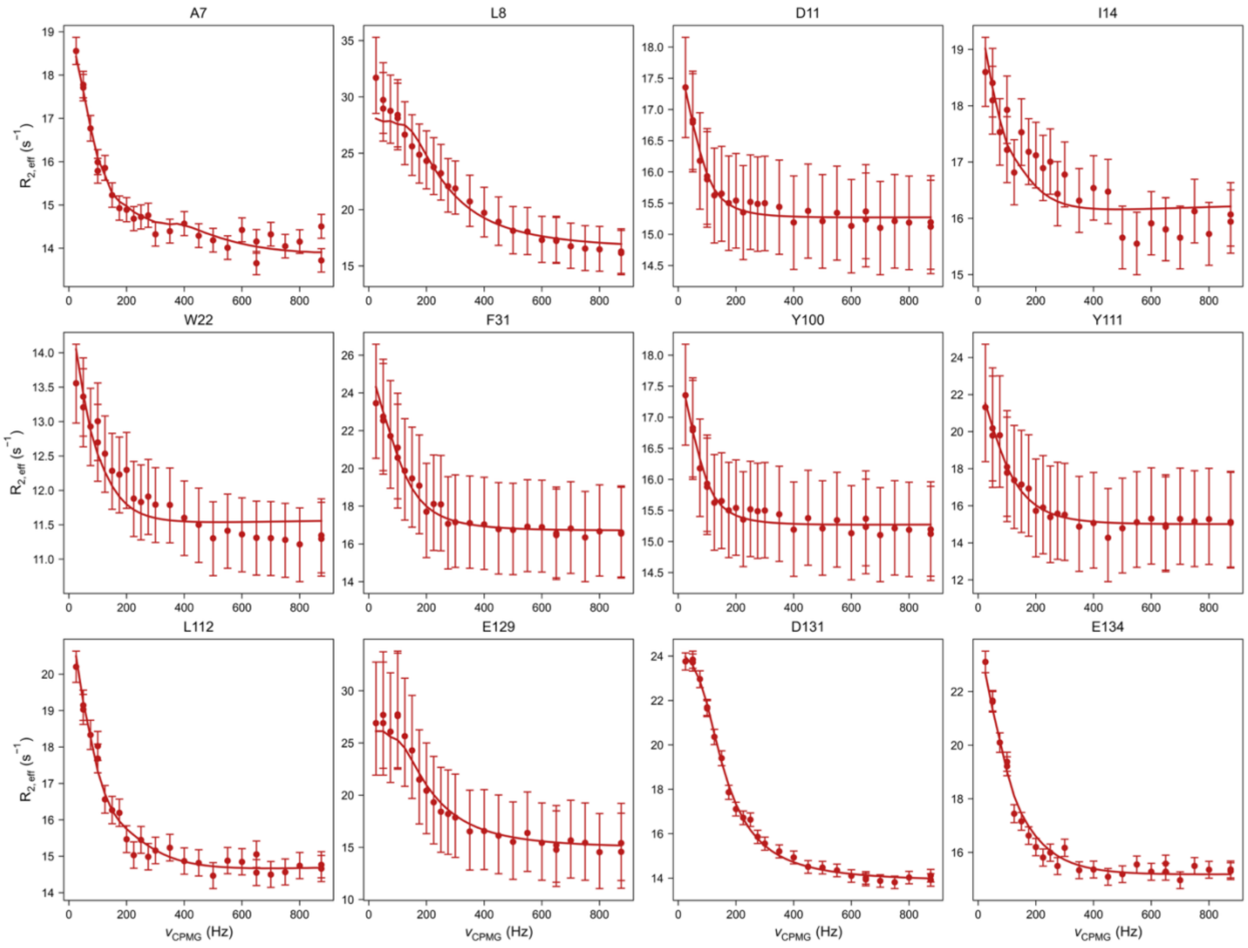


**Supplementary Fig. 21. $^{15}$N CPMG relaxation dispersion curves of M20I residues.** Plots of $R_{2,eff}$ vs. $\nu_{CPMG}$ for selected backbone amides undergoing millisecond-level chemical exchange. These residues map to the M20 loop (A7, L8, D11, I14, W22), the substrate binding pocket (F31), residues interacting with the M20 loop (Y100, Y111, L112), and the F-G loop (E129, D131, E134). Solid lines denote global fits applied to the dispersion data, driven by residues A7, L112, E129, and D131, to extract overarching exchange parameters. Error bars reflect uncertainties derived from PeakFit and ChemEx analysis. The reduced $\chi^2$ for the global fit is 0.48.

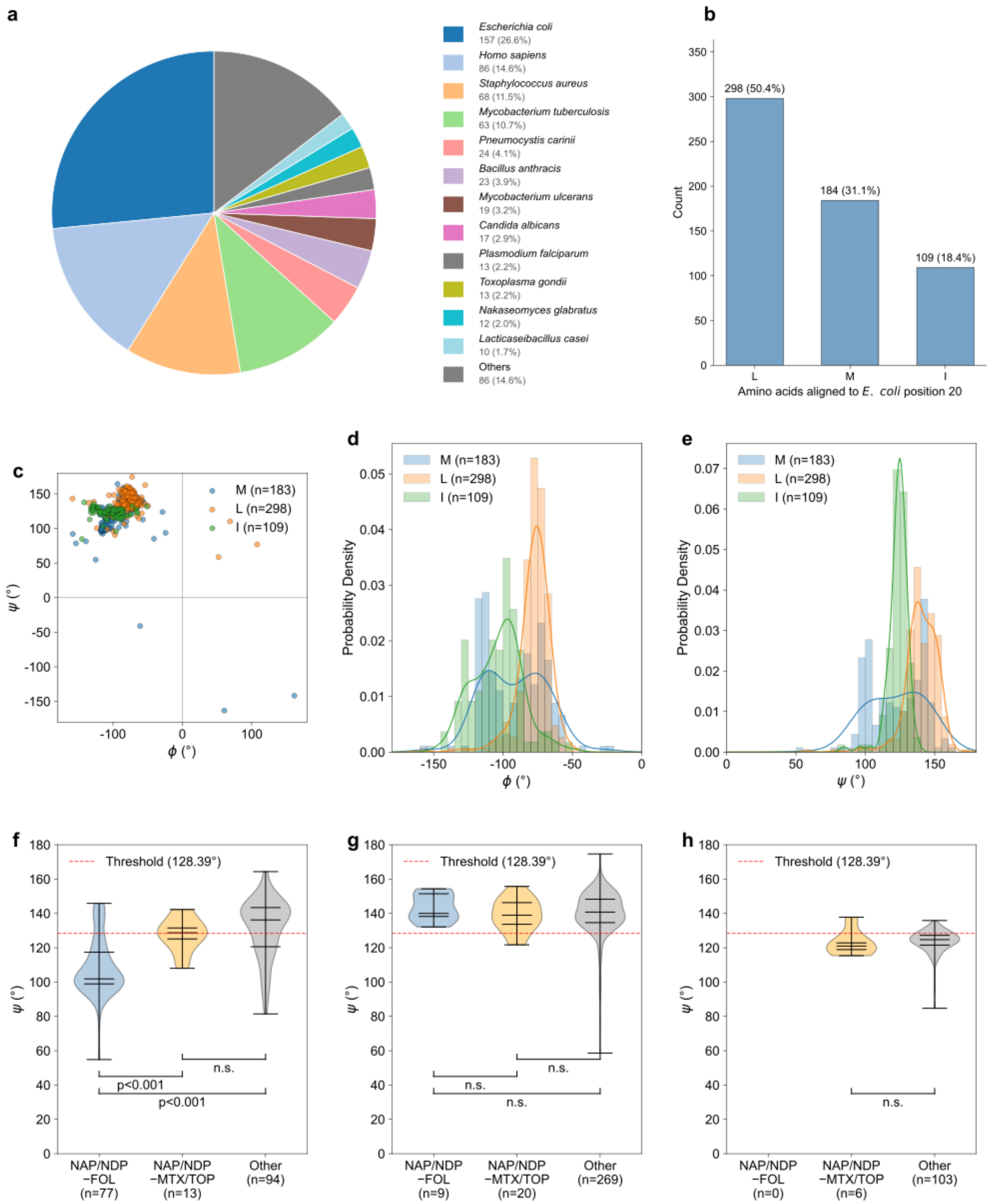


**Supplementary Fig. 22. Meta-analysis of DHFR position 20 conformational plasticity across the PDB.** Analyses utilize all available DHFR structures (n = 591) with sequences aligned to *E. coli* DHFR (UniProt: P0ABQ4). **a, b**, Distribution of source species (**a**) and amino acid composition (**b**) at the aligned position 20. In **a**, species with fewer than 10 available structures are grouped as “Others”. **c**, Ramachandran plot (ϕ vs. ψ) for structures containing methionine (M), leucine (L), or isoleucine (I) at position 20. **d, e**, Probability density functions for the ϕ (**d**) and ψ (**e**) dihedral angles, highlighting the broad conformational sampling of native M compared to the locally rigid distribution of I. **f–h**, Violin plots detailing the ψ angle distributions for M (**f**), L (**g**), and I (**h**) across distinct ligand environments: NAP/NDP ($NADP^+$/NADPH) paired folate (FOL), methotrexate or trimethoprim (MTX/TOP), or other combinations. The red dashed line (ψ = 128.39°) denotes the boundary between low- and high-ψ M subpopulations, defined by the intersection of a two-Gaussian mixture model fitted to the ψ distribution of M. This reference line illustrates how native M dynamically shifts its conformation to accommodate different ligands, whereas I remains sterically restricted. Inside the violins, black horizontal bars denote the 25th, 50th, and 75th percentiles. *p*-values were determined via two-sided Mann–Whitney U tests (n.s. means not significant).

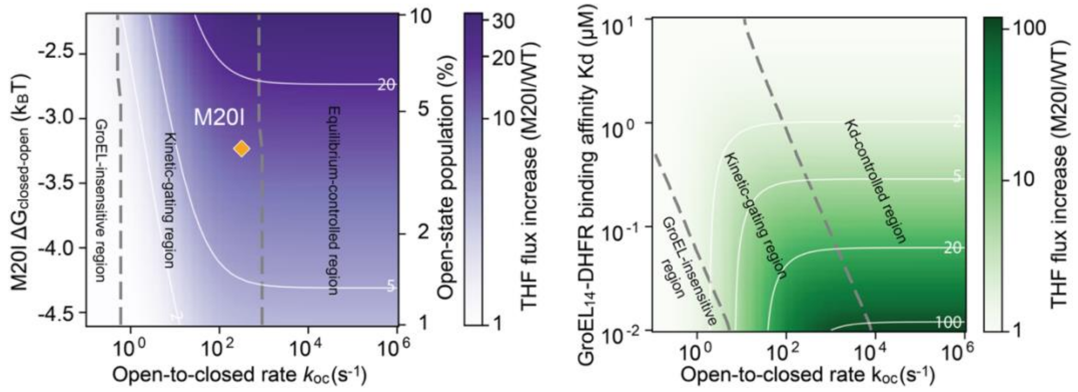


**Supplementary Fig. 23. Kinetic modeling of GroEL/S-mediated THF metabolic flux enhancement. (Left)** Predicted THF flux increase (M20I/WT) plotted against the open-to-closed transition rate ($k_{oc}$) and the free-energy difference between the closed and open states of M20I ($\Delta G_{closed\text{-}open}$). The model defines GroEL-insensitive, kinetic-gating, and equilibrium-controlled regimes. The orange diamond denotes the experimental coordinates for M20I, as determined via NMR CPMG measurements. **(Right)** Predicted flux enhancement mapped against $k_{oc}$ and the $GroEL_{14}$–DHFR binding affinity ($K_d$). This landscape highlights that kinetics predominantly determines the THF flux increase across a wide range of $GroEL_{14}$–DHFR interaction strength.

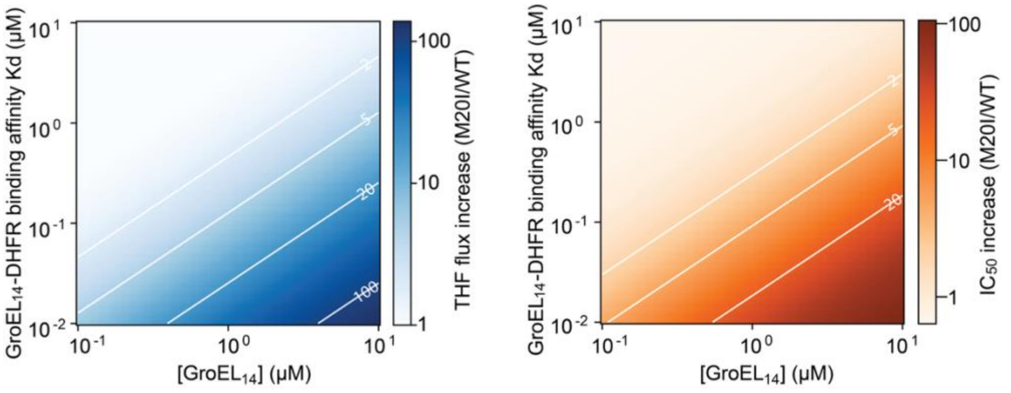


**Supplementary Fig. 24. Dependence of M20I functional rescue on $GroEL_{14}$ concentration and binding affinity. (Left)** Predicted THF metabolic flux increase of the M20I mutant relative to WT plotted as a function of the $GroEL_{14}$ concentration and the $GroEL_{14}$–DHFR binding affinity ($K_d$). **(Right)** Predicted $IC_{50}$ increase of M20I relative to WT mapped across the identical $K_d$ and $GroEL_{14}$ concentration parameter space. Both predicted landscapes illustrate that functional and phenotypic rescue scales synergistically with high chaperone availability and strong interaction strength.

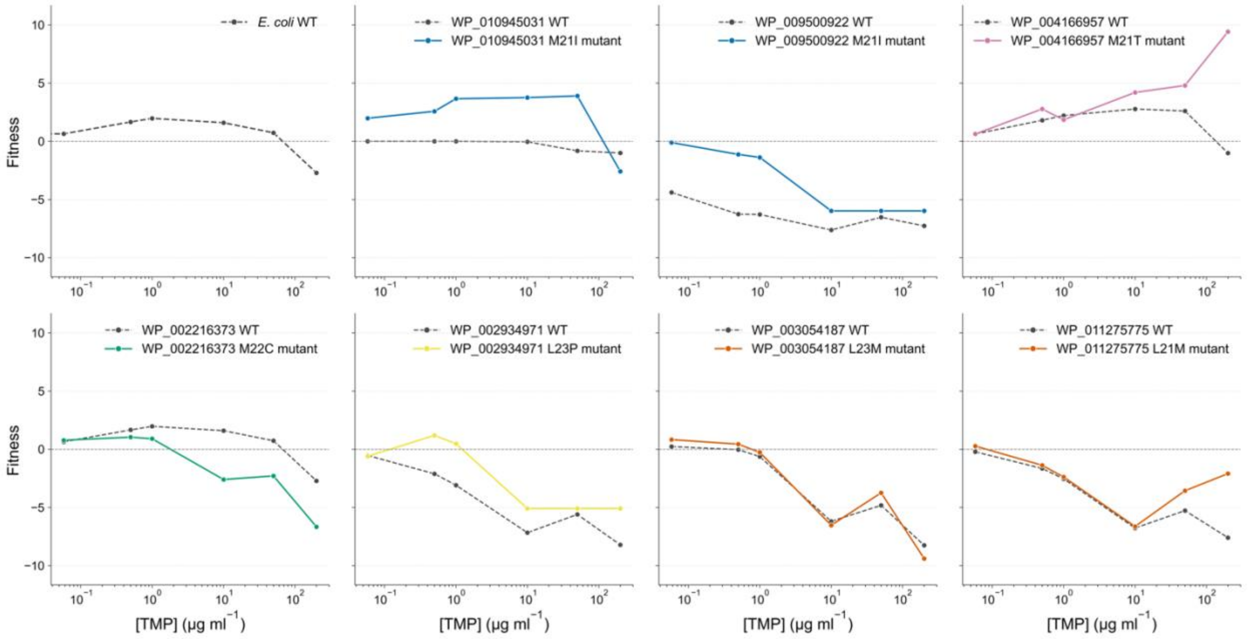


**Supplementary Fig. 25. TMP-response curves of all identified position 20 single-substitution mutants.** Fitness trajectories across varying TMP concentrations are displayed for all seven single-mutant variants identified in the dataset. Each plotted variant only contains a single amino acid substitution at a site corresponding to reference position 20 in *E. coli* DHFR. Fitness was defined as the dimensionless log2-transformed fold change in normalized barcode counts at each drug concentration relative to the no-drug baseline. Solid colored lines indicate the mutants, while dashed black lines represent their corresponding unmutated wild-type homologs. Notably, across all identified variants, only the M-to-I substitutions exhibit enhanced fitness relative to their wild-type counterparts. The mutation labels provided in each panel (e.g., M21I, L23P) denote the exact amino acid position within that specific homolog's sequence. The *E. coli* WT fitness trajectory is shown in the top-left panel as a baseline reference.

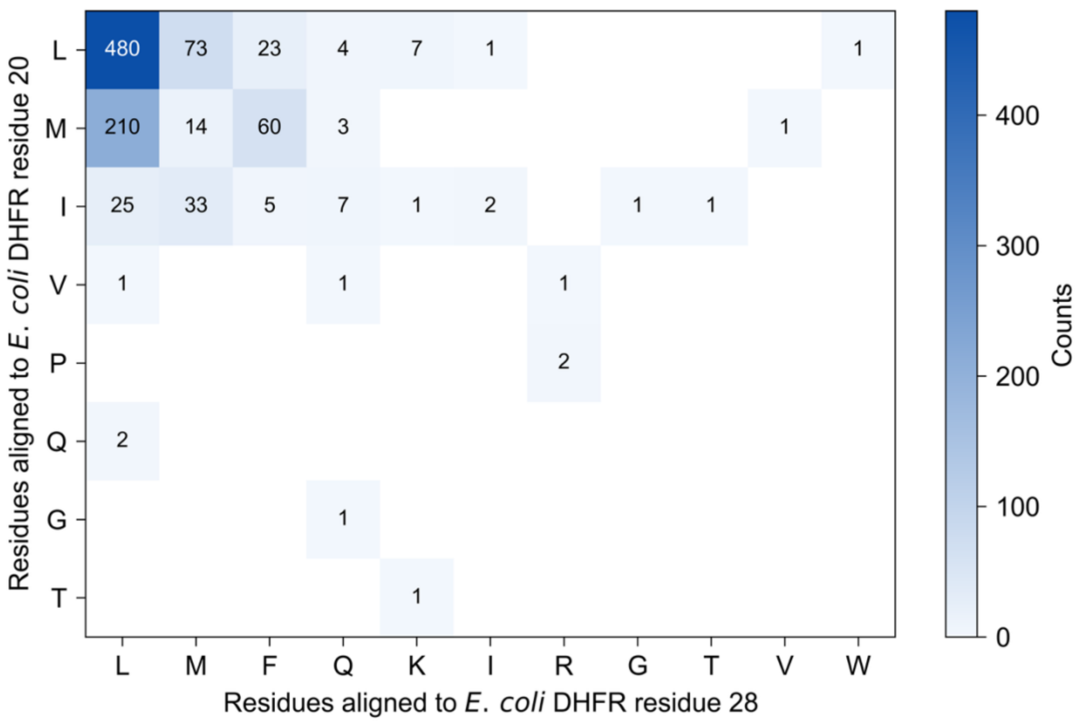


**Supplementary Fig. 26. Distribution of homolog residue pairs aligned to *E. coli* DHFR positions 20 and 28.** DHFR homologs from the lib15/Codon 1 library were categorized according to their amino acids at reference positions 20 and 28 (aligned to *E. coli* DHFR, UniProt: P0ABQ4). The heatmap displays the absolute count of homologs for each observed residue pair.

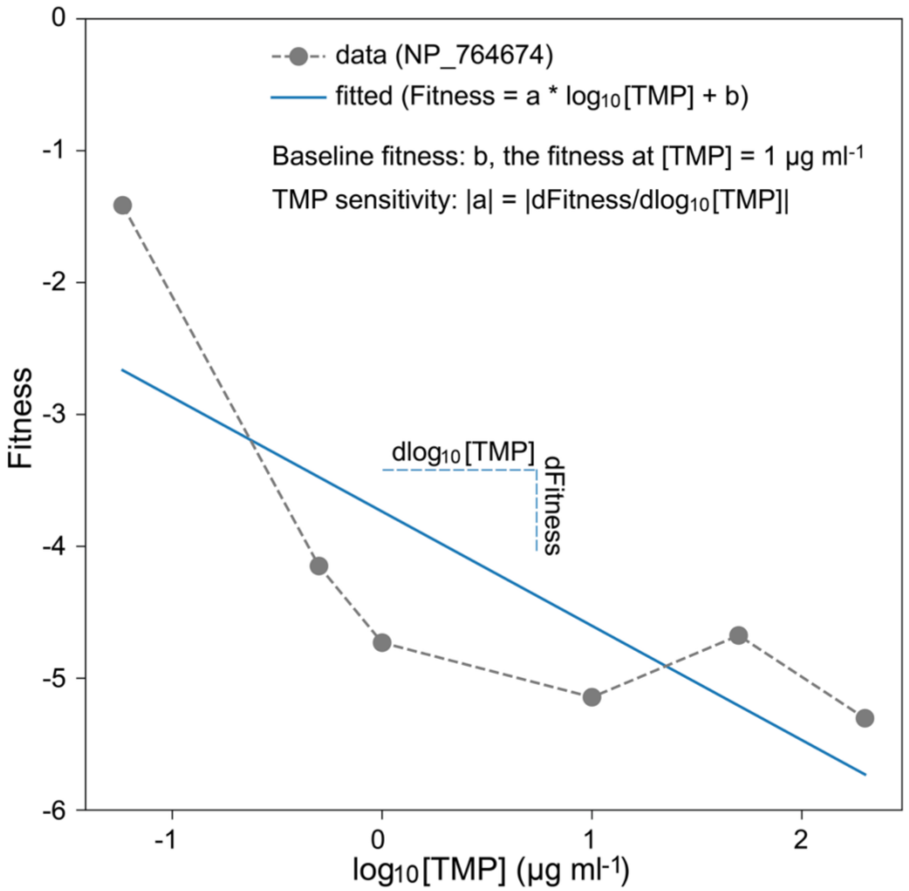


**Supplementary Fig. 27. Schematic representation of TMP-response parameters derived from homolog fitness analyses.** Grey points connected by a dashed line show the experimental fitness values for a representative homolog (NP_764674) across varying TMP concentrations. The solid blue line indicates the linear fitting. The intercept $b$ was used as the fitted baseline fitness, corresponding to the estimated fitness at 1 μg ml$^{-1}$ TMP. The absolute value of slope, $|a|$, was used as the TMP sensitivity, corresponding to the magnitude of fitness change per unit increase in $\log_{10}$[TMP].

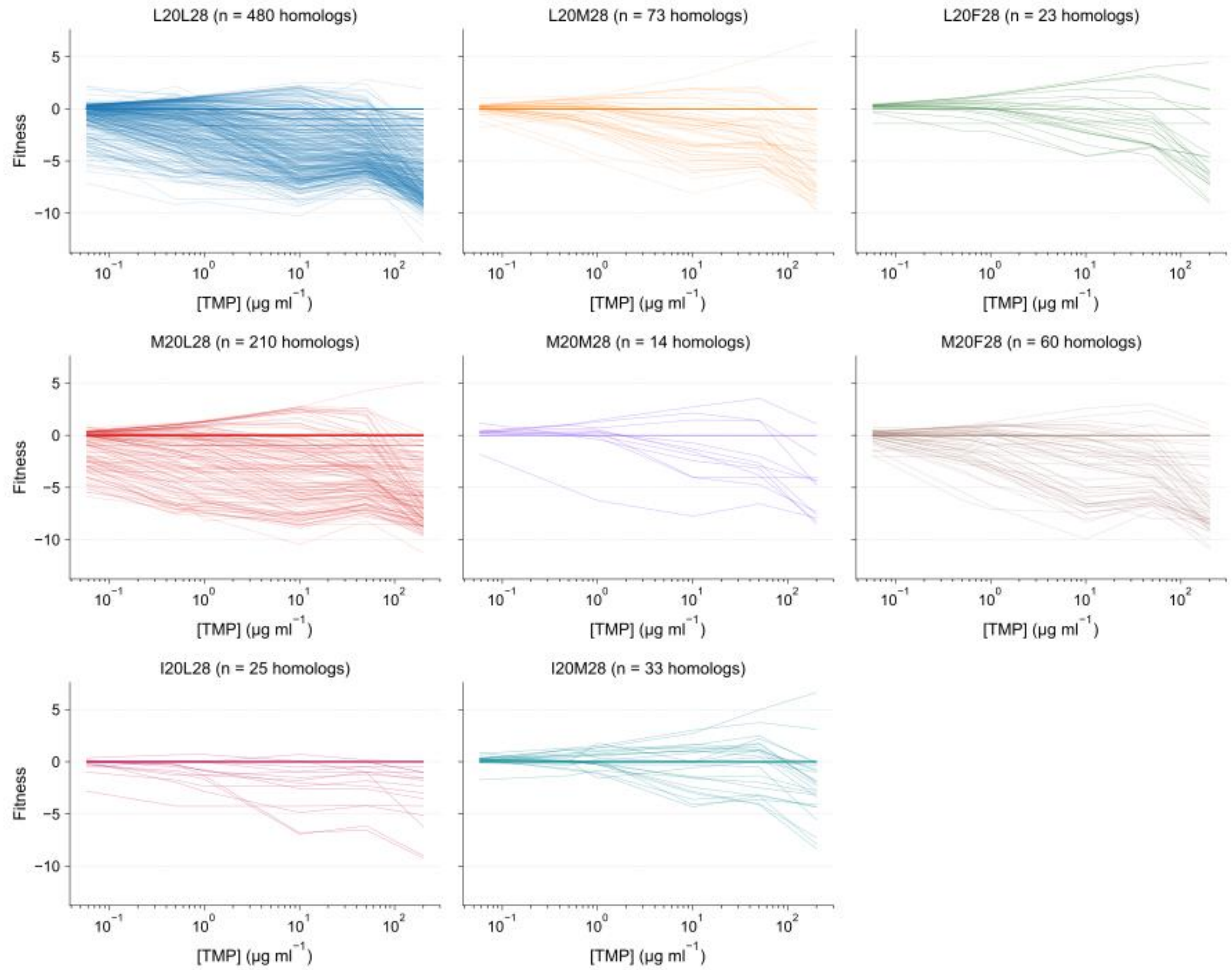


**Supplementary Fig. 28. TMP-response curves of DHFR homologs grouped by position 20 and 28 residue pairs.** Each line depicts the fitness trajectory of an individual homolog across varying TMP concentrations. Only residue-pair groups containing a minimum of 10 homologs are displayed.

**Supplementary Table 1. Summary of DHFR promoter and coding sequence (CDS) mutations identified via directed evolution.**

**Notes:** Trajectories 1–4 spanned approximately 60 days across 80 passages, whereas trajectories 5–16 spanned approximately 45 days across 60 passages. All trajectories were Sanger-sequenced at passage 60. For trajectories 1–4, an additional sequencing was performed at passage 80; mutations unique to this later time point, if any, are explicitly annotated.

| | **GroEL/S Overexpression** | | **Control** | |
|---|---|---|---|---|
| **Trajectory index** | **Promoter** | **CDS** | **Promoter** | **CDS** |
| **1** | none | W30R | none | none |
| **2** | T-15C | M20I @ Passage 60; M20I, L24R @ Passage 80 | T-29C | none |
| **3** | none | P21L @ Passage 60; I5F @ Passage 80 | none | none |
| **4** | T-35C | none | none | none |
| **5** | none | D27E, L28R | none | P21Q |
| **6** | none | none | none | L28R |
| **7** | none | P21L | none | none |
| **8** | none | none | none | D27E |
| **9** | T-5C | none | none | none |
| **10** | T-5C | P21L | T-29C | D27E |
| **11** | T-2C | none | none | A26T, D27E |
| **12** | T-5C | D27E | none | none |
| **13** | T-35C | none | A-9G | W30G, I94L |
| **14** | none | P21L | none | none |
| **15** | none | none | T-5C | D27E, F153C |
| **16** | A-4G | none | none | none |

**Supplementary Table 2. Kinetic model parameters.**

| Notation | Definition | Value in M20I model | Value in WT model | Comment |
|---|---|---|---|---|
| $[\mathbf{DHFR}]_A$ | Concentration of active DHFR | NA | NA | |
| $[\mathbf{DHFR}]_I$ | Concentration of inhibited DHFR | NA | NA | |
| $[\mathbf{DHFR}]_I^c$ | Concentration of inhibited DHFR in closed state | NA | NA | |
| $[\mathbf{DHFR}]_I^o$ | Concentration of inhibited DHFR in open state | NA | NA | |
| $[\mathbf{GroEL_{14}}:\mathbf{DHFR}$ | Concentration of $GroEL_{14}$-DHFR complex | NA | NA | |
| $[\mathbf{GroEL_{14}}]$ | Concentration of $GroEL_{14}$ | 2.2 μM | NA | 1 |
| $[\mathbf{DHFR}]_{total}$ | Total concentration of DHFR in cytoplasm | 0.4 μM | 0.4 μM | 2 |
| $[\mathbf{TMP}]$ | Concentration of TMP | 88.57 μM | 88.57 μM | 3 |
| $k_{disso}$ | Dissociation rate constant of TMP separating from DHFR | 0.058 $s^{-1}$ | 0.092 $s^{-1}$ | 4 |
| $k_{asso}$ | Association rate constant of TMP binding to DHFR | 20 $\mu M^{-1}$ $s^{-1}$ | 20 $\mu M^{-1}$ $s^{-1}$ | 4 |
| $k_G$ | Reaction rate constant of GroEL/S refolding process | 1/8 $s^{-1}$ | NA | 5 |
| $k_{on}$ | Association rate constant of $GroEL_{14}$ and DHFR | $3.18\times10^9$ $M^{-1}$ $s^{-1}$ | NA | 6 |
| $k_{off}$ | Dissociation rate constant of $GroEL_{14}$ and DHFR | Varied with $K_d$ | NA | |
| $\rho^c$ | Population of inhibited DHFR in closed state | NA | NA | |
| $\rho^o$ | Population of inhibited DHFR in open state | NA | NA | |
| $k_{co}$ | Closed-to-open rate constant | Varied | NA | |
| $k_{oc}$ | Open-to-closed rate constant | Varied | NA | |
| $K_d$ | Binding affinity of $GroEL_{14}$ and DHFR | Varied from 0.01 μM to 10 μM | NA | |
| $\Delta G_{closed-open}$ | Free energy difference between closed-state and open-state | Varied with $k_{co}$ and $k_{oc}$ | NA | |
| $k_{cat}$ | Catalytic constant of DHFR enzymatic reaction | 6.8 $s^{-1}$ | 5.3 $s^{-1}$ | 7 |
| $K_M$ | Michaelis constant of DHFR enzymatic reaction | 1.9 μM | 2.2 μM | 7 |
| $[\mathbf{THF}]$ | Concentration of tetrahydrofolate (THF) | NA | NA | |

| | | | | |
|---|---|---|---|---|
| **[DHF]** | Concentration of dihydrofolate (DHF) | 10 μM | 10 μM | 8 |
| $\boldsymbol{\eta}$ | Viscosity of *E. coli* cytoplasm | 3 mPa s | 3 mPa s | 9 |
| $\boldsymbol{r_{GroEL_{14}}}$ | Radius of gyration of $GroEL_{14}$ | 6.6 nm | NA | 10 |
| $\boldsymbol{r_{DHFR}}$ | Radius of gyration of DHFR | 2 nm | NA | 11 |
| $\boldsymbol{T}$ | Temperature | 310 K | 310 K | |

Comments:

1. GroEL protomer abundance is obtained from PaxDb: Protein Abundance Database v6.0, *E. coli* str. K-12 substr. MG1655 - Whole organism (Integrated) dataset, with value as 6886 parts per million (ppm). Given *E. coli* contains $2.36 \times 10^6$ proteins per cell[1], there are around 16,251 GroEL protomers in each cell. Since *E. coli* cell volume is 0.86 $\mu m^3$ [1], *in vivo* GroEL protomer concentration is around 31.4 μM (i.e., $GroEL_{14}$ ~ 2.2 μM).
2. *In vivo* DHFR concentration is calculated in the same way as that of GroEL. DHFR abundance from PaxDb is 88.5 ppm, which converts to cellular concentration as 0.4 μM.
3. *In vivo* TMP concentration is obtained from Manna *et al.* (2021)[2], where *E. coli* BW25113 is treated with 5 μM TMP and harvested after 1 hour incubation. TMP concentration of 1 μM was used for numerical calculation in fig. 5e to mimic the condition used in fig. 3g.
4. Association rate constant between TMP and DHFR is obtained from Cayley *et al.* (1981)[3] with and dissociation rate constant calculated to reproduce measured $K_i$ in fig. 2b.
5. Previous work places the ATP-driven GroEL/GroES reaction cycle on the timescale of several seconds, often approximated as ~8 s [4]; accordingly, we modeled the productive GroEL/S step with reaction rate constant as 1/8 $s^{-1}$.
6. The binding between $GroEL_{14}$ and its substrate proteins is close to diffusion-limited[5–7], therefore, we use diffusion rate constant $k_{diff}$ as a proxy for the association rate constant of $GroEL_{14}$ and DHFR. Based on Smoluchowski equation,

$$k_{on} = k_{diff} = 4\pi\left(D_{GroEL_{14}} + D_{DHFR}\right)\left(r_{GroEL_{14}} + r_{DHFR}\right)$$

where

$$D = \frac{k_B T}{6\pi\eta r}$$

7. Measured by this work.
8. Since a canonical basal intracellular DHF concentration in *E. coli* is not well established, DHF was assumed to be 10 μM as an order-of-magnitude estimate for the modeling analysis. As DHF concentration was treated as a constant and the same value was used in both the WT and M20I models, this assumption does not affect the relative comparison between the two conditions.
9. The viscosity of the E. coli cytoplasm was based on a reported cytoplasmic microviscosity measured in live *E. coli* cells[8].
10. A radius of gyration of 2 nm was used for DHFR, consistent with the size expected for a compact folded protein of approximately 18 kDa and its structure.
11. A radius of gyration of 6.6 nm was used for $GroEL_{14}$, as reported by a solution X-ray scattering measurement for native *E. coli* $GroEL_{14}$[9].

**Supplementary Table 3. Experimentally characterized GroEL-recognizable peptide motifs.**

**Notes:** * Peptide designations match the original literature, defaulting to parent protein names where unspecified. ** In the absence of explicit $K_d$ values, relative affinity comparisons are restricted to intra-study data. *** PAxpS90T serves as the reference sequence. Amino acid identities are represented by hyphens (–), while divergent positions are indicated by the single-letter code of the substituted residue.

| Reference | Peptide* | Sequence | Secondary Structure | Binding strength** |
|---|---|---|---|---|
| 10 | N-terminal α-helix of rhodanese | STKWLAESVRAGK | α-helix | 10-1000 µM (estimated) |
| 11 | vsv-C | KLIGVLSSLFRPK | α-helix | n/a |
| 12 | Residues 19–35 from eosinophil cationic protein (ECP) | NPPRCTIAMRAINNYRWR | n/a | n/a |
| 13*** | PAxpS90T (pS residues 67-86) | PLSTEALLKQVDYLIRSKWV | n/a | 53 nM |
|  | Gly5 | -------GG-G-GG------ |  | 370 nM |
|  | Gly4 | -------GG-GG-------- |  | 100 nM |
|  | Gly3 | -------GG-G--------- |  | 36 nM |
|  | Ser5 | -------SS-S-SS------ |  | 196 nM |
|  | Gln4-1 | ----------Q-QQQ----- |  | 136 nM |
|  | Gln4-2 | -----QQQ--Q--------- |  | 96 nM |
|  | Gln5 | -----QQQ--Q----Q---- |  | 180 nM |
|  | Pro2 | ---------P-P-------- |  | 37 nM |
|  | Pro3-1 | ------P--P-P-------- |  | 62 nM |
|  | Pro3-2 | ------P--P-----P---- |  | 72 nM |
|  | Pro4 | ------P--P-P---P---- |  | 109 nM |
|  | HCH 2 | ------R---D--RR----D |  | 50 nM |
|  | HCH 3 | ------RR------R----- |  | 27 nM |
|  | HCH 4 | ------RR------------ |  | 62 nM |

| | | | | |
|---|---|---|---|---|
|  | HCH 5 | ------RR--D--R------ |  | 62 nM |
| 14 | P23 (the N-terminal 23 amino acid signal sequence of pre-b-lactamase) | LRIQHFRVALIPFFAAFSLPVFG | n/a | 10-100 nM |
| 15 | Rhodanese peptides | Two long peptides at about 7-9,000 Da and 11,000 Da | n/a | n/a |
| 16 | Gp31 | AQAGDEEVTESGLIIGKRVQ | β-hairpin | n/a |
| 17 | APA08c (hydrophilic) | CNCK(FITC)APETKWCAESCRAGK | α-helix: 55%; β-strand: 45% | no binding detetcetd |
| | APA09r (hydrophobic) | CNCK(FITC)APETALCWLACSVRAGK | α-helix: 19%; β-strand: 81% | no binding detetcetd |
| | APA09c (hydrophobic) | CNCK(FITC)APETALCWLACSVRAGK | α-helix: 71%; β-strand: 29% | 106 µM |
| | APA14c (hydrophobic) | CNCKAPETALCWLACSVRAGK | α-helix: 57%; β-strand: 27%; others: 16% | bound |
| 18 | Bamph (basic amphiphilic) | PLYKKIIKKLLES | n/a | 0.54 µM |
| | Namph (neutral amphiphilic) | PLYQQIIQQLLNS | | 3.57 µM |
| | Aamph (acidic amphiphilic) | PLYEEIIEELLES | | >10 µM |
| | Ahphil (acidic hydrophilic) | PQYEENNEEQQES | | >10 µM |
| | Bhphil (basic hydrophilic) | PQYKKNNKKQQES | | 3.44 µM |
| 19 | 7 residues of a 17-residue N-terminal tag attached to the mini GroEL | GLVPRGS | Extended conformation | n/a |
| 20 | GroES | KRKEVETK AGGIVLTGSAA | β-hairpin | n/a |
| 21 | AMPH+ | ALYKKIIKKLLESK | α-helix | 5 nM |
| | AMPH- | ALYKKIIKKLLESK | less α-helix | 51 nM |

| | | | | |
|---|---|---|---|---|
| | AMPHR | ALYKKIIKKLLESK | unstructured | 196 nM |
| | NON-AMPH+ | ALYKIKKIKLLESK | α-helix | 62 nM |
| | NON-AMPH- | ALYKIKKIKLLESK | unstructured | 83 nM |
| | NON-AMPHR | ALYKIKKIKLLESK | unstructured | 215 nM |
| 22 | P22 tailspike | DYVKFPGIETLL | n/a | +++ |
| | vsv-C | KLIGVLSSLFRPK | α-helix | +++ |
| | Rho2-L6 | STKWLLESVRAGK | n/a | +++ |
| | AL | YKALSEALKSAK | n/a | +++ |
| | ID (amphipathic variants of Rho2) | YKALAESLKSAK | α-helix | +++ |
| | Rho2 | STKWLAESVRAGK | α-helix | +++ |
| | D-Rho2 (chirality variants of Rho2) | STKWLAESVRAGK | Left-handed α-helix | +++ |
| | Xhat1 | YSTATLSLGHHAVP | β-strand | +++ |
| | Rho2-5P6 | STKWLPAESVRAGK | n/a | +++ |
| | L,D-Rho2 (chirality variants of Rho2) | STKWLAESVRAGK | unable to form α-helix | +++ |
| | KE (non-amphipathic variants of Rho2) | YKLSAEKLSAAK | α-helix | ++ |
| | ES24D | KRKEVETKSAGDIVLTGSAA | n/a | ++ |
| | ES24G | KRKEVETKSAGGIVLTGSAA | n/a | ++ |
| | Rho2-P5 | STKWPAESVRAGK | n/a | ++ |
| | "control" | GPENRGDSCA | n/a | ++ |
| | Rho2-P5G6 | STKWPGESVRAGK | n/a | ++ |
| | CRAPB3 | SKPHVEIRQDGD | β-strand | + |
| | KWK | KWK | n/a | + |
| 23 | Rho (N-terminal α-helix of rhodanese) | STKWLAESVRAGK | α-helix | n/a |

| | | | | |
|---|---|---|---|---|
| 24 | SBP (strongly binding peptide) | FHYEIWIPPHRG | β-hairpin | 2 μM |
| | 1 | SWMTTPWGFLHP | | |
| | 3 | SSPWWLVSFTST | | |
| | 4 | SHSLIWRIPLLH | | |
| | 5 | IYVPWYYAENLP | | |
| | 6 | YNYSWNGVVFVP | | |
| | 7 | AQSTPLMKPQKS | | |
| | 8 | DQTTLQRFLGSH | | |
| | 9 | QTIKPPITVHPS | | |
| | 10 | QYNHILGYLPFQ | | |
| | 11 | IMDPQNSKVTVA | n/a | n/a |
| | 12 | LPIQNAKRSMVS | | |
| | 13 | IMSPWDESFWNY | | |
| | 14 | ASESYVLFPGTR | | |
| | 15 | SNWHGPLSYQLM | | |
| | 16 | ALPLQDTAATLS | | |
| | 17 | QEIYLTPRGPQQ | | |
| | 18 | IDRTQMWRQSDL | | |
| | 19 | INRDHPLHAGQP | | |
| | 20 | HQTPQSLARWSL | | |
| | 21 | HSLRAIQLITGM | | |
| 25 | Peptide 1 in Ref. [24] | SWMTTPWGFLHP | β-hairpin | n/a |
| 26 | SBP-W2DP6V | SDMTTVWGFLHP | 310/α-helix | 17.1 μM |
| 27 | Peptide 1 in Ref. [24] | SWMTTPWGFHLP | β-hairpin | n/a |

**Supplementary Table 4. X-ray crystallography statistics, related to Fig. 4a.**

| | M20I:NADPH:TMP |
|---|---|
| **PDB ID** | XXXX |
| **Wavelength (Å)** | 1.541840 |
| **Resolution range** | 49.53 - 1.898 (1.966 - 1.898) |
| **Space group** | P 21 21 21 |
| **Unit cell** | 34.1561 45.5504 99.0501 90 90 90 |
| **Total reflections** | 86516 (6206) |
| **Unique reflections** | 12754 (1034) |
| **Multiplicity** | 6.8 (5.2) |
| **Completeness (%)** | 98.47 (85.31) |
| **Mean I/sigma(I)** | 4.81 (0.36) |
| **Wilson B-factor** | 20.39 |
| **R-merge** | 0.1756 (0.7688) |
| **R-meas** | 0.19 (0.8551) |
| **R-pim** | 0.07142 (0.3658) |
| **CC1/2** | 0.992 (0.686) |
| **CC*** | 0.998 (0.902) |
| **Reflections used in refinement** | 12580 (1034) |
| **Reflections used for R-free** | 653 (51) |
| **R-work** | 0.1885 (0.2815) |
| **R-free** | 0.2343 (0.3403) |
| **CC (work)** | 0.960 (0.797) |
| **CC (free)** | 0.947 (0.597) |
| **Number of non-hydrogen atoms** | 1520 |
| **Macromolecules** | 1353 |
| **Ligands** | 119 |
| **Solvent** | 92 |
| **Protein residues** | 159 |
| **Nucleic acid bases** | |
| **RMS (bonds)** | 0.021 |
| **RMS (angles)** | 1.74 |
| **Ramachandran favored (%)** | 97.45 |
| **Ramachandran allowed (%)** | 2.55 |
| **Ramachandran outliers (%)** | 0 |
| **Rotamer outliers (%)** | 2.07 |
| **Clashscore** | 8.96 |
| **Average B-factor** | 21.79 |
| **Macromolecules** | 21.55 |
| **Ligands** | 18.84 |
| **Solvent** | 27.76 |

**Supplementary Table 5. Reagent and Resources.**

| Reagent of Resource | Source | Identifier |
|---|---|---|
| Terrific Broth (TB) | BD Difco™ | 243820 |
| LB broth | Millipore® | 1.10285 |
| M9 minimal salts | BD Difco™ | 248510 |
| Glycerol | VWR Chemicals BDH® | BDH1172 |
| D-(+)-Glucose monohydrate | Millipore® | 49159 |
| MOPS | Sigma-Aldrich® | M3183 |
| Citric acid | VWR Chemicals BDH® | BDH4136 |
| Ferric ammonium citrate | VWR AMRESCO® | 0846 |
| Magnesium sulfate ($MgSO_4$) | Sigma-Aldrich® | M2643 |
| Calcium chloride ($CaCl_2$) | VWR Chemicals BDH® | BDH4122 |
| Arabinose | Alfa Aesar | A11921 |
| Trimethoprim | Sigma-Aldrich® | T7883 |
| isopropyl β-D-1-thiogalactopyranoside (IPTG) | RPI | I56000 |
| Kanamycin sulfate | Millipore® | 420311 |
| Streptomycin sulfate salt | Sigma-Aldrich® | S9137 |
| Ampicillin | VWR | 0339 |
| Chloramphenicol | Calbiochem® | 220551 |
| Anhydrotetracycline (aTc) | Takara | 631310 |
| Bicine | Sigma-Aldrich® | B3876 |
| Imidazole | Millipore® | 5710-OP |
| Sodium chloride (NaCl) | VWR Chemicals BDH® | BDH9286 |
| Potassium chloride (KCl) | Sigma-Aldrich® | P3911 |
| Tris base | Sigma-Aldrich® | T4661 |
| Tris-Buffered Saline (TBS) | Corning® | 46-012-CM |
| Phosphate Buffered Saline (PBS) | Corning® | 46-013-CM |
| Benzonase® Nuclease | Millipore® | 70664 |
| cOmplete™, Mini, EDTA-free Protease Inhibitor Cocktail | Roche | 11836170001 |
| Lysozyme | Thermo Scientific™ | 89833 |
| TCEP Hydrochloride | RPI | T26500 |
| 0.5 M EDTA | Invitrogen™ | AM9261 |
| Dithiothreitol (DTT) | RPI | D11000 |
| cOmplete™ His-Tag Purification Resin | Roche | 5893682001 |
| Methotrexate (MTX)-Agarose | Sigma-Aldrich® | M0269 |
| Ammonium sulfate ($(NH_4)_2SO_4$) | Sigma-Aldrich® | 09978 |
| Monopotassium phosphate ($KH_2PO_4$) | G-Biosciences | RC084 |
| Dipotassium phosphate ($K_2HPO_4$) | G-Biosciences | RC081 |
| Potassium tetraborate tetrahydrate ($K_2B_4O_7 \cdot 4H_2O$) | Sigma-Aldrich® | P5754 |
| HEPES sodium salt | Sigma-Aldrich® | H3784 |

| | | |
|---|---|---|
| D-Glucose (U-$^{13}C_6$) | Cambridge Isotope Laboratories | CLM-1396 |
| Ammonium chloride ($^{15}N$) | Cambridge Isotope Laboratories | NLM-467 |
| Ammonium sulfate ($^{15}N_2$) | Cambridge Isotope Laboratories | NLM-713 |
| Deuterium oxide "100%" (D, 99.96%) +0.01 mg/mL DSS | Cambridge Isotope Laboratories | DLM-6DB-10X0.7 |
| MEM Vitamins 100x Solution | Corning® | 25-020-CI |
| Acetone | VWR Chemicals BDH® | BDH1101 |
| L-Malate Dehydrogenase (L-MDH) | Roche | 10127914001 |
| Pyruvate Kinase (PK) | Roche | 10128155001 |
| Phosphoenol-pyruvate (PEP-K) | Roche | 10108294001 |
| NADP | Roche | 10128031001 |
| Oxaloacetic acid | Sigma-Aldrich® | O7753 |
| Magnesium acetate | VWR Chemicals BDH® | BDH4170 |
| bis-ANS | Invitrogen™ | B153 |
| NADPH | Roche | 10621706001 |
| Dihydrofolic acid (DHF) | Sigma-Aldrich® | D7006 |
| Sodium hydroxide (NaOH) | Macron | 7708-10 |
| Dimethyl sulfoxide (DMSO) | Sigma-Aldrich® | 276855 |
| Adenosine 5′-triphosphate disodium salt hydrate (ATP) | Sigma-Aldrich® | A2383 |
| EveryBlot Blocking Buffer | Bio-Rad | 12010020 |
| Urea | Sigma-Aldrich® | RES9692U-A102X |
| Guanidine hydrochloride (GdnHCl) | Sigma-Aldrich® | G3272 |
| DSP | Thermo Scientific™ | A35393 |
| B-PER™ II Bacterial Protein Extraction Reagent | Thermo Scientific™ | 78260 |
| Tween-20 | Sigma-Aldrich® | P9416 |
| Thiamine hydrochloride | Sigma-Aldrich® | T1270 |
| Casamino acids | Teknova | C2000 |
| D-Glucose 6-phosphate sodium salt | Sigma-Aldrich® | G7879 |
| Glucose-6-phosphate Dehydrogenase | Sigma-Aldrich® | G6378 |
| Rabbit anti-DHFR antibody | Pacific Immunology | Customized |
| HRP-linked Goat anti-Rabbit IgG | Bio-Rad | STAR124P |
| HRP-linked Horse Anti-Mouse IgG | Cell Signaling Technology | 7076S |
| Mouse anti-GroEL antibody | abcam | ab82592 |
| Rabbit anti-GroEL antibody | abcam | ab90522 |
| Pierce™ Protein A/G Magnetic Beads | Thermo Scientific™ | 88803 |
| HiLoad 16/600 Superdex 75 pg column | Cytiva | 28989333 |
| HiPrep Q Fast Flow 16/10 column | Cytiva | 28936543 |
| HiLoad 16/600 Superose 6 pg column | Cytiva | 29323952 |

| | | |
|---|---|---|
| Superdex 200 Increase 10/300 GL column | Cytiva | 28990944 |
| XK 16/20 Column | Cytiva | 28988937 |
| Amicon® Stirred Cell | Millipore® | UFSC40001 |
| Low Pressure/Vacuum (LPV) NMR tube | SP Wilmad® | 535-LPV-200M |

**Supplementary Table 6. Plasmids.**

| Plasmid | Vector | Protein encoded | Comments |
|---|---|---|---|
| pET-28a(+)-DHFR-6His | pET-28a(+) | *E. coli* DHFR (WT and single mutants) | Synthesized by GenScript Biotech (Nanjing, China); for protein purification. |
| pET-28a(+)-DHFR | pET-28a(+) | *E. coli* DHFR (WT and M20I) | Synthesized by GenScript Biotech (Nanjing, China) with C-terminal 6×His-tag removed; for protein purification. |
| pOA-GroEL/S | pOA | *E. coli* GroEL (no tag) and *E. coli* GroES (C-terminal 6×His-tag) | Gift from Dr. Shimon Bershtein; for protein purification. |
| pGro7 | N/A | *E. coli* GroEL and *E. coli* GroES | Obtained from Takara Bio (Cat. #3340); for GroEL/S overexpression. |
| pEmpty | N/A | None | Constructed from pGro7; control. |
| pdCas9-bacteria | p15A | Catalytically inactive bacterial Cas9 (S. pyogenes) | pdCas9-bacteria was a gift from Stanley Qi[28] (Addgene plasmid # 44249 ; http://n2t.net/addgene:44249 ; RRID:Addgene_44249). |
| pgRNA-bacteria-groSP | pUC19 | N/A | pgRNA-bacteria was a gift from Stanley Qi[28] (Addgene plasmid # 44251 ; http://n2t.net/addgene:44251 ; RRID:Addgene_44251).<br><br>Customized guide RNA (gRNA): tgaccagagaaatggggatg (5'→3'). |
| pgRNA-bacteria-empty | pUC19 | N/A | pgRNA-bacteria was a gift from Stanley Qi[28] (Addgene plasmid # 44251 ; http://n2t.net/addgene:44251 ; RRID:Addgene_44251).<br><br>Guide RNA (gRNA) was removed. |
| pfolA-GFP | N/A | Green fluorescent protein (GFP) | Obtained from Horizon Discovery (Cat. #PEC3876-202384323) |

**Supplementary Table 7. Primers.**

| Primer | Sequence (5'→3') | Comments |
|---|---|---|
| pEmpty_fwd | taattgccctgcacctcg | Construct pEmpty from pGro7 |
| pEmpty_rvs | tgataactctcctttgagaaagtccg | |
| pred_groSp_fwd | AATGGGGATGgttttagagctagaaatagcaag | Construct pgRNA-bacteria-groSP from pgRNA-bacteria |
| pred_groSp_rev | TCTCTGGTCAactagtattatacctaggactg | |
| empty_gRNA_fwd | gttttagagctagaaatagcaag | Construct pgRNA-bacteria-empty from pgRNA-bacteria |
| empty_gRNA_rev | actagtattatacctaggactg | |
| pfolA_c-2t_GFP_FWD | AAAAAAAATTaTCGCCACTATACG | Introduce c-2t mutation to folA promoter |
| pfolA_c-2t_GFP_REV | ATCGGGAAATCTCAATG | |
| pfolA_g-4a_GFP_FWD | AAAAAATTGTtGCCACTATACG | Introduce g-4a mutation to folA promoter |
| pfolA_g-4a_GFP_REV | TTATCGGGAAATCTCAATG | |
| pfolA_c-5t_GFP_FWD | AAAAATTGTCaCCACTATACGTAAAG | Introduce c-5t mutation to folA promoter |
| pfolA_c-5t_GFP_REV | TTTATCGGGAAATCTCAATG | |
| pfolA_g-9a_GFP_FWD | ATTGTCGCCAtTATACGTAAAG | Introduce g-9a mutation to folA promoter |
| pfolA_g-9a_GFP_REV | TTTTTTTATCGGGAAATCTC | |
| pfolA_c-15t_GFP_FWD | GCCACTATACaTAAAGCGTAAAC | Introduce c-15t mutation to folA promoter |
| pfolA_c-15t_GFP_REV | GACAATTTTTTTATCGGGAAATC | |
| pfolA_c-29t_GFP_FWD | AGCGTAAACCaTCGTCGACTG | Introduce c-29t mutation to folA promoter |
| pfolA_c-29t_GFP_rev | TTACGTATAGTGGCGAC | |
| pfolA_c-35t_GFP_FWD | AACCGTCGTCaACTGGTGCGA | Introduce c-35t mutation to folA promoter |
| pfolA_c-35t_GFP_REV | TACGCTTTACGTATAGTGGC | |

**Supplementary Table 8. Bacterial strains.**

| Strain | Genotype | Plasmid | Comments |
|---|---|---|---|
| *E. coli* BL21(DE3) | / | None | NEB (Cat. #C2527) |
| *E. coli* DH5α | / | None | NEB (Cat. #C2987) |
| *E. coli* BW25113 | / | None | CGSC: 7636 |
| *E. coli* TG1/pOA | *E. coli* TG1 | pOA-GroEL/S | Gift from Dr. Shimon Bershtein |
| JL01 | BW25113 *folA*(M20I) | None | This study |
| JL02 | BW25113 *folA*(L28R) | None | This study |
| JL03 | BW25113 *folA*(W30C) | None | This study |
| JL04 | BW25113 | pdCas9-bacteria; pgRNA-bacteria-groSP | This study |
| JL05 | BW25113 | pdCas9-bacteria; pgRNA-bacteria-empty | This study |
| JL06 | BW25113 *folA*(M20I) | pdCas9-bacteria; pgRNA-bacteria-groSP | This study |
| JL07 | BW25113 *folA*(M20I) | pdCas9-bacteria; pgRNA-bacteria-empty | This study |
| JL08 | BW25113 *folA*(L28R) | pdCas9-bacteria; pgRNA-bacteria-groSP | This study |
| JL09 | BW25113 *folA*(L28R) | pdCas9-bacteria; pgRNA-bacteria-empty | This study |
| JL10 | BW25113 *folA*(W30C) | pdCas9-bacteria; pgRNA-bacteria-groSP | This study |
| JL11 | BW25113 *folA*(W30C) | pdCas9-bacteria; pgRNA-bacteria-empty | This study |
| JL12 | BW25113 | pGro7 | This study |

| | | | |
|---|---|---|---|
| JL13 | BW25113 | pEmpty | This study |
| JL14 | BW25113 *folA*(M20I) | pGro7 | This study |
| JL15 | BW25113 *folA*(M20I) | pEmpty | This study |

### Reference


1. Milo, R. What is the total number of protein molecules per cell volume? A call to rethink some published values. *BioEssays* **35**, 1050–1055 (2013).
2. Manna, M. S. *et al.* A trimethoprim derivative impedes antibiotic resistance evolution. *Nat. Commun.* **12**, 2949 (2021).
3. Cayley, P. J., Dunn, S. M. J. & King, R. W. Kinetics of substrate, coenzyme, and inhibitor binding to Escherichia coli dihydrofolate reductase. *Biochemistry* **20**, 874–879 (1981).
4. Ueno, T., Taguchi, H., Tadakuma, H., Yoshida, M. & Funatsu, T. GroEL Mediates Protein Folding with a Two Successive Timer Mechanism. *Mol. Cell* **14**, 423–434 (2004).
5. Perrett, S., Zahn, R., Stenberg, G. & Fersht, A. R. Importance of electrostatic interactions in the rapid binding of polypeptides to GroEL1. *J. Mol. Biol.* **269**, 892–901 (1997).
6. Sparrer, H., Lilie, H. & Buchner, J. Dynamics of the GroEL – Protein Complex: Effects of Nucleotides and Folding Mutants. *J. Mol. Biol.* **258**, 74–87 (1996).
7. Gray, T. E. & Fersht, A. R. Refolding of Barnase in the Presence of GroE. *J. Mol. Biol.* **232**, 1197–1207 (1993).
8. Chen, E., Esquerra, R. M., Meléndez, P. A., Chandrasekaran, S. S. & Kliger, D. S. Microviscosity in E. coli Cells from Time-Resolved Linear Dichroism Measurements. *J. Phys. Chem. B* **122**, 11381–11389 (2018).
9. Igarashi, Y. *et al.* Solution X-ray scattering study on the chaperonin GroEL from *Escherichia coli*. *Biophys. Chem.* **53**, 259–266 (1995).
10. Landry, S. J. & Gierasch, L. M. The chaperonin GroEL binds a polypeptide in an .alpha.-helical conformation. *Biochemistry* **30**, 7359–7362 (1991).
11. Landry, S. J., Jordan, R., McMacken, R. & Gierasch, L. M. Different conformations for the same polypeptide bound to chaperones DnaK and GroEL. *Nature* **355**, 455–457 (1992).

12. Rosenberg, H. F., Ackerman, S. J. & Tenen, D. G. Characterization of a distinct binding site for the prokaryotic chaperone, GroEL, on a human granulocyte ribonuclease. *J. Biol. Chem.* **268**, 4499–4503 (1993).

13. Dessauer, C. W. & Bartlett, S. G. Identification of a chaperonin binding site in a chloroplast precursor protein. *J. Biol. Chem.* **269**, 19766–19776 (1994).

14. Zahn, R. *et al.* Thermodynamic Partitioning Model for Hydrophobic Binding of Polypeptides by GroEL: I. GroEL Recognizes the Signal Sequences of β-lactamase Precursor. *J. Mol. Biol.* **242**, 150–184 (1994).

15. Hlodan, R., Tempst, P. & Hartl, F. U. Binding of defined regions of a polypeptide to GroEL and its implications for chaperonin-mediated protein folding. *Nat. Struct. Biol.* **2**, 587–595 (1995).

16. Landry, S. J., Taher, A., Georgopoulos, C. & van der Vies, S. M. Interplay of structure and disorder in cochaperonin mobile loops. *Proc. Natl. Acad. Sci.* **93**, 11622–11627 (1996).

17. Brazil, B. T. *et al.* Model Peptide Studies Demonstrate That Amphipathic Secondary Structures Can Be Recognized by the Chaperonin GroEL (cpn60)*. *J. Biol. Chem.* **272**, 5105–5111 (1997).

18. Hutchinson, J. P., Oldham, T. C., El-Thaher, T. S. H. & Miller, A. D. Electrostatic as well as hydrophobic interactions are important for the association of Cpn60 (groEL) with peptides. *J. Chem. Soc. Perkin Trans. 2* 279–288 (1997) doi:10.1039/a604880c.

19. Buckle, A. M., Zahn, R. & Fersht, A. R. A structural model for GroEL–polypeptide recognition. *Proc. Natl. Acad. Sci.* **94**, 3571–3575 (1997).

20. Xu, Z., Horwich, A. L. & Sigler, P. B. The crystal structure of the asymmetric GroEL–GroES–(ADP)7 chaperonin complex. *Nature* **388**, 741–750 (1997).

21. Preuss, M., Hutchinson, J. P. & Miller, A. D. Secondary Structure Forming Propensity Coupled with Amphiphilicity Is an Optimal Motif in a Peptide or Protein for Association with Chaperonin 60 (GroEL). *Biochemistry* **38**, 10272–10286 (1999).

22. Wang, Z., Feng, H., Landry, S. J., Maxwell, J. & Gierasch, L. M. Basis of Substrate Binding by the Chaperonin GroEL. *Biochemistry* **38**, 12537–12546 (1999).

23. Kobayashi, N., Freund, S. M. V., Chatellier, J., Zahn, R. & Fersht, A. R. NMR analysis of the binding of a rhodanese peptide to a minichaperone in solution1. *J. Mol. Biol.* **292**, 181–190 (1999).

24. Chen, L. & Sigler, P. B. The Crystal Structure of a GroEL/Peptide Complex: Plasticity as a Basis for Substrate Diversity. *Cell* **99**, 757–768 (1999).

25. Wang, J. & Chen, L. Domain Motions in GroEL upon Binding of an Oligopeptide. *J. Mol. Biol.* **334**, 489–499 (2003).

26. Li, Y., Gao, X. & Chen, L. GroEL Recognizes an Amphipathic Helix and Binds to the Hydrophobic Side*. *J. Biol. Chem.* **284**, 4324–4331 (2009).

27. Izert-Nowakowska, M. A. *et al.* Targeted protein degradation in Escherichia coli using CLIPPERs. *EMBO Rep.* **26**, 3994–4016 (2025).

28. Qi, L. S. *et al.* Repurposing CRISPR as an RNA-Guided Platform for Sequence-Specific Control of Gene Expression. *Cell* **152**, 1173–1183 (2013).
